# Cytoplasmic localization of PUS7 facilitates a pseudouridine-dependent enhancement of cellular stress tolerance

**DOI:** 10.1101/2025.09.22.675887

**Authors:** Minli Ruan, Sean M. Engels, Matthew R. Burroughs, Xiaoyan Li, Rosella Stower, Talia Tzadikario, Connor Powell, Dylan Bloch, Oleksandra Fanari, Stuart Akeson, Daniel E. Eyler, Chase A. Weidmann, Sara Rouhanifard, Miten Jain, Lydia M. Contreras, Kristin S. Koutmou

**Affiliations:** University of Michigan, Department of Biological Chemistry, Ann Arbor, MI 48109; University of Texas, McKetta Department of Chemical Engineering, Austin, TX 78712; University of Michigan, Department of Chemistry, Ann Arbor, MI 48109; Northeastern University, Department of Bioengineering, Boston, MA 02120

## Abstract

Pseudouridine (Ψ) is an abundant post-transcriptional modification found across all classes of RNA. It has been widely speculated that Ψ inclusion in mRNAs might provide an avenue for cells to control gene expression post-transcriptionally. Here we demonstrate that one of the principal mRNA pseudouridylating enzymes, pseudouridine synthase 7 (PUS7), exhibits a stress-induced accumulation in the cytoplasm of yeast and human epithelial lung cells. Stress-induced and cytoplasmic localization of PUS7 promote Ψ-incorporation into hundreds of mRNA targets. Furthermore, engineered PUS7 cytoplasmic localization increases cellular fitness under ROS and divalent metal ion stress. Consistent with this, transcripts modified upon PUS7 cytoplasmic localization are enriched within mRNAs encoding proteins involved in divalent metal metabolism and ROS stress pathways. In contrast, tRNA sites modified by PUS7 (Ψ13 and Ψ35 are unperturbed). Quantitative proteomics reveal a reshaping of the proteome upon PUS7 relocalization under stress, with proteins involved in metal and ROS homeostasis being particularly sensitive to PUS7 localization. Collectively, our data demonstrate that PUS7 localization alters mRNA pseudouridylation patterns to modulate protein production and enhance cellular fitness.

## INTRODUCTION

Post-transcriptional chemical modifications are essential modulators of RNA structure and function in all organisms. The significance of modifications in noncoding RNA biology has been acknowledged since the 1960s. However, the idea that messenger RNA (mRNA) modifications can regulate gene expression has only recently gained traction. Breakthroughs in deep-sequencing technology enabled the transcriptome-wide mapping of chemical modifications to mRNAs^1^. Foundational work has led to the identification of three modifications -N6-methyladenosine (m^6^A), inosine (I), and pseudouridine (Ψ)

– that are widely incorporated into eukaryotic mRNAs and fluctuate in response to cellular and environmental conditions^2,3^. Changes in m^6^A and inosine modification are associated with alterations in mRNA stability, maturation, and translation – ultimately impacting key cellular processes, including development and innate immunity. The biological significance (if any) of Ψ and the remaining > 10 modifications mapped to mRNAs remains an open question^2–7^. Here, we demonstrate that altering the distribution of Ψ can influence the proteome to impact the cellular response to reactive oxygen species (ROS) and metal ion stress.

Pseudouridine is the C5-glycoside isomer of uridine and is incorporated into mRNAs at levels nearly equivalent to m^6^A, the most abundant mRNA modification^2,8^(**Fig. 1a**). Ψ has remained under-studied relative to m^6^A and inosine, largely because robust technologies to map Ψ within mRNA sequences are only now emerging^2,8,9^. Additionally, in contrast to m^6^A and inosine, the enzymes that catalyze Ψ formation in mRNAs modify all three classes of RNA central to the protein synthesis machinery: mRNA, tRNA, and rRNA^10,11^. This has limited the utility of traditional enzyme knock-out/knock-down studies for assessing Ψ function due to the likelihood of pleiotropic effects. As such, our understanding of how Ψ contributes to mRNA biology lags behind that of the other two most widespread modifications^5^. Current insights into the possible molecular-level functions of Ψ in mRNA come from studies that use a combination of *in vitro* and cellular reporter approaches. These investigations suggest that Ψ likely has a broad range of impacts, as its inclusion in mRNAs can alter splicing, translation, and RNA-protein binding^12–16,17^.

**Fig. 1.**
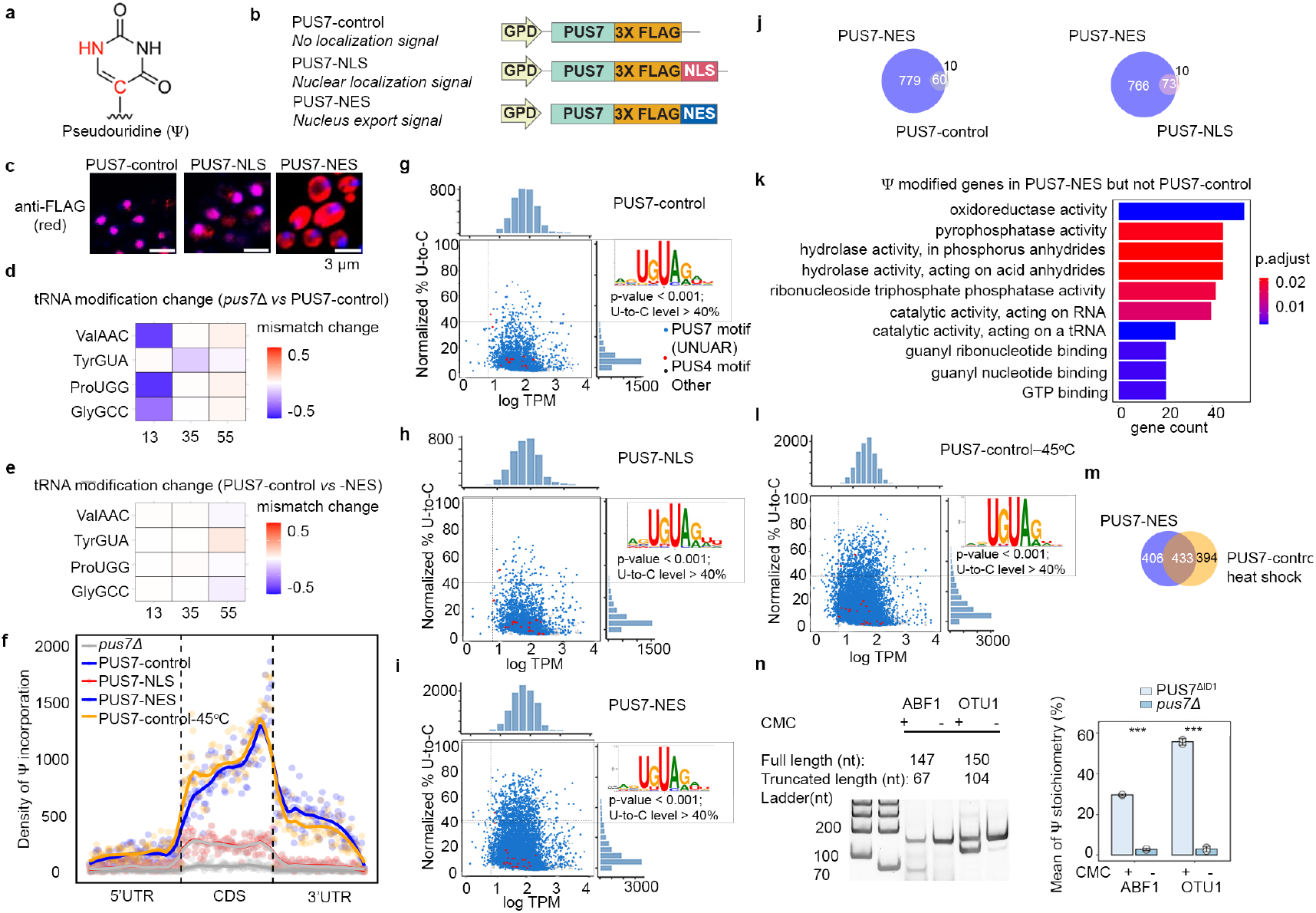
mRNA Ψ sites increase significantly upon PUS7 cytoplasmic localization. **a**, The chemical structure of pseudouridine. **b**, Design and cloning strategy of engineered PUS7 plasmids with different localization tags, including 3X FLAG tagged (PUS7-control), nuclear localization signal (NLS), and nuclear export signal (NES) fused 3X FLAG. **c**, Immunofluorescence showing localization of PUS7-control, and PUS7-NLS and PUS7-NES using anti-FLAG antibody. **d**, *pus7*Δ yeast lack Ψ13 and Ψ35 in tRNA which are installed by PUS7, but not Ψ55 (n = 2). **e**, PUS7-control and PUS7-NES yielded similar level levels of Ψ13 and Ψ35 (n = 2). **f**, Density maps showing potential Ψ (Mod-p ID p < 0.001, RNA002) (U-to-C mismatch) distribution across the 5′ UTR, CDS, and 3′ UTR regions in PUS7-NES, PUS7-NLS, PUS7-control (-heat shock (45°C, 15min)), and *pus7*Δ strains. **g–i, l**, Scatter plots showing the number of total direct reads versus the relative occupancy of potential Ψ (% U-to-C mismatch) for PUS7-control (**g**), PUS7-NLS (**h**), and PUS7-NES (**i**), PUS7-control-heatshock (45 ^°^C, 15min) (**l**) as detected by Mod-p ID (p < 0.001). Marginal distributions show U-to-C mismatch levels (top) and log TPM (right). Dot colors indicate 5-mer motifs recognized by PUS7 (blue) and TRUB1 (red). k-mer analysis was performed for high-confidence Ψ sites (p < 0.001 and mismatch level > 40%, RNA002). **j, m**, Venn diagrams showing the overlap of high-confidence Ψ sites between PUS7-NES *vs* PUS7-control and PUS7-NES *vs* PUS7-NLS (**j**), and PUS7-NES *vs* PUS7-control-heat shock (45 ^°^C, 15min) (**m**). **k**, Top 10 GO enrichment for Ψ modified genes in PUS7-NES but not PUS7-control. **n**, Gel showing Ψ incorporation in mRNAs from CLAP experiments, with quantification comparing PUS7^ΔID1^ and *pus7*Δ. For source gel data, see **Supplementary Fig. 8**. Student’s *t*-test (unpaired, two-tailed); data are shown as mean ± SD. *P < 0.05, **P < 0.01, ***P < 0.001 (*n* = 2).

In cells, Ψ is incorporated by a conserved family of enzymes, pseudouridine synthases (PUS enzymes) ^10,17^. Dysregulation of PUS enzyme activity is associated with adverse health outcomes ranging from intellectual disabilities to cancers^19^. Bacterial PUS enzymes are traditionally thought to act only on noncoding RNAs^20^. However, recent evidence suggests that a small subset of *E. coli* mRNAs possessing structural and sequence motifs similar to targeted noncoding RNAs is also modified^21^. Eukaryotic PUS enzymes are distinguished from their bacterial counterparts by additional structural features surrounding their conserved catalytic cores^18^. There are three enzymes responsible for the bulk of mRNA pseudouridylation in eukaryotes: PUS1, PUS4 (TRUB1 in human), and PUS7^22–26^. The full complement of factors dictating how individual eukaryotic PUS enzymes recognize their diverse RNA targets *in vivo* remains to be determined.

Ψ-mapping of eukaryotic transcriptomes and reporter RNAs indicate that PUS1 and the human TRUB1 prefer to modify uridines (Us) within particular secondary structure motifs^22,24^. Conversely, biochemical studies suggest that PUS7 is quite promiscuous in solution, modifying a wide array of substrates whose only requirement for modification is a consensus sequence (UN<u>U</u>AR). Despite this observed promiscuity in solution, PUS7 confoundingly only modifies a small proportion (< 0.3%) of mRNA UN<u>U</u>AR sites inside cells under standard laboratory conditions^23,24^. Unlike PUS1 and TRUB1, the discrimination of PUS7 for a modest subset of targets cannot be explained by mRNA secondary structure preferences alone. These observations, coupled with a report that yeast PUS7 relocalizes upon heat shock, raise the possibility that PUS7 largely selects mRNA targets based on its physical access to UN<u>U</u>AR sites^23,27^. Such a model is not unheard of for mRNA-modifying enzymes, as adenosine deaminases (ADARs) localization impacts the sites of inosine inclusion^30–32^, and m^6^A deposition at consensus DR<u>A</u>CH sequence motifs is suppressed by competition for those sites with exon-junction complexes^33,34^. Here, we use a set of genetically engineered cells with differentially localized PUS7 to investigate how PUS7 compartmentalization impacts mRNA pseudouridylation and protein production. Our findings reveal that shifts in PUS7 localization are associated with changes in mRNA pseudouridylation and protein production in response to metal and ROS stress in yeast and human cells. Additionally, PUS7 localization shifts to the cytoplasm under metal and ROS stress confer cells a fitness advantage, suggesting that mRNA pseudouridylation can have biological consequences.

## RESULTS

### Tools for investigating the effect of PUS7 subcellular localization

We generated yeast cells with differentially localized PUS7 to explore how the location of PUS7 within a cell might impact mRNA biology and cellular fitness. Concerned that changing the compartments of PUS7 with the endogenous PUS7 promoter and/or genetic locus might alter its transcriptional regulation and activity levels, we utilized ectopic (plasmid-borne) expression constructs driven by a strong, constitutive GPD1 promoter. We transformed *pus7*Δ BY4741 *S. cerevisiae* with low-copy (CEN origin) plasmids expressing FLAG-tagged PUS7 with either no C-terminal targeting sequence [*PUS7-control*], a Nuclear Localization Sequence [*PUS7-NLS*], or a Nuclear Export Sequence [*PUS7-NES*] (**Fig. 1b**). PUS7 mRNA and protein levels are similar for all three constructs in *pus7*Δ yeast, and, as expected, higher than endogenous PUS7 in wild-type PUS7 yeast (**Extended Data Fig. 1a-e**). We verified the localization for all constructs by immunofluorescence (**Fig. 1c**). As expected, PUS7-control and PUS7-NLS show predominantly nuclear PUS7-FLAG signal (23% cytoplasmic) while PUS7-NES is shifted to the cytoplasm (63%) (**Extended Data Fig. 1f, Supplementary Fig. 1**).

PUS7 modifies both tRNAs and mRNAs in cells. To evaluate if tRNA modification patterns are altered between our constructs, we used an established nanopore direct RNA sequencing (DRS) method to map the pseudouridylation landscape of tRNAs isolated from PUS7-control, -NLS, and -NES expressing cells^35^. Nanopore sequencing detects RNA modifications by identifying miscalls based on sequence alignment and error models, allowing us to identify locations of tRNA modifications. As expected, *pus7*Δ yeast lacks Ψ13 and Ψ35 in tRNAs^36^, which are known to be modified by PUS7 (**Fig. 1d, Supplementary Fig. 2**). Critically, *pus7*Δ [*PUS7-NES*] and *pus7*Δ [*PUS7-control*] yeast yielded similar levels of Ψ13 and Ψ35 (**Fig. 1e, Extended Data Fig. 2**), indistinguishable from wild-type *PUS7* yeast (**Supplementary Fig. 3-4**).

### PUS7 localization shapes mRNA target selection in S. cerevisiae

To evaluate the influence of PUS7 localization on mRNA pseudouridylation, we generated transcriptome-wide maps of Ψ in mRNAs isolated from cells expressing PUS7-control, PUS7-NLS, or PUS7-NES. Ψ-sites were identified by comparing natively modified mRNA with *in vitro* transcribed (IVT) unmodified RNA sequences using Mod-p ID^37^. Based on previous work, we selected an IVT-control normalized p-value cutoff of 0.001 for all potential Ψ sites, and 40% U-to-C mismatch, along with p-value < 0.001 for high-confidence Ψ sites^37^. We controlled for inherent nanopore sequencing mismatch rates by comparing levels of all nucleotide mismatch rates. Only U-to-C mismatch rates, representing Ψ, differed significantly between cell lines (**Supplementary Fig. 5a**). Similar sequencing depth was attained across cells expressing all three PUS7 variants, with PUS-NES and PUS7-NLS achieving the sequence depth within 91-98% of PUS7-control (**Supplementary Table 1**). Cells expressing PUS7-control and nuclear-localized PUS7-NLS have comparable numbers and distributions of high-confidence mRNA Ψ-sites (81% shared) (**Fig. 1f-h, Extended Data Fig. 3a, and Supplementary Table 2**). We observe an increased number of Ψ-sites relative to those previously reported for cells expressing endogenous levels of PUS7. This is expected given that PUS7 is present at elevated levels in our engineered cell lines (**Extended Data Fig. 1e, 3b**). Despite this, the distribution of PUS7 sites is in line with previous observations showing yeast PUS7 incorporates Ψ sites primarily into mRNA coding regions (CDS, ∼80%) and 5′ untranslated regions (UTRs, ∼13%)^23,24^ (**Extended Data Fig. 3c**). In both PUS7-control and PUS7-NLS expressing cells, the number of Ψ detected in the 3’/5’UTRs and CDS correlates with the prevalence UN<u>U</u>AR motifs in each region (**Extended Data Fig. 3d-f**).

In contrast to PUS7-NLS, mRNAs isolated from cytoplasm-localized PUS7-NES cells exhibit an 11-fold increase in the number of Ψ sites relative to PUS7-control (**Fig. 1f, i**). As expected, k-mer analysis indicates that the most common sequences modified in PUS7-NES cells are within the UN<u>U</u>AR consensus sequence. The majority of Ψ sites map to mRNA CDS and 3’ UTR regions (**Fig. 1f, Extended Data Fig. 3c-d**). In addition to the > 750 additional Ψ sites in mRNAs from PUS7-NES cells, nearly all of the sites mapped in PUS7-control and PUS7-NLS cells were also detected (86% and 88%, respectively) (**Fig. 1j, Supplementary Table 3**). **Ψ** is enriched in mRNAs encoding oxidoreductase and hydrolase activities in transcripts isolated from PUS7-NES, but not PUS7-control cells (**Fig. 1k**). The mRNA poly(A) length and number of detectable introns do not differ between mRNAs isolated from PUS7-NES, PUS7-NLS, and PUS7-control cells (**Supplementary Fig. 5b-d**). Together with the lack of changes observed in tRNA modification status, our data suggest that the nuclear function of PUS7 is primarily maintained in PUS7-NES expressing cells.

### Relocalization of PUS7 under heat shock increases mRNA Ψ-sites

Our supposition that PUS7 localization correlates with Ψ incorporation was further tested by examining the Ψ distribution in cells expressing PUS7-control exposed to heat shock (45 °C). As previously reported, we find that PUS7-control partially localizes to the cytoplasm upon heat shock, where it targets 11-fold more substrates with UN<u>U</u>AR motifs (**Fig. 1f, l, Extended Data Fig. 4a and Supplementary Fig. 6**)^38^. 52% of these additional targets are shared among mRNAs isolated from cells expressing PUS7-NES grown under no stress (**Fig. 1m, Supplementary Table 4**). This suggests that the shared sites arise due to the localization change, while the remaining Ψ sites unique to PUS7-control under heat shock arise from additional regulatory mechanisms (**Extended Data Fig. 4b-c**).

When PUS7-NES cells are exposed to heat shock, the number of Ψ sites increases by an additional 4-fold relative to those observed in mRNAs isolated from unstressed PUS7-NES cells **(Extended Data Fig. 4d)**. The bulk (91.8%) of Ψ sites in PUS7-NES cells identified under normal growth conditions are retained under heat shock (**Extended Data Fig. 4e, Supplementary Table 5**). Analysis of the 1703 mRNAs (containing 3,566 new Ψ sites) uniquely modified in PUS7-NES under heat shock reveals that heat shock-dependent sites are mainly from mRNAs that were not previously modified (**Extended Data Fig. 4e-f**). This is consistent with a model in which some new Ψ-sites arise on previously inaccessible UN<u>U</u>AR motifs upon heat shock. These sites could have been inaccessible for several reasons, including being covered by the ribosome, RNA-binding proteins (RBPs), or RNA structural constraints. The mRNAs only modified in PUS7-NES expressing cells under heat shock are enriched in pathways related to structural molecule activity, oxidoreductase activity, and ligase activity (**Extended Data Fig. 4g-i**).

### Loss of PUS7 insertion domain 1 increases Ψ-sites via relocalization

Eukaryotic PUS enzymes contain additional domains relative to their bacterial counterparts. We investigated the contribution of one such domain, insertion domain (ID1), to PUS7 localization in cells (**Extended Data Fig. 5a**). ID1 was selected for investigation because an endogenous NLS sequence is predicted in ID1 by NLSExplorer^39^ (**Extended Data Fig. 5b**). Our prior work suggested that ID1 might contribute to PUS7 substrate selection^27^. To examine if ID1 alters PUS7 subcellular localization, we imaged *S. cerevisiae pus7*Δ cells expressing fluorescently tagged PUS7 (PUS7-EYFP) and a PUS7 construct that lacks ID1 (PUS7^ΔID1^-EYFP). The population of PUS7^ΔID1^-EYFP localized to the cytoplasm is modestly increased (7%) relative to PUS7-EYFP (**Extended Data Fig. 5c, Supplementary Fig. 7**). DRS analysis of mRNAs isolated from *pus7 Δ* yeast cells expressing untagged PUS7^ΔID1^ revealed a 3.6-fold increase in the number of mRNA substrates modified by PUS7^ΔID1^ relative to PUS7-control (**Extended Data Fig. 5d**). 80% of Ψ sites mapped in PUS7^ΔID1^ overlap with those we observe in mRNAs from PUS7-NES cells (**Supplementary Table 6**).

We sought to corroborate a subset of the novel yeast mRNA Ψ sites we identified in our DRS analyses by the orthogonal biochemical technique, <u>C</u>MC-RT and ligation <u>a</u>ssisted <u>P</u>CR analysis of Ψ modification (CLAP)^40^ (**Extended Data Fig. 6a**). The sites investigated by CLAP were arbitrarily selected from abundant mRNAs with Ψ targeted in cells expressing PUS7^ΔID^ (**Extended Data Fig. 6b**). All sites investigated are not identified as Ψ in DRS analysis of *pus7 Δ*. CLAP assays substantiated our DRS analyses, we found that all nine of the new Ψ sites we investigated were modified with substantial stoichiometries (30-66%) among biological replicates (**Fig. 1n, Extended Data Fig. 6c-d, Supplementary Fig. 8**). These findings further support the hypothesis that the physical access of PUS7 to mRNA UN<u>U</u>AR sequences from PUS7 cytoplasm localization is a key determinant of substrate selection (**Supplementary Fig. 9-10**).

### S. cerevisiae PUS7 cytoplasmic localization enhances fitness

Our data demonstrate that PUS7-NES cells modify many more mRNAs than cells expressing PUS7-control and PUS7-NLS, but do not perturb tRNA modification. We next wanted to ask if selectively altering the Ψ mRNA landscape has cellular consequences. To begin addressing this question, we assessed the fitness of *S. cerevisiae pus7Δ* cells expressing either PUS7-control, PUS7-NLS, and PUS7-NES under 50 different stress conditions (**Supplementary Fig. 11-14**). Cellular fitness was compared through a combination of spot-plating and minimum inhibitory concentration (MIC) measurements. No stress conditions were observed where PUS7-NLS cells demonstrated growth differences from PUS7-control (**Supplementary Fig. 11-14**). In contrast, PUS7-NES cells exhibited decreased sensitivity to seven stressors, including heat shock and stressors associated with metal ion and reactive oxygen species (ROS) stress (Co(II), Zn(II), Ni(II), Cu(II), Fe(II), H_2_O_2,_ cumene-OOH (CU, a strong ROS inducer) (**Fig. 2a-d, Extended Data Fig. 7, Supplementary Fig. 11-12**). Overall, cells with PUS7-NES had 1.2-to 12-fold higher MIC values (Zn(II) and Co(II), respectively) in the presence of these stressors (**Extended Data Fig. 7**). Furthermore, expressing a catalytically inactive version of PUS7 (PUS7-D256A-NES) under a low-expression promoter did not enhance stress tolerance nor Ψ upregulation (**Extended Data Fig. 8, Supplementary Fig. 15**). This demonstrates that the observed stress tolerance instead requires PUS7 catalytic activity.

**Fig. 2.**
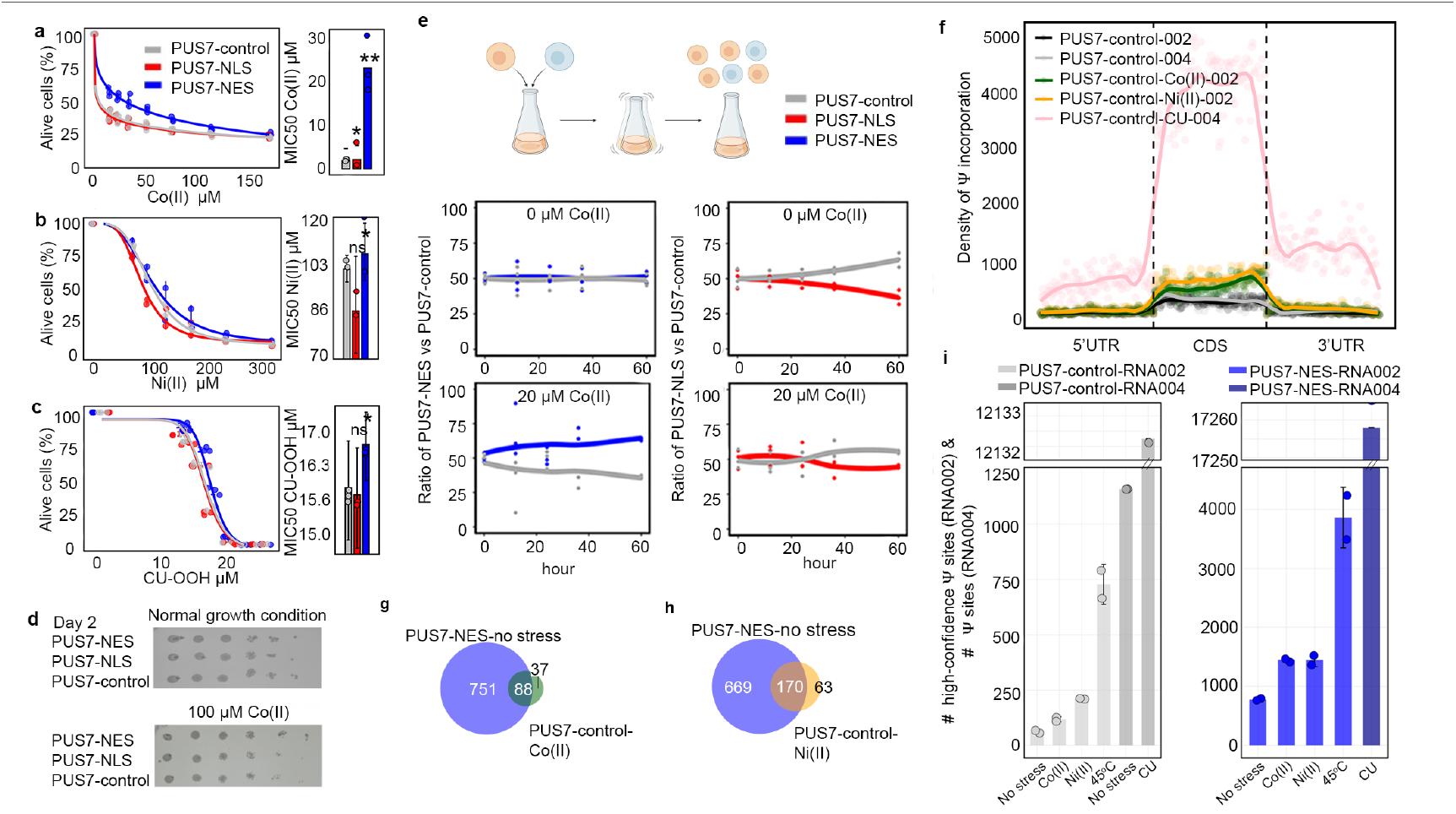
PUS7 relocalization influences cellular stress tolerance by upregulating mRNA Ψ levels. **a-c**, MIC50 test showing stress tolerance increasing toward CoCl_2_ (II), NiSO_4_(II) and CU for yeast with PUS7-NES compared to PUS7-NLS or PUS7-control. Paired t test (two tailed) p-value 0.0049 for NES-Co(II), p-value 0.0464 for NLS-Co(II) (*n* = 3); p-value 0.0482 for NES-Ni(II), p-value 0.1059 for NLS-Ni(II) (*n* = 2); p-value 0.026 for NES-CU, p-value 0.1509 for NLS-CU (*n* = 3). Data are shown as mean ± SD. **d**, Spot plating assay showing the enhanced stress resistance of PUS7-NES yeast to CoCl_2_ for 2 days (*n* = 2). **e**, Schematic of competitive fitness and results showing PUS7-NES/NLS *vs* PUS7-control growth under Co(II) stress and no stress growth conditions. **f**, Density maps of potential Ψ (U-to-C mismatch and dorado fraction) distribution across 5′ UTR, CDS, and 3′ UTR in PUS7-control, PUS7-control–Co(II) (30 μM, 4 h), PUS7-control–Ni(II) (200 μM, 4 h), and PUS7-control–CU (2 mM, 1 h). PUS7-control–Co(II) and –Ni(II) were sequenced with RNA002, while PUS7-control–CU was sequenced with RNA004; PUS7-control was sequenced with both RNA002 and RNA004. Ψ sites were identified using Mod-p ID (p < 0.001) for RNA002 and Dorado modkit fraction >20% (UNUAR motif filter) for RNA004. **g-h**, Venn diagrams showing overlap of high-confident Ψ sites (p < 0.001, U-to-C mismatch > 40% RNA002) between PUS7-NES and PUS7-control-Co(II) (**g**), PUS7-control-Ni(II) (**h**). **i**, Plot of high-confidence Ψ sites in PUS7-control and PUS7-NES under no stress growth condition, Co(II) and Ni(II) (RNA002), and potential Ψ sites in PUS7-control under no stress and CU, as well as PUS7-NES under CU (RNA004), displayed as mean ± SD (*n* = 2).

To further substantiate the findings of our initial screens, we conducted competitive fitness assays in unstressed and Co(II) stress conditions. In these experiments, cultures were inoculated with an equal number of *pus7 Δ* cells expressing plasmids bearing PUS7-control and either nuclear (NLS) or cytoplasmic (NES)-localized PUS7 (**Fig. 2e**). The relative populations of each cell type were measured at time points across 60 hours using qPCR targeting the plasmid expression. We observed that PUS7-control cells outcompeted those expressing PUS7-NLS in the presence and absence of Co(II). However, while cells expressing PUS7-control and PUS7-NES cells grow similarly under unstressed conditions, when grown in the presence of Co(II), PUS7-NES expressing cells outcompete PUS7-control cells (**Fig. 2e**). This finding is consistent with our MIC measurements (**Fig. 2a**).

### mRNA Ψ-sites increase under metal ion and ROS-related stress

We next began exploring if cytoplasmic PUS7-mediated Ψ upregulation might contribute to stress tolerance under divalent metal ion and ROS conditions, by performing direct RNA sequencing on PUS7-control cells treated with either Co(II), Ni(II), or CU. Ψ incorporation is significantly increased in mRNAs isolated from PUS7-control cells grown under these stresses relative to untreated cells (**Fig. 2f, Extended Data Fig. 8)**. Most Ψ sites that we observe were not detected in *pus7*Δ under the same stress conditions (**Extended Data Fig. 8**). Notably, 70% and 73% of high-confidence Ψ sites predicted in mRNAs isolated from Co(II)- and Ni(II)-treated cells, respectively, are also observed in cells expressing PUS7-NES under no stress (**Fig. 2g-h**). These data suggest the possibility that some of the stress-induced Ψ sites might result from PUS7 cytoplasmic relocalization upon these stresses.

PUS7-NES cells exposed to Co(II), Ni(II), and CU exhibit additional Ψ sites, similar to what we observed for PUS7-NES cells exposed to heat shock (**Fig. 2i, Extended Data Fig. 9a-c**). This is consistent with the model we proposed in which PUS7 target selection depends on both PUS7 compartmentalization and substrate accessibility (which fluctuates with changes in translation and RBP-remodeling under stress). GO term analysis of mRNAs specifically modified under Co(II), Ni(II), and CU stress indicates that Ψ is significantly enriched in transcripts encoding proteins related to hydrolase activity, oxidoreductase activity, and ATP hydrolysis activity (p-value < 0.05) in PUS7-NES cells (**Extended Data Fig. 9**). 31% of Ψ enriched mRNAs encode proteins associated with ROS or metal binding (**Supplementary Table 7-12**)^41,42^. Search of the metal protein databank revealed that proteins within these GO-term categories account for the vast majority of the Co(II) binding proteins with known structures (**Extended Data Fig. 9e**)^43^. The accumulation of Ψ sites in these GO-term categories suggests that mRNA pseudouridylation by PUS7 might contribute to these stress-induced cellular responses in cells expressing PUS7-NES.

### Cytoplasmic localization of yeast PUS7 reduces ROS accumulation

Our data indicate that shifting PUS7 to the cytoplasm increases cellular fitness during oxidative stress. To examine the potential contribution of PUS7 localization to mediating the cellular oxidative stress response, we compared cellular ROS levels in *pus7*Δ cells expressing no PUS7, PUS7-control, PUS7-NLS, and PUS7-NES using H_2_DCFDA (DCF). DCF is a colorimetric indicator of ROS that passively diffuses into cells^44^. Under unstressed conditions, cells uniformly exhibit very low ROS levels (**Extended Data Fig. 10a**). However, upon treatment with the ROS-inducing agent CU, we observed a marked difference in ROS accumulation (**Extended Data Fig. 10a-c**). ROS levels increase by 4-, 5-, and 10-fold in cells expressing PUS7-control, PUS7-NLS, and no PUS7, respectively. Comparatively, PUS7-NES cells only exhibit a 2-fold increase in ROS levels (**Fig. 3a)**. Taken together with our Ψ-mapping studies, these results indicate that cytoplasmic PUS7 reduces cellular ROS accumulation under oxidative stress, raising the possibility that the increased incorporation of Ψ by PUS7-NES into ROS-related transcripts might help to mediate cell stress tolerance.

**Fig. 3.**
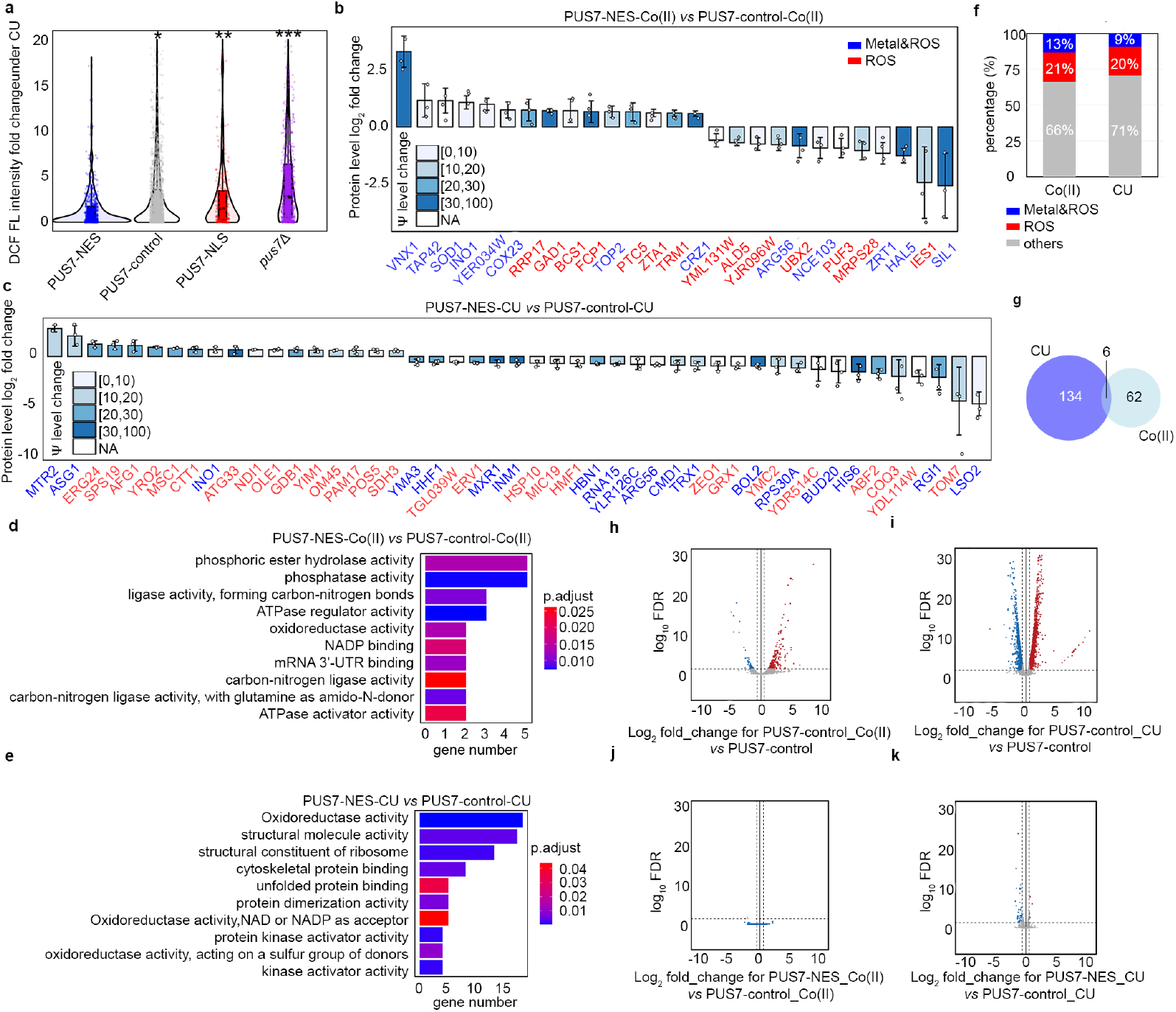
Engineered cytoplasmic PUS7 influences cellular stress tolerance by regulating cellular ROS and protein expression levels. **a**, Cellular ROS level fold change for different groups of cells under CU stress (2 mM 30 min) *vs* no stress growth condition, indicated by DCF (green fluorescence). Unpaired t-test (two-tailed) *P < 0.05, **P < 0.01, ***P < 0.001 (n>150 cells. median (middle line), 25th, 75th percentile (box). **b-c**, Proteomics data showing significant expression level changed proteins (metal-binding/ROS related) and their Ψ level change (shown by bar color) between PUS7-NES and PUS7-control under 30 µM cobalt stress for 4 h (**b**) and 2 mM CU for 1 h (**c**). Metal-binding proteins in blue text label. ROS-related proteins are labeled in red text. Student’s two-sample t-test (unpaired, two-sided). p-value < 0.05 and |log_2_ fold-change| > 1.5 as significant changes (*n* = 3), displayed as mean ± SD. **d-e**, Top 10 GO terms with significant p-values (< 0.05) for proteins with differential expression between PUS7-NES under Co(II) with PUS7-control under Co(II) (**d**), as well as under CU (**e**). **f**, Distribution of expression level changed proteins between PUS7-NES and PUS7-control under 30 µM cobalt stress for 4 h and 2 mM CU for 1 h. **g**, Venn diagrams showing overlap of expression level changed proteins between PUS7-NES *vs* PUS7-control between Co(II) and CU stress. **h-i**, Volcano plots illustrate RNA abundance log_2_fold change between PUS7-control under 30 μM Co(II) for 4 h (**h**) and 2 mM CU for 1 h (**i**) *vs* under no stress growth condition. Student’s two-sample t-test (unpaired, two-sided). Red dots: upregulation; blue dots: downregulation. (n = 2). **j-k**, Volcano plots illustrate RNA abundance log_2_fold change between PUS7-NES under 30 μM Co(II) for 4 h (**j**) and 2 mM CU for 1 h (**k**), and PUS7-control under the same stress. Student’s two-sample t-test (unpaired, two-sided). Red dots: upregulation; blue dots: downregulation. (n = 2).

### The protein landscape is altered by PUS7 cytoplasmic localization

Previous *in vitro* and cell-based reporter studies indicate that the inclusion of pseudouridine in mRNAs can impact translation and splicing^5,12–15,45,46^. These data suggest that the targeted insertion of Ψ into mRNAs by PUS7 might help shape the cellular proteome. We sought to determine if the pseudouridylation of endogenous transcripts can influence the protein landscape by conducting quantitative proteomic studies on *pus7 Δ S. cerevisiae* expressing PUS7-control and PUS7-NES grown in the presence of no stress, Co(II), or CU. As expected, the presence of either stress significantly alters the distribution of proteins in PUS7-control cells (**Extended Data Fig. 10d-g**). In cells grown in Co(II), with differentially localized PUS7 constructs, 68 proteins exhibited expression level changes between PUS7-control and PUS7-NES cells (**Extended Data Fig.11**). The changes were even more dramatic upon CU treatment. We observed 27 proteins with changed expression levels between PUS7-control and PUS7-NLS cells grown in CU. In PUS7-NES cells, 140 proteins exhibit significant differences in their expression levels relative to PUS7-control (**Extended Data Fig. 12**). Notably, PUS7 itself is expressed at higher levels in Co(II) stress, but not unstressed or CU-treated conditions (**Extended Data Fig.11b, 12c**). Overall, 2.4% and 4.8% of all detected proteins (2889 total proteins in Co(II), and 2927 in CU) have changed levels in PUS7-NES cells. These data indicate that PUS7-NES expression is associated with a shift in the cellular proteome under Co(II) and CU stress.

Proteins with differential expression are most likely to have hydrolase and oxidoreductase activities (**Fig. 3d-e**). 9 (in Co(II)) and 13 (in CU) of the proteins with changed levels bind metal and/or are involved in metal transport. Even more proteins (14 in Co(II) and 28 in CU) function in ROS stress pathways (e.g., CRZ1, SOD1, ZRT1, and INO1) (**Fig. 3b-c,f**). The mRNAs encoding proteins with fluctuating levels commonly have altered Ψ modification profiles in PUS7-NES cells relative to PUS7-control. Under Co(II) and CU stress, 91% and 81% of these mRNAs possess Ψ (**Fig. 3b-c, Extended Data Fig. 13**). This is a modest change in Ψ incorporation relative to proteins with unchanged levels (**Extended Data Fig. 13c, f**). There are six proteins whose expression levels change in PUS7-NES cells grown in both Co(II) and CU (ARG1, ARG3, ARG8, ARG56, HXK1, INO1) (**Fig. 3g**). The mRNAs encoding all of these proteins also have their Ψ modification status altered between PUS7-control and PUS7-NES cells under these stress conditions (**Supplementary Table 13**). In general, mRNA pseudouridine incorporations were observed when protein levels both increased and decreased. Similar observations have been made for protein expression from m^6^A-modified mRNAs^47,48^, suggesting that the outcome of pseudouridylation is context dependent. Furthermore, mRNA sequence may play a role in this, as we find that an extended UN<u>U</u>AR motif, ANUGΨAG, is 1.4-fold more prevalent on mRNAs encoding proteins that exhibit altered levels in PUS7-NES cells (**Extended Data Fig. 14**).

We also considered the possibility that proteome changes between PUS7-control and PUS7-NES cells might result from perturbation in tRNA usage. To test this idea, we compared the distribution of codons within mRNAs encoding proteins exhibiting altered expression levels to those that are not changed under Co(II) stress. There were no differences in codon usage between these groups (**Supplementary Fig. 16**). This is consistent with the idea that altered tRNA pools are not primarily responsible for the protein level fluctuations we observe in PUS7-control *vs* PUS7-NES cells.

### Ψ-level changes do not broadly affect mRNA steady-state levels

In addition to assessing protein output, we also investigated the possible correlation between mRNA abundance and pseudouridylation using our DRS data. As expected, we find that RNA abundance is markedly altered between cells expressing any single PUS7 construct grown in unstressed conditions (30°C) *vs* heat shock (45°C), Co(II), Ni(II), or CU (**Fig. 3h-i, Extended Data Fig. 15, Supplementary Fig. 17**). However, the abundance of mRNAs remains generally consistent between yeast cells expressing PUS7-NES or PUS7-control under unstressed, heat shock, Co(II) or CU treatment (**Fig. 3j-k, Extended Data Fig. 16**). This is striking because the levels of Ψ-incorporation and protein expression for many transcripts are altered under these same conditions (**Fig. 1f, Fig. 3b-c**). Forty mRNAs have small changes in their steady-state levels between cells lacking PUS7 (*pus7*Δ) and PUS7-control cells (**Extended Data Fig. 16d**). None of these mRNAs contains predicted Ψ sites (p<0.001), suggesting these RNA level changes are unlikely to be directly driven by mRNA pseudouridylation. Instead, loss of tRNA pseudouridylation in *pus7*Δ cells may drive changes in these transcript abundances. We further performed RT-qPCR to compare the mRNA levels for five abundant transcripts with greater Ψ-incorporation in PUS7-NES cells than in PUS7-control cells grown at 30 ^°^C. No significant differences were observed in the steady-state levels of 4 of these mRNAs, while the abundance of a single mRNA (MRP2) was reduced by 11% in PUS7-NES cells (**Extended Data Fig. 16e-f**). Though we did not directly assess transcription rates or RNA stability, these data suggest that the stress tolerance imparted by PUS7-NES does not operate by influencing mRNA steady-state levels. We posit that these changes more likely arise from alterations in translation and/or RNA-protein binding (**Fig. 3**); however, we cannot rule out the possibility of compensatory changes in transcription or degradation.

### S. cerevisiae PUS7 shifts to the cytoplasm upon ROS-related stress

The Ψ sites specifically mapped in PUS7-NES cells grown in the absence of stress largely overlap with the Ψ sites we observed in mRNA from PUS7-control cells exposed to metal and ROS-related stress (**Fig. 2g-h**). Given this, we explored the possibility that PUS7 might alter its localization patterns upon stress. To examine the subcellular positioning of PUS7 in yeast under different stressors, we monitored the cytoplasmic and nuclear distribution of PUS7-EYFP under 23 stress conditions (CU, tert-butyl-OOH (TB), NaAsO_2_, methylglyoxal (MG), Cd(II), Co(II), Zn(II), Ni(II), Mg(II), Cu(II), NaCl, H_2_O_2_, Hygromycin B, Puromycin, Actinomycin D, Paromomycin, 37°C, 45°C, starvation (15 min and 60 min), 5-fluorouracil (5-FU), Pyocyanin and Thiolutin). PUS7-EYFP was used instead of immunofluorescence because of the inability of *S. cerevisiae* PUS7 to be recognized explicitly by the available human PUS7 antibodies (**Supplementary Fig. 18**). The fraction of PUS7-EYFP in each compartment was quantified with the Cellpose for at least 100 cells^4849^. While changes in PUS7-EYFP localization were not observed for many stress conditions (**Extended Data Fig. 17**), we found nine stresses (Co(II), Zn(II), Cd(II), Ni(II), heat, CU, TB, MG and H_2_O_2_) that induce a partial but considerable shift of PUS7-EYFP to the cytoplasm (**Fig. 4, Extended Data Fig. 18, Supplementary Fig. 19**). Heat, CU and TB promoted the most significant change of PUS7-EYFP to the cytoplasm (increased by 27%, 29% and 12%, respectively). Apart from heat, all of the stresses that induce a cytoplasmic shift in PUS7-EYFP can produce ROS^50,51^. These results are consistent with the observation that CU and heat shock lead to higher levels of Ψ upregulation in PUS7-control (**Fig. 2i**). Furthermore, increased doses of Co(II), Ni(II), Cd(II), CU, MG, TB and NaAsO_2_ promote a higher ratio of cytoplasmic PUS7-EYFP under transient stress (< 4 hours) (**Extended Data Fig. 19, Supplementary Fig. 20**). These results imply that the propensity of PUS7 to alter localization could contribute cellular mechanisms for fine tuning regulatory responses to ROS stress in yeast.

**Fig. 4.**
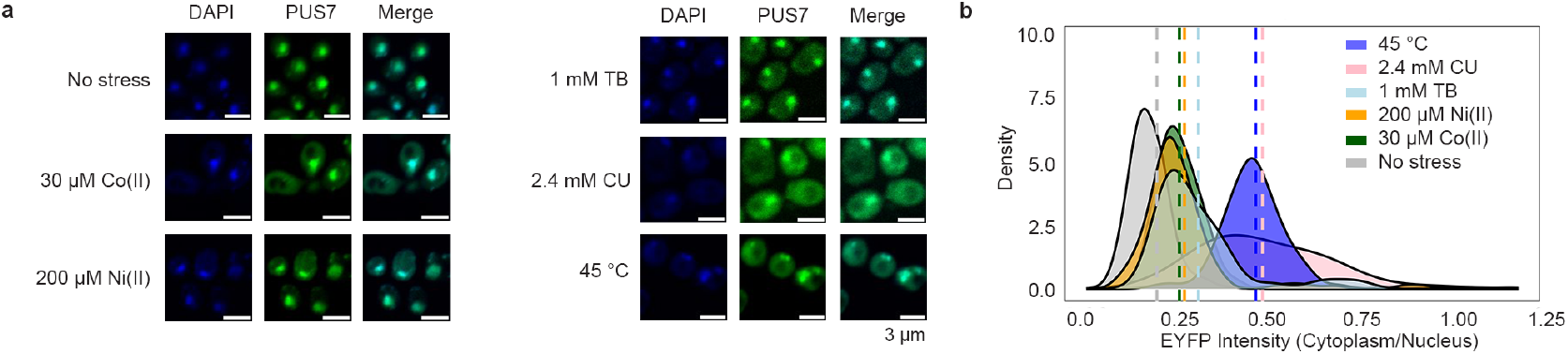
PUS7 relocalization under divalent metal and ROS-related stress conditions. **a-b**, Visualization of PUS7 localization (PUS7-EYFP) under no stress conditions and various stress conditions, including exposure to cobalt (30 µM) for 4 h, nickel (200 µM) for 4 h, 1 mM TB for 30 min, 2.4 mM CU for 1 h, and heat shock (45 °C) for 1 h. Scale bar: 3 µm. Quantification of the Fluorescence Intensity (FI) ratio changes in the cytoplasm *vs* in the nucleus for PUS7 under the no stress growth condition and different stress conditions for more than 100 cells, represented by an intensity map (n >100 cells).

### mRNA Ψ landscape changes in lung epithelial cells upon metal stress

We examined the distribution of Ψ in human mRNAs in a divalent metal-rich physiologically relevant context. BEAS-2B lung epithelial cells were selected as our model because lung cells commonly experience oxidative stress during normal respiration and additional metal stress when exposed to smoking and air pollution. Furthermore, lung cells have previously been shown to contain a high proportion of pseudouridylated sites compared to other human cell types ^40,46–48^. BEAS-2B lung epithelial cells were exposed to a standard environmental pollutant heavy in divalent metal ions (urban particulate matter NIST SRM 1648a, 125 μg/mL for 24 hrs). The location and stoichiometry of Ψ was evaluated with bisulfite-induced deletion sequencing (BID-seq) (**Fig. 5a, Extended Data Fig. 20**)^26,55^. Across all mRNAs, a roughly equivalent number of Ψ sites were detected in unexposed and exposed cells, with 570 and 584 sites, separately, with a similar distribution of Ψ/U stoichiometries (**Fig. 5b,c**, Extended Data Fig. 20b-d). Most human mRNA Ψ sites are in mRNA CDS and 3’ UTRs across both conditions, consistent with previous reports (**Extended Data Fig. 20e**). The modification status of 329 sites (56%) remained essentially unchanged in both conditions, suggesting that these sites are constitutively pseudouridylated. Some transcripts possessed multiple Ψ sites that could be both constitutively modified or unique to either the exposed or unexposed condition. No large shifts in steady-state mRNA levels were observed upon pseudouridylation, akin to our observations in yeast (**Extended Data Fig. 21a-d**). This also excludes the possibility that the unique sites observed in mRNAs from PM-exposed cells arise from changes in RNA abundance (**Extended Data Fig. 21e-f, Supplementary Fig. 21**).

**Fig. 5.**
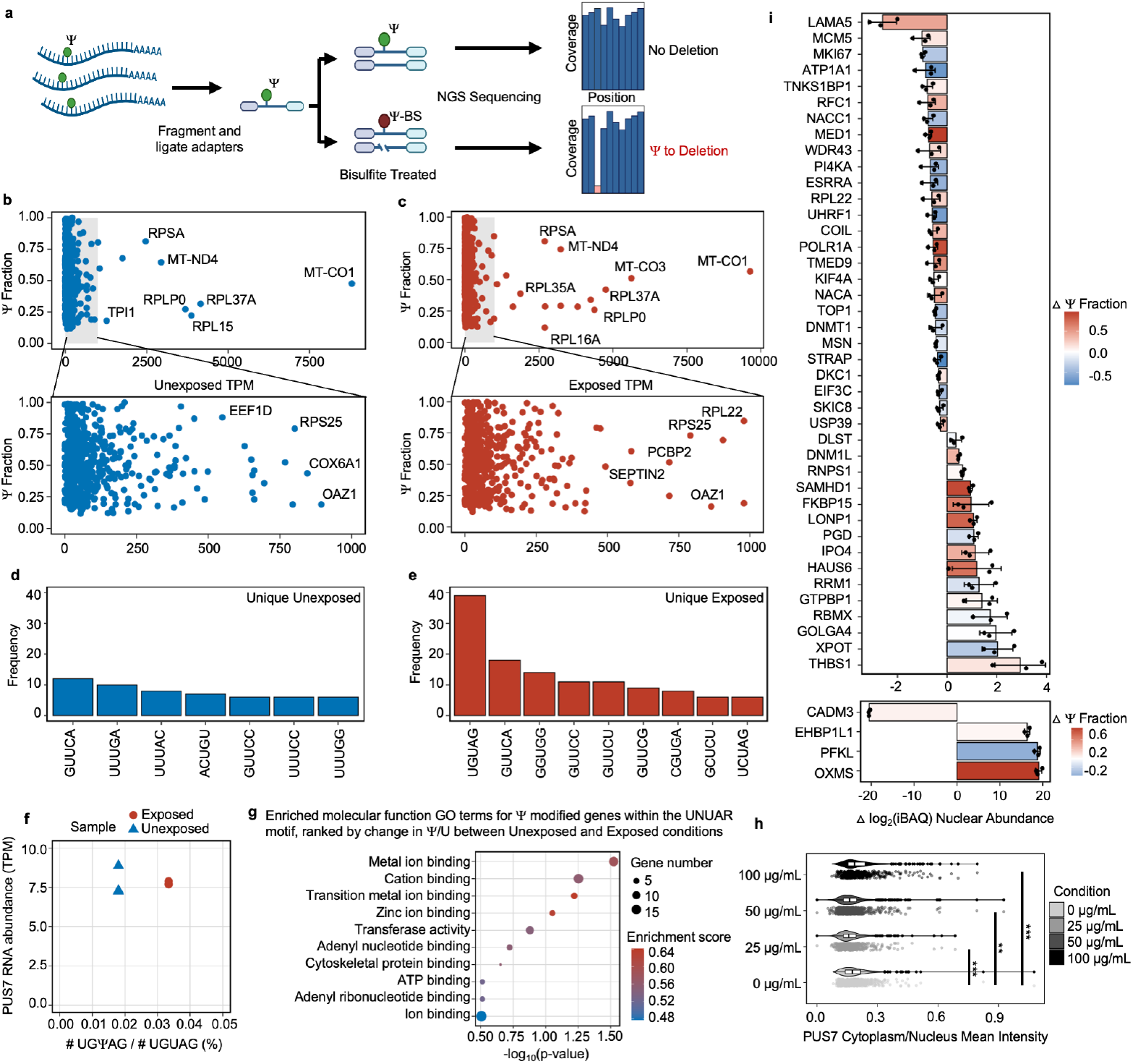
PUS7 partially relocalizes and differentially modifies mRNA transcripts in human lung cells upon exposure to environmental pollutants (urban particulate matter NIST SRM 1648a, 125 μg/mL for 24 hrs). **a**, BID-Seq uses bisulfite treatment followed by next-generation sequencing to determine the location of Ψ sites and the stoichiometry of Ψ to uridine at each site (*n*=3). **b**,**c**, Scatter plot showing genes containing Ψ sites. The x-axis displays a measure of transcript abundance, transcripts per million (TPM), and the y-axis shows the degree of Ψ incorporation (*n*=3). **d, e**, Number of detected Ψ sites that fell within the most common five-nucleotide motifs for the unexposed condition (**d**), or sites unique to the exposed condition (**e**). **f**, Fraction of the Ψ modified PUS7 k-mer (UGΨAG) divided by the total number of UGUAG observed versus the TPM of the PUS7 for unexposed and exposed cells. Student’s t-test (unpaired, two-tailed) (*n* = 3). **g**, Bubble plot showing the GO Molecular Function terms enriched in genes containing modified UNUAR motifs. The input genes were ranked based on the change in the level of Ψ/U modification between the unexposed and exposed conditions and analyzed with gseGO. **h**, Quantification of PUS7 distribution between the cytoplasm and nucleus for individual cells measured through immunofluorescence staining. (Student’s t-test (unpaired, two-sided)) (*n=*400*+ cells per condition*). **i**, Bar plots displaying the change in iBAQ score between the exposed and unexposed conditions for proteins in the nuclear fractions as measured by mass spectrometry. A positive value indicates an increase in protein in the exposed condition relative to the unexposed condition, while a negative value indicates the opposite. Bars are colored to indicate the change in Ψ incorporation between the unexposed and exposed conditions, where a positive value indicates an increase in Ψ incorporation in the exposed condition and vice versa. Student’s t-test (unpaired, two-tailed) (*n = 3*), displayed as mean ± SD.

Consistent with previous studies of human mRNAs, we find that 28% of mRNA Ψ sites fall within the established PUS7 (12%) and TRUB1 (16%) targeting sequence motifs (UN<u>U</u>AR and GU<u>U</u>CN, respectively) (**Extended Data Fig. 21g**)^22–24,26^. However, the distribution of Ψ present in PUS7 and TRUB1 motifs differs between mRNAs isolated from exposed and unexposed cells. In sites exclusive to mRNAs in unexposed cells, the most common motif modified is a TRUB1 consensus sequence GU<u>U</u>CN (**Fig. 5d**). In contrast, sites identified exclusively in exposed cells are highly enriched for the PUS7 consensus sequence UN<u>U</u>AR, with a 0.7-fold increase compared to the UN<u>U</u>AR modified in unexposed cells (**Fig. 5e, f**). As in *S. cerevisiae*, the abundance of PUS7 encoding transcripts and protein is unchanged between conditions (**Fig. 5f, Extended Data Fig. 21h-i**). This indicates that the differences in Ψ incorporation do not arise from changes in PUS7 expression. However, other PUS enzymes have altered mRNA abundance under PM exposure. PUS10, PUS1, and TRUB1 have increased transcript levels, while TRUB2 and RPUSD1 are downregulated (**Extended Data Fig. 21j**). 8%, 9% and 22% of the Ψ-modified mRNAs specific to air pollutant-exposed human lung epithelial cells are also Ψ-modified in *S. cerevisiae* expressing PUS7-NES under Co(II), Ni(II), or CU stress (**Supplementary Tables 14-16**). Furthermore, GO analysis reveals an enrichment in Ψ modified transcripts encoding proteins with metal ion binding (9 %) and oxidoreductase activities (14 %) (p-value < 0.05), similar to yeast (**Fig. 5g and Supplementary Tables 17-19**). UN<u>U</u>AR motifs are not enriched within these pathways. This indicates that the observed enrichment is not due to a high background of UN<u>U</u>AR motifs, but rather reflects that PUS7 modifies more genes within these pathways (**Fig. 5g, Extended Data Fig.21k, Extended Data Fig. 22**).

#### PUS7 partially relocalizes in lung epithelial cells upon metal stress

We investigated the possibility that the redistribution of Ψ sites in mRNAs upon particulate matter exposure might also be promoted (at least in part) by shifts in human PUS7 subcellular localization. To test this idea, we performed immunofluorescence assays with PUS7 antibodies in BEAS-2B cells expressing endogenous levels of PUS7. The cell bodies and nuclei were imaged by phalloidin and DAPI staining (**Extended Data Fig. 23**) and quantified using CellProfiler^56^. Similar to our findings in *S. cerevisiae* (**Fig. 4**), we observe a statistically significant change in PUS7 localization in particulate matter-exposed BEAS-2B cells (p-value < 0.001 under 25 ug/ml, p-value < 0.01 under 50 ug/ml, and p-value < 0.001 under 100 ug/ml, Welch’s t-test) (**Fig. 5h**). We confirmed that these localization changes upon PM exposure were not due to an increase in cellular death by proxy using an anti-ψH2AX antibody to stain for DNA damage events (**Extended Data Fig. 23, Supplementary Fig. 22**). The shifts that we observe are more subtle than in yeast, likely because of the dramatically lower concentration of divalent metal ions in the particulate matter. This is a reasonable supposition given that we see that the extent of PUS7 localization in yeast cells is dependent upon stressor concentration (**Extended Data Fig. 19**). This result shows that PUS7 partial relocalization and Ψ redistribution upon divalent metal ion stress is conserved between yeast and human cells.

### Protein expression changes in BEAS-2B cells upon PM exposure

We next examined if the incorporation of Ψ into mRNAs correlates with protein levels in BEAS-2B cells by conducting mass spectrometry analyses of fractionated cytosolic and nuclear proteins extracted from unexposed and PM-exposed cells. 15 % (406 out of 2669) and 6 % (162 out of 2561) proteins exhibited expression level changes in the nuclear fraction and the cytosolic fraction under PM exposure relative to unstressed conditions (**Supplementary Table 20-24**). These changes assuredly arise from changes in both protein synthesis and stability. As in yeast, the nuclear and cytosolic mRNAs encoding these proteins have a modest enrichment in Ψ (1.3% and 3% increase) relative to proteins with unchanged expression levels (**Fig. 5i, Extended Data Fig. 24, Supplementary Table 25**). Among these differentially expressed proteins, 5 and 6 proteins overlapped between human cells under PM exposure and yeast cells under Co(II) and CU treatment, respectively (**Supplementary Tables 14**,**16**). Overall, our data suggest a model in which the pseudouridylation of mRNAs is associated with cellular proteome-level changes in both yeast and humans.

## DISCUSSION

Enzymatically incorporated chemical modifications are emerging as key modulators of the post-transcriptional life of mRNAs. Studies of two modifications, m^6^A and inosine, largely account for our understanding of how mRNA modifications can impact biological systems. Pseudouridine is nearly as abundant as m^6^A and inosine in mRNAs, but the biological consequences (if any) associated with the inclusion of Ψ in mRNAs have remained enigmatic^2,8^. Here, we demonstrate that the distribution of Ψ in mRNAs fluctuates in response to divalent metal ions and ROS stress. This change in the mRNA modification landscape is enabled by the stress-induced cytoplasmic localization of PUS7, in conjunction with other accessibility-related factors (e.g., RBPs) (**Fig. 6**). We find that incorporation of Ψ into mRNAs correlates with an altered cellular protein landscape. This is consistent with our observation that yeast expressing cytoplasm-localized PUS7 exhibits an increased tolerance to divalent metal ions and ROS stress, and reduced cellular ROS accumulation. Collectively, our data reveal that mRNA pseudouridylation can contribute to changes in the protein landscape under stress, and provide a conserved mechanism for how PUS7 alters Ψ incorporation in mRNA during stress.

**Fig. 6.**
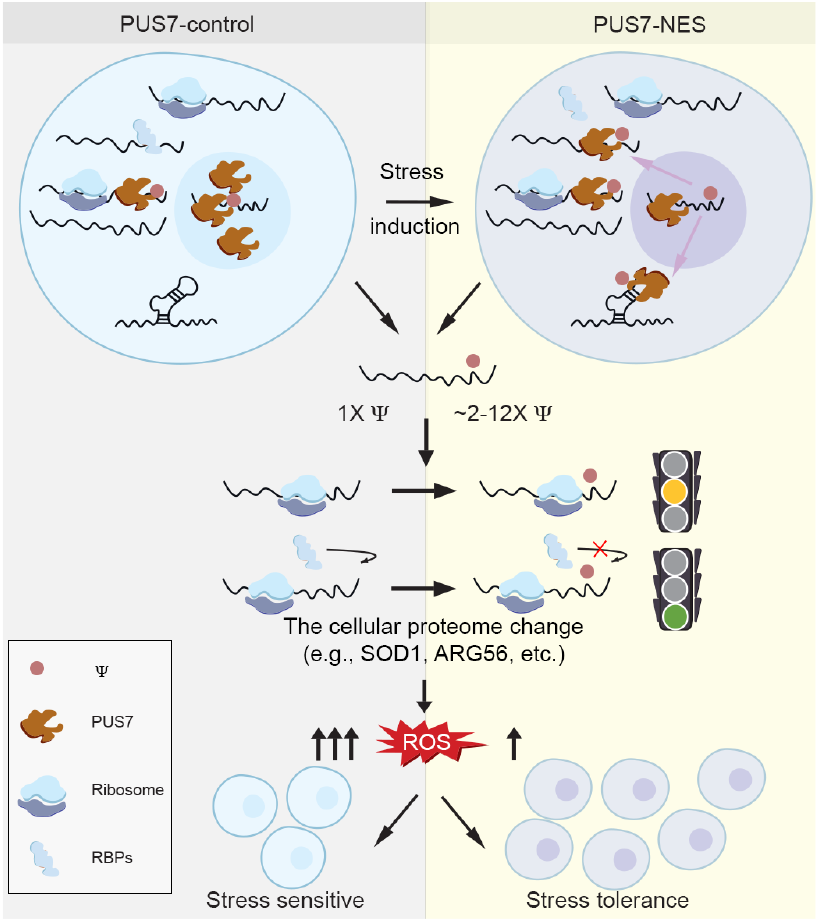
Model of PUS7 relocalization regulating cellular fitness under stress. PUS7 localizes into the cytoplasm upon heat shock, divalent metal ion/ROS stress, which adds Ψ to more mRNA targets, especially ROS-related mRNAs. This impacts the cellular proteome, with Ψ directly perturbing translation, and/or protein-RNA interactions that modulate protein synthesis (e.g., PUF proteins^75^). This further reduces the accumulation of cellular ROS and induces cell stress tolerance.

The discovery that PUS7 localization shapes mRNA targeting is supported by our previous biochemical investigations, indicating that PUS7 is a promiscuous enzyme *in vitro*^27,28^. This work suggested that PUS7 *in vivo* substrate selection may be determined by its access to a given UN<u>U</u>AR sequence. Our findings are similar to observations that the subcellular location of ADARs acting on RNA is a major determinant of their substrate selection^31,32^. Given that the yeast homolog of TRUB1, PUS4, relocalizes into the cytoplasm and adopts a prion state to confer a fitness advantage upon nutrient starvation, we posit that PUS enzyme localization may be a common determinant of PUS *in vivo* mRNA substrate selection^57^. Additionally, PUS4 and PUS7 relocalize under different conditions, leading us to suggest the possibility that different PUS enzymes might be best suited to help cells mitigate different types of stress. In line with this supposition, our data reveal that the levels of mRNAs expressing a variety of PUS enzymes change in response to varying stress conditions in both yeast and humans (**Extended Fig. 21j and Supplementary Fig. 23**).

Our data indicate that ROS and metal ion stress promote the partial relocalization of PUS7 to the cytoplasm, where it modifies more mRNAs involved in ROS and metal ion homeostasis. Since ROS and metal ion homeostasis are intertwined, exposure to metal ions like Co(II) can cause oxidative stress^51^. If mRNA pseudouridylation is biologically consequential, then increased pseudouridylation could reasonably be predicted to enhance cellular stress tolerance and lower cellular ROS levels. Indeed, that is what we observe (**Fig. 2-3**). Notably, the tRNA modification status at Ψ13 and Ψ35 is not altered when PUS7 partially relocalizes, indicating that the biological consequences of PUS7 pseudouridylation that we observe in PUS7-NES cells may be mRNA mediated (**Fig. 1d-f**). While these data cannot directly ascribe a specific role for PUS7 in the ROS stress response, they support the hypothesis that mRNA pseudouridylation can have biological consequences in cells. The idea that PUS7 might contribute to ROS stress is especially intriguing given that we observe PUS7 relocalization in human BEAS-2B cells upon PM exposure, which is a form of ROS stress.

In contrast to m^6^A, we find that Ψ-inclusion does not broadly shift mRNA steady-state levels (**Extended Data Fig. 16**). However, proteins that undergo expression level changes under ROS stress are more likely to be encoded by pseudouridylated mRNAs in both *S. cerevisiae* and BEAS-2B cells. We find that pseudouridine incorporation levels vary (up and down) in mRNAs in a stress-dependent manner. Similarly, protein levels shift in both directions (higher and lower) from mRNAs when Ψ-incorporation levels shift (**Fig. 3b-c, Extended Data Fig. 13**). Fluctuations in pseudouridylation are also observed on mRNAs encoding proteins that do not exhibit changes in their steady-state levels, however, to a lesser degree. These observations suggest that the incorporation of Ψ does not always have a uniform effect on protein expression. Nonetheless, when it does exert an effect, our data suggest a model in which Ψ either directly perturbs translation, and/or influences protein-RNA interactions that modulate protein synthesis (**Fig. 6**). Consistent with this, a handful of studies indicate that Ψ can slow the ribosome in a context-dependent manner^13,58,59^. Additionally, analysis of mRNAs targeted in PUS7-NES, but not in PUS7-control cells, using BioGRID^60^, shows that these mRNA encoded proteins have physical interactions with proteins whose levels change during stress (**Supplementary Fig. 24**).

Our observations are reminiscent of sites for other key modulators of gene expression – including m^6^A, inosine, and miRNAs – which have varying (or no) effect on protein expression depending on their sequence context, stoichiometry of incorporation, and available RNA protein binding (RBP) networks^3,61–63^. Similarly, chemical modifications in proteins (e.g., methylation, acetylation) also do not exert uniform effects on protein function and stability^64^. Our findings raise the possibility that Ψ incorporation into mRNAs does not always contribute to modulating protein expression linearly or directly – by exerting a significant influence on a small number of genes - but instead acts in a complex and context-dependent manner.

## METHODS

### Experiments performed in *S*. *cerevisiae*

#### Strains and culture conditions

*S. cerevisiae* strains BY4741 and BY4741 *pus7*Δ::*kanMX* were obtained from Horizon Discovery and validated by PCR using primer sequences in **Supplementary Table 26**. Yeast were cultured according to standard procedures. Plasmids were introduced by lithium acetate transformation and selected/maintained by growth in CSM-URA. Three independent clones were isolated from each transformation and preserved as glycerol stocks (The transformation efficiency for each group is around 40–80% (P4-P8)). Biological replicate experiments used independent clones.

#### Plasmid construction

The *PUS7* ORF was PCR amplified from BY4741 genomic DNA and cloned into pENTR/D-TOPO (Invitrogen, K240020). The D256A mutation was introduced into the *PUS7* ORF by site-directed mutagenesis using PfuTurbo (Agilent 600250). pAG416GPD-ccdB-EYFP^65^ was a gift from Susan Lindquist (Addgene plasmid # 14220; http://n2t.net/addgene:14220; RRID:Addgene_14220). Synthetic fragments bearing 3xFLAG, 3xFLAG-NES, and 3xFLAG-NLS coding sequences were purchased from IDT (see Table S1) and inserted via NEB HiFi assembly (NEB E5520) into pAG416GPD-ccdB-EYFP to generate pAG416GPD-ccdb-3xFLAG, pAG416GPD-ccdb-3xFLAG-NES, and pAG416GPD-ccdb-3xFLAG-NLS. NEB HiFi assembly was used to replace the GPD promoter with the minimal *CYC1* promoter in all three localization plasmids. Gateway cloning (Invitrogen™ 11791019) was used to introduce the *PUS7* and *pus7(D256A)* coding sequences into each of the six pAG416GPD and pAG416CYC localization vectors. Plasmid sequences are presented in **Supplementary Table 27** (As a control, we prepared *pus7*Δ [*GPD1-PUS7(D256A)-control*]. To our surprise, this strain expressing the catalytically inactive PUS7(D256A) demonstrated a dramatic slow growth phenotype, to the extent they were experimentally intractable (**Supplementary Data Fig. 15**). This led us to utilize *pus7*Δ [*GPD1-EYFP*] as a control for many experiments, since expression of a recombinant protein from a strong constitutive promoter can be a stressor in its own right. Astute readers will note that this is far from a perfect negative control, as PUS7 may possess non-enzymatic activities that could contribute to observed phenotypes. To address this concern, we prepared six additional strains in which PUS7 expression is driven by a weak, constitutive version of the *CYC1* promoter: *pus7*Δ [*CYC1-PUS7-control*], *pus7*Δ [*CYC1-PUS7(D256A)-control*], *pus7*Δ [*CYC1-PUS7-NLS*], *pus7*Δ [*CYC1-PUS7(D256A)-NLS*], *pus7*Δ [*CYC1-PUS7-NES*], and *pus7*Δ [*CYC1-PUS7(D256A)-NES*]. These strains all demonstrated equivalent growth under normal conditions (**Supplementary Data Fig. 15**) and were used to exclude the key phenotypes observed in the [*GPD1-PUS7*] strains.

#### Spot plating

Yeast cultures were grown overnight in CSM-URA, then diluted to an OD_600_ of 1.0. Five-fold serial dilutions were prepared, and 3µL of each dilution was spotted onto the relevant plate. Plates were incubated for 2 to 4 days at 30°C before photography.

#### MIC50 assays

Overnight yeast cultures in CSM-URA were diluted to an OD_600_ of 0.1 in CSM-URA containing inhibitors at the indicated concentrations in 96-well plates. Plates were incubated at 30°C with shaking at 900 rpm for 12 hours, before absorbance measurement on a SpectraMax iD3 Multi-Mode Microplate Reader. The IC_50_ was calculated using the following equation:

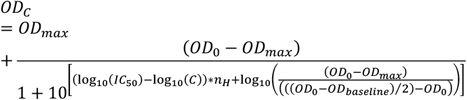

Wherein *C* is the concentration of inhibitor, *OD*_*C*_ is the final OD_600_ of a culture with inhibitor concentration *C, OD*_0_ is the final OD_600_ of culture without inhibitor, *OD*_*max*_ is the final OD_600_ of the culture containing the maximum tested concentration of inhibitor, *OD*_*baseline*_ is the starting OD_600_ of all cultures, and *IC*_*50*_ is the concentration of inhibitor at which 50% growth inhibition is observed.

#### Competitive fitness

Experimental variables tested include PUS7 localization (NES vs NLS), stress treatment (no stress *vs* cobalt exposure), and time points. Yeast were individually cultured for 48 hours in CSM-URA, then diluted to OD_600_ 0.05 and grown for 24 hours to obtain log-phase cells. These cell suspensions were diluted to an OD_600_ of 0.2 and combined pairwise in equal volumes. Two-fold serial dilutions of the co-cultures were prepared and incubated. Every twelve hours, a mid-log phase coculture was identified (OD between 0.1 and 0.4), a sample was taken, and the coculture was serially diluted for further incubation until the total coculture time was 60-72 hours. Plasmid DNA was extracted (Zymo D2004), and the relative abundance of each strain in the co-culture was determined by qPCR using primers specific to each Pus7 localization construct (Thermo K0392). Primer sequences are in **Supplementary Table 28**.

#### Imaging

Mid-log phase yeast were fixed with 3.7% (w/v)(final) formaldehyde at room temperature for 30 minutes, then collected by centrifugation. Fixed cells were washed four times with ice-cold SPC buffer (1.2 M sorbitol, 128 mM dibasic potassium phosphate, 33 mM citric acid) before being attached to poly-L-lysine-coated slides in the same buffer. Slides were dehydrated in -20°C MeOH for 5 minutes and air dried. Intrinsically fluorescent samples were mounted with ProLong™ Gold Antifade Mountant with DAPI (Invitrogen™, P36941) according to manufacturer instructions and imaged.

For immunofluorescence, slides were rehydrated in a humidity chamber and then incubated with blocking buffer (5% normal goat serum, 1% Tween-20, 1X PBS). Slides were incubated with a 1:500 dilution of Abcam #ab18230 in blocking buffer overnight at 4°C, then washed four times with blocking buffer. A one-hour incubation with a 1:1000 dilution of Goat Anti-Mouse IgG H&L (Abcam #ab150115) was performed, followed by four washes. After drying, slides were mounted with ProLong™ Gold Antifade Mountant with DAPI (Invitrogen™, P36941). Images were captured using a Nikon A1 inverted confocal Microscope using 408 nm channel and 647 nm channel with oil immersion at 100X magnification (n=119).

#### Localization quantification

Yeast PUS7 localization quantification was performed with Cellpose^49^. The whole cell region was defined using the bright field channel, and the nucleus region using the 408 nm binary images. Scripts were used to calculate the fraction of Pus7 signal in the cytoplasm and are available on GitHub (https://github.com/mlruan/Imaging_data_quantify).

#### Intracellular ROS measurements

Mid-log phase yeast were harvested and treated with stressor as appropriate, then stained with 10 μM of 2’,7’-dichlorofluorescein diacetate^44^ for 30 minutes. Cells were collected, washed twice with SPC buffer, attached to a poly-L-lysine-coated slide, and mounted with ProLong™ Gold Antifade Mountant with DAPI (Invitrogen™ P36941). Quantification was performed with Cellpose, using the bright field channel to define whole cell boundaries and the 488 nm channel for the ROS signal.

#### Ligation-assisted PCR analysis of Ψ modification (CLAP)

The published protocol^40^ was followed exactly. CLAP primer sequences were designed using Primer3^66^ and checked for specificity using primer BLAST. The CLAP primers (designed by primer3 for PAF1, PRO2, ABF1, OTU1, FAS1, MIS1, MRPL36, NMT1, and DCW1), adaptor, and splint sequences are listed in **Supplementary Table 29**.

#### Western blots

Mid-log phase yeast were harvested and broken with acid-washed glass beads in STE buffer (10 mM Tris-Cl, 10 mM EDTA, 0.5% SDS, pH 7.5) at 4°C, then clarified at 10,000×*g*. Lysate concentrations were determined by Bradford assay (Bio-Rad 5000201). Between 5 and 50 µg of total protein per sample were separated by SDS-PAGE and transferred to a PVDF membrane (MilliporeSigma™ IPFL07810) using a semi-dry apparatus (100V for 60 minutes). The membrane was stained with LI-COR Revert™ 700 Total Protein Stain (LI-COR 926-11011), imaged, and destained according to the manufacturer’s protocol. Membranes were blocked (LI-COR 927-60001), then incubated with a 1:1,000 dilution of anti-DDDDK antibody (Abcam #ab18230) in blocking buffer with 0.2% Tween 20 overnight at 4°C. The membrane was washed three times in 1X TBS with 0.1% Tween 20. The secondary antibody was Novus Biologicals™ (#NB120519) at a 1:1,000 dilution in blocking buffer with 0.2% Tween 20 and 0.01% SDS. After three more washes, membranes were imaged on a Bio-Rad ChemiDoc MP.

#### Proteomics

The yeast proteomics experiment was performed by the Michigan Proteomics Core. Quantitative analysis of yeast proteins was performed with a Data Independent Acquisition workflow using the spectrum library approach described by Pino *et al*^67^. Biological triplicate samples were processed using an LC-MS method with 30 30-minute gradient per replicate. A t-test was employed to obtain the p-value.

#### Preparation of total RNA, tRNA, and mRNA

Mid-log phase yeast were harvested by centrifugation and flash frozen with liquid nitrogen. Pellets were resuspended in ice-cold 60 mM sodium acetate, 8.4 mM EDTA, pH 5.2, then extracted three times with equal volumes of 125:24:1 phenol:chloroform: isoamyl alcohol and 1% (final) SDS, and twice with chloroform. Nucleic acids were collected by ethanol precipitation and treated with DNase I (Roche 04716728001), then precipitated again to yield total RNA. tRNA was prepared by separating total RNA via anion exchange chromatography on a Resource Q column (Cytiva). tRNA-containing fractions were pooled and concentrated by ethanol precipitation. Poly-A RNA enrichment of RNA was performed on total RNA using *mRNA quantification by RT-qPCR:* 2 µg total RNA was reverse transcribed with RevertAid First Strand cDNA Synthesis Kit (Thermo Scientific™, K1622) using random hexamer primers to produce cDNA.1 μL of the produced cDNA was used to perform a RT-qPCR with Luminaris Color HiGreen qPCR Master Mix (Catalog number: K0392) according to the manufacturer’s protocol. Primers sequences for AIM6, ATP5, MRP2,MRSP18, RPT5, RDN18-1are in **Supplementary Table 28**. Samples were loaded in a 364-well plate and run in an Applied Biosciences 7900HT RT PCR System. This experiment was conducted with three biological replicates, and normalization was performed using the signal from 18S ribosomal RNA.

#### Nanopore sequencing of yeast mRNA

Some experiments were performed before RNA004 flow cells became commercially available, while others were performed after RNA002 flow cells were discontinued. Yeast mRNAs were extracted as described above and prepared for sequencing using the Direct RNA Library Kits (ONT SQK-RNA002 for RNA002 and SQK-RNA004 for RNA004). The in vitro transcribed mRNA (IVT-mRNA) control library was prepared as follows. mRNA was extracted from *pus7*Δ [pAG416 *GPD1p-PUS7-control*] and converted to a cDNA library using a cDNA-PCR Sequencing Kit (ONT SQK-PCS109). Nanopore adapters were ligated to the cDNA library using T4 DNA Ligase (NEB M0202M). Reverse transcription to generate the IVT-mRNA library was performed with SuperScript III Reverse Transcriptase (Invitrogen 18080044) for RNA002 and Induro® Reverse Transcriptase (NEB M0681) for RNA004. The IVT-mRNA library was prepared for sequencing using the same procedure as for cell-derived mRNA. Direct mRNA sequencing was performed on a Promethion device controlled by MinKnow v19.12.5 (ONT Flow Cells R9.4.1 for RNA002, R10.4.1 for RNA004) using 500 ng of poly(A) RNA ligated to the ONT RT adaptor. Multi-fast5 files generated by RNA002 were basecalled using Guppy v3.2.10 with the high-accuracy model (rna_r9.4.1_70bps_hac.cfg). Multi-pod5 files generated by RNA004 were basecalled using Dorado v0.9.0 with the high-accuracy model (rna004_130bps_sup@v5.1.0) and -- modified-bases pseU. In both cases, reads were mapped to the *S. cerevisiae* reference genome (S288C_reference_sequence_R64_4_1_20230823) using Minimap2 v2.17 with options -ax splice -uf -k14.

#### Pseudouridine detection in yeast mRNA

Ψ sites were detected in RNA002 data using Mod-p ID v1.0.0 (https://github.com/RouhanifardLab). The U-to-C mismatch frequency was calculated by comparing native RNA sequences against IVT controls. The significance (p-value) of each U site was determined based on the number of reads and the mismatch level. Sequence-dependent expected error (IVT_kmer_Analysis.csv, **Supplementary Table 30**) and read coverage were used to correct mismatch frequencies. Potential Ψ sites were defined by the following criteria: >10 reads, p-value < 0.001. High-confidence Ψ sites were defined by the following criteria: >10 reads, normalized U-to-C mismatch > 40% (native – IVT), p-value < 0.001, IVT control U-to-C mismatch < 10% with >7 reads coverage. Transcript abundance (TPM) was calculated using StringTie. Ψ sites were detected in RNA004 data using modkit v0.5.0. Potential Ψ sites were defined using the following criteria: >20 reads and --modified-bases pseU fraction > 20% and UNUAR motif filter for PUS7 target analysis.

#### Nanopore sequencing of tRNAs and data processing

We followed published, detailed methods^68,69^ for Nanopore sequencing and modified base calling of tRNAs. Briefly, direct tRNA sequencing was performed on 300 ng of tRNA ligated to the ONT RT adaptor and sequenced on a PromethIon (ONT Flow Cell R10.4.1) controlled by MinKnow 24.06.16. The version of Dorado used for basecalling was v0.8.2 (rna004_130bps_sup@v5.1.0).

#### Gene ontology, sequence logo, and plot preparation

Gene Ontology (GO) analysis of Molecular Function was performed using R with the GO file downloaded from geneontology.org for yeast. The sequence motifs were generated by the seqLogo with the calculated kmers (with Ψ calling rate over 40%) as input. All the plots for Nanopore Direct RNA sequencing were generated through R ggplot2.

### Experiments in BEAS-2B cells

#### Culture conditions

BEAS-2B cells (ATCC CRL-9609) were cultured from cryopreserved stocks in collagen-coated T-75 culture flasks according to ATCC guidelines. Cells were seeded at 3,000 cells/cm^2^ and cultured in 23mL of Bronchial Epithelial Cell Growth Medium (BEGM) (Lonza, CC-3170) without gentamicin-amphotericin. Cells were grown at 37°C in a humidified incubator with a 5% CO_2_ atmosphere, and complete media exchanges were performed every 48 hours. After approximately 4 days, the cultures reached ∼70% confluency, and cells were sub-cultured into 6-well plates coated with Type 1 collagen (Advanced BioMatrix, Cat#5005) and allowed to attach to the growth surface for 24 hours before exposure to particulate matter.

#### Particulate Matter Exposure

The BEAS-2B cells were exposed to an urban particulate matter mixture purchased from NIST (SRM 1648a). Just before the start of the exposures, the PM mixture was weighed using an analytical balance and suspended in sterile DI H2O in a 5 mg/mL stock solution. The suspension was sterilized by UV irradiation for 30 minutes. The stock was diluted with BEGM medium to reach the tested concentrations of 125 µg/mL. To begin the exposures, media from the well plates was removed and replaced with equal volumes of PM-containing media for exposed cells or fresh media for unexposed control cells. The cells were incubated at 37°C in a humidified incubator with a 5% CO_2_ atmosphere for a 24-hour exposure period before downstream analysis.

#### Immunofluorescence

BEAS-2B cells adhered to chamber well slides (Lab-Tek, 177402), precoated with type 1 collagen (Advanced BioMatrix, Cat#5005) and fibronectin (VWR, 103700-654), and exposed to particulate matter as described above, 24 hours after seeding. Following exposure, cells were washed with pre-warmed PBS for 5 min, then fixed by incubation for 15 min at 37°C in a freshly prepared, methanol-free 4% formaldehyde solution (Thermo Scientific, 28906) in PBS. Cells were rinsed 3× with PBS before being permeabilized by incubation in a 0.1% Triton-X PBS solution for 5-10 min. Cells were again rinsed 3× with PBS and then blocked with 1% bovine serum albumin (BSA) in PBS for 1 h at room temperature (RT). Cells were subsequently incubated overnight at 4°C, rocking in a 1% BSA in PBS solution containing a 1:200 dilution of anti-PUS7 mouse primary antibody (Invitrogen, MA5-25265) and a 1:400 dilution of anti-γ-H2AX rabbit primary antibody (Abcam, AB81299). Cells were then washed 3×, 5 min in PBS and incubated for 1 h, protected from light at RT in a 1% BSA in PBS solution containing a 1:500 dilution of goat anti-mouse IgG (H+L) AlexaFluorTM 488 secondary antibody (Invitrogen, A11001) and a 1:500 dilution of donkey anti-rabbit IgG (H+L) AlexaFluorTM Plus 647 secondary antibody (Invitrogen, A32795). Cells were again washed 3×, 5 min in PBS. Afterwards, cells were stained with AlexaFluorTM 594 Phalloidin (Invitrogen, A12381) and mounted using ProLong Gold antifade reagent with DAPI (Invitrogen, P36935) according to manufacturer protocols to enable visualization of the F-actin structure and nuclei, respectively. Coverslips were sealed onto microscope slides using clear nail polish. Slides were stored in the dark at 4°C until imaging. Fluorescent images were acquired using a Nikon W1 Spinning Disk Confocal Microscope at 20× resolution using 4 laser lines (405 Diode: DAPI Nuclear Stain; 488 Diode: AlexaFluorTM 488 PUS7; 561 Diode: AlexaFluorTM 594 Phalloidin/Actin Stain; 640 Diode: AlexaFluorTM 647 γ-H2AX).

#### Localization quantification

CellProfiler^70^ was employed to determine the distribution of PUS7 between the nucleus and cytoplasm. Briefly, the nuclear area was segmented using the DAPI stain (405 channel), and the cell body area was determined by subtracting the nuclear area from the area within the Phalloidin stain (561 channel). Cells were filtered only to include those with a mean PUS7 (488) intensity greater than 0.03 in subsequent analyses. The mean intensity in the 488 channel was used to quantify the PUS7 abundance within each segmented area. A ratio was calculated for each cell by dividing the mean cytoplasmic intensity by the mean nuclear intensity. Similarly, CellProfiler was used to quantify the per-cell mean intensity of γ-H2AX signal through the 640 channel.

#### Western blotting

Total protein was extracted from adherent cells using M-PER (Thermo Scientific, 78501) supplemented with Halt Protease Inhibitor Cocktail (Thermo Scientific, 78429). Protein extracts were quantified through the Bradford assay (Thermo Scientific, 1863028). SDS-PAGE separated ten micrograms of total protein per sample, then transferred to a nitrocellulose membrane using a semi-dry apparatus (35 minutes at 17 V). The membrane was blocked with 5% (w/v) BSA in 1X TBS, then washed twice with TBS + 0.5% Tween 20, then twice with TBS. The first primary antibody was 1:10,000 anti-GAPDH (Proteintech, 10494-1-AP) in 1% BSA + TBS.

After a 2-hour room temperature incubation, the membrane was washed as before, then incubated for one hour at room temperature with 1:5,000 HRP-conjugated secondary antibody (Invitrogen 31460) in TBS with 1% BSA. The membrane was washed as before, then incubated overnight at 4°C with a 1:2,000 dilution of anti-PUS7 (Invitrogen MA5-25265) in TBS with 1% BSA. After washing, the membrane was incubated with 1:2,500 HRP-conjugated secondary antibody (Promega W4201) for 90 minutes, then washed again. Chemiluminescent detection was performed using the Clarity Western ECL substrate (Bio-Rad 1705060) on a Bio-Rad ChemiDoc XRS+ and quantified using ImageLab software (Bio-Rad).

#### Proteomics

BEAS-2B cells were grown and subjected to PM exposure/unexposed control as described above. Cytoplasmic and nuclear protein fractions were extracted in biological triplicate from PM-exposed and unexposed cells using the NE-PER Nuclear and Cytoplasmic Extraction Reagents (ThermoFisher, 78833) following the manufacturer’s protocol. Halt Protease Inhibitor Cocktail (ThermoFisher, 78425) was added to the extraction reagents to prevent degradation of extracted proteins. Nuclear and cytoplasmic protein extracts (35 μg/sample) were submitted to the University of Texas at Austin Biological Mass Spectrometry Facility for LC-MS/MS analysis to determine relative protein abundances in each subcellular fraction for both PM-exposed and unexposed cells. Samples were processed for LC-MS/MS analysis following a modified version of the SP4 protocol that can be found at (https://wikis.utexas.edu/display/proteomicscore/SP+4+Protocol). Briefly, samples were denatured in 6M urea and mixed at 25ºC for 10 minutes, 800 rpm. Next, samples were reduced by adding TCEP to a concentration of 5 mM and mixed at 25ºC for 20 minutes, 800 rpm. Samples were alkylated by adding 2-chloroacetamide to a concentration of 10 mM and mixed at 25ºC for 20 minutes, 800 rpm in the dark. Afterwards, 1 mL of ice-cold acetonitrile was added, and samples were gently vortexed. Samples were stored overnight at -20ºC. The next day, samples were pelleted through centrifugation, and the supernatant liquid was removed. Samples were washed thrice in 80% ethanol through centrifugation and decanting, with care taken to avoid disrupting the pellet. Digestion was performed overnight at 37 °C, 600 rpm by resuspending samples in ammonium bicarbonate digestion buffer and adding trypsin at a ratio of 1 ug trypsin:50 ug protein. The next day, samples were pelleted by centrifugation and transferred to a fresh tube. Acetonitrile was added, and samples were dried by vacuum centrifugation. To remove additional salts, samples were processed using the Ziptip protocol (https://wikis.utexas.edu/pages/viewpage.action?pageId=56332998). Afterwards, samples were resuspended in 0.1 % formic acid in water and used as input for LC-MS analysis. Samples were run on the Dionex LC and Orbitrap Fusion 2 (Thermo Scientific) with a 180-minute run time. Raw data files were analyzed using MaxQuant against the human proteome. Output data was filtered using a minimum Andromeda score threshold of 3064. Differences in nuclear and cytoplasmic protein abundances for PM-exposed versus unexposed samples were facilitated by comparing the log2FoldChange of iBAQ values generated from LC-MS/MS. A Student’s T-test was used to evaluate statistically significant changes in log2(iBAQ) values between samples.

#### Genetic interaction analysis

The genetic interaction between the genes incorporating high Ψ levels (U to C mismatch level >30%), as determined through nanopore data, along with all proteins exhibiting expression level changes, identified by proteomics data, was analyzed using Cytoscape and R. Specifically, the analysis utilized RCy3, a package within R to connect to Cytoscape to directly visualize, analyze, and explore networks.

#### mRNA extraction

RNA extraction from BEAS-2B cells was performed by lysing cells with TRIzol Reagent (Invitrogen, 15596026). The RNA underwent DNase I treatment and was purified using a Direct-zol RNA Miniprep Kit (Zymo Research, R2052) according to the manufacturer’s protocol. The purity of the RNA was confirmed using a Nanodrop 2000 Spectrophotometer (Thermo Scientific), and RNA concentration was determined using a Qubit™ 4 Fluorometer (ThermoFisher) RNA Broad Range Assay Kit (ThermoFisher, Q10210). RNA quality was determined using an Agilent Bioanalyzer, and all RNA samples used for further analysis had RIN scores of 10. Two rounds of mRNA selection were performed using NEBNext Poly(A) mRNA Magnetic Isolation Module (NEB, E7490L). The eluted mRNA was used for downstream sequencing analyses.

#### BID-seq

The BID-seq protocol was performed as previously described with slight variations^55^. Briefly, mRNA was recovered as described above, and 100 ng of mRNA was used as input for each of 3 biological replicates for both the exposed and unexposed conditions. Input RNA was fragmented using RNA Fragmentation Reagents (Invitrogen) and purified with RNA Clean & Concentrator-5 Kit (Zymo). Fragmented RNA was then end-repaired using T4 Polynucleotide Kinase (NEB) and purified again. The 3’ and 5’ sequencing adapter ligation steps were carried out exactly as previously described, and the RNA was purified with the RNA Clean & Concentrator-5 Kit and eluted in 10 μL RNase-free H_2_O. Then, 1.5 μL of each sample was aliquoted as the “input” library. The bisulfite treatment was performed by mixing the remaining 8.5 μL of the RNA with an aqueous solution of 2.4 M Na_2_SO_3_ and 0.36M NaHSO_3_ for 3 h at 70°C. The treated RNA was then incubated with RNA Desulphonation Buffer (Zymo), purified with the RNA Clean & Concentrator-5 Kit, and eluted in 10 μL RNase-free H_2_O.

Reverse transcription was then performed on both the treated and input samples using SuperScript IV Reverse Transcriptase (Invitrogen). The cDNA was treated with RNase H, and PCR was performed to generate libraries for sequencing as previously described using NEBNext Multiplex Oligos for Illumina (Index Primers Set 1). The libraries were purified on a 3.5% (w/v) low-melting-point agarose gel with pBR322 DNA-MspI Digest (NEB) as a size marker. The libraries were excised from the gel between 200 and 400 bp and purified using the MinElute Gel Extraction Kit (Qiagen).

The libraries were then submitted to the University of Texas Genomic Sequencing and Analysis Facility for sequencing. The final libraries were checked for size and quality with the Bioanalyzer High Sensitivity DNA Kit (Agilent), and their concentrations were measured using the KAPA SYBR Fast qPCR kit (Roche). The pooled libraries were sequenced on a NovaSeq 6000 S1 Flow Cell (Illumina), and a sequencing depth of ∼140 million reads per sample was achieved with paired-end, 2 x 150bp read length.

#### BID-seq data analysis

The Docker image containing the BID-seq analysis pipeline v 2.0 (https://github.com/y9c/pseudoU-BIDseq) was accessed through Apptainer and run on the Texas Advanced Computing Center Lonestar 6 system. The PM exposed and unexposed samples were run separately using identical parameters. Before running the pipeline, the Human Reference Genome was assembled and indexed using the *Homo sapiens* GRCh38 DNA primary assembly fasta file and Homo_sapiens.GRCh38.110.gtf from Ensembl using the genomeGenerate run mode in STAR (39). The reference fasta files for contamination and small noncoding RNAs were taken from the Chaun He group’s GitHub (https://github.com/chelab/db/tree/main/reference_sequence). The YAML command files were formatted to accommodate the paired-end sequencing reads, as specified in the pipeline vignette. From the raw output of the pipeline, we filtered the putative Ψ sites using the following parameters: 1) Deletion rate must be above 5% in bisulfite-treated samples (with a deletion count above five). 2) Deletion rate must be below 3% in input libraries. 3) Read coverage depth must be above 20 for both input and bisulfite-treated libraries. 4) Deletion rate must be 1.5-fold over background for any given sequence motif.

#### Differential gene expression analysis

Differential gene expression data were computed using the output files from the BID-seq pipeline described above. The aligned reads in each.bam file output from STAR for the input (non-bisulfite-treated) samples of both the unexposed and exposed conditions were counted using HTSeq^71^. The DESeq2 package was then used to quantify differential gene expression in R^72^. Differential expression was determined relative to the counts from unexposed control cells. Significantly differentially expressed genes were defined as having a log_2_(Fold Change) ≥1 and padj<0.05, where the adjusted p-value is determined using the Benjamini-Hochberg correction.

*U*

#### NUAR motif enrichment analysis

All 5-mer sequence frequencies within every human RNA transcript were computed by running the k-mer extraction module of MathFeature^73^ against the complete human cDNA. fasta file (Homo_sapiens.GRCh38.cdna.all.fa). The resultant frequencies of all 5-mers belonging to the UNUAR motif (8 total) for each transcript were combined to generate a UNUAR frequency-per-transcript statistic. Transcripts corresponding to a specified GO term were determined using the BioMart Ensembl database^74^. The UNUAR frequencies of these corresponding transcripts were compared against a background of the entire human transcriptome to evaluate enrichment of the UNUAR motif within these GO term pathways.

### Statistical analysis

All experiments were performed in multiple, independent experiments, as indicated in the figure legends. All statistics and tests are described fully in the text or figure legend.

### Inclusion & Ethics Statement

All experiments involving human cells were conducted in accordance with institutional guidelines and approved protocols.

## DATA AVAILABILITY

Source Data are provided with this paper. Sequences were aligned to the yeast genome version S288C_R64-4-1. All FASTQ files generated in this work have been made publicly available in NIH NCBI SRA under Bioproject PRJNA1106629 (Human data), and PRJNA1112448 (Yeast data) Proteomics data is available for download from the PRIDE repository under Project accession: PXD052537 (Human data), PXD052684 (Yeast data) and MassIVE MSV000098950.

## CODE AVAILABILITY

Pseudouridine modification detection code can be found in https://github.com/RouhanifardLab. BID-seq analysis pipeline v 2.0 can be found in https://github.com/y9c/pseudoU-BIDseq. Open-source software from Oxford Nanopore Technologies is available on GitHub (https://github.com/nanoporetech/).

## ACKNOWLEDGEMENTS

We thank the following funding sources for their support: National Science Foundation (NSF CAREER 2045562 to K.S.K.), National Institute of Health (R35 GM128836 to K.S.K., K22CA262349 to C.W., R01HG011087 to Meni Wannunu, and R01HG012856 to S.H.R.), and American Heart Association (24PRE1194152 to M.R.). The authors acknowledge the funding support of this study from the National Institutes of Health R21ES032124 (L.M.C.), the Welch Foundation F-1756 (L.M.C.), and Hypothesis Fund Award (L.M.C.). This work was supported by the National Science Foundation Graduate Research Fellowship Program under Grant Nos. DGE 2137420 (S.M.E.) and DGE 2137420 (M.R.B). The authors acknowledge the Texas Advanced Computing Center (TACC) at UT Austin for providing high-performance computing resources. The authors also thank the Genomic Sequencing and Analysis Core Facility at UT Austin, RRID: SCR_021713, for assistance with RNA-sequencing, the Center for Biomedical Research Support Biological Mass Spectrometry Facility at UT Austin, RRID: SCR_021728, for assistance with human mass spectrometry proteomics, and the Microscopy Core at the University of Michigan for assistance with yeast imaging.

## AUTHOR CONTRIBUTIONS

M.R. designed and performed most of the experiments related to yeast cells (including plasmid construction, preparation, and analysis of Nanopore and proteomic samples, cellular ROS level detection, growth curves/MICs, competitive fitness assays, CLAP, and imaging under stress conditions) and prepared the initial draft of the manuscript. S.M.E., M.R.B., and L.M.C. carried out human cell experiments, including exposures, BID-seq, and fluorescence microscopy; drafted the human cell portion of the manuscript; and conducted proteomics experiments on human samples. X.L. performed tRNA extraction, tRNA Nanopore sequencing, data analysis, and contributed to CLAP experiments. R.S. and D.E.E. generated the PUS7-D256A construct, performed RNA extraction, and carried out stress tolerance assays. T.T., C.P., D.B., O.F., S.A., S.H.R., and M.J. ran Nanopore experiments with yeast mRNA samples and guided data analysis. C.W. contributed intellectually during monthly meetings and collaborated on PUS7 site selection. K.S.K. designed and supervised the project and wrote the manuscript. All the authors reviewed and approved the final version of the manuscript.

## EXTENDED DATA FIGURES

**Extended Data Fig. 1.**
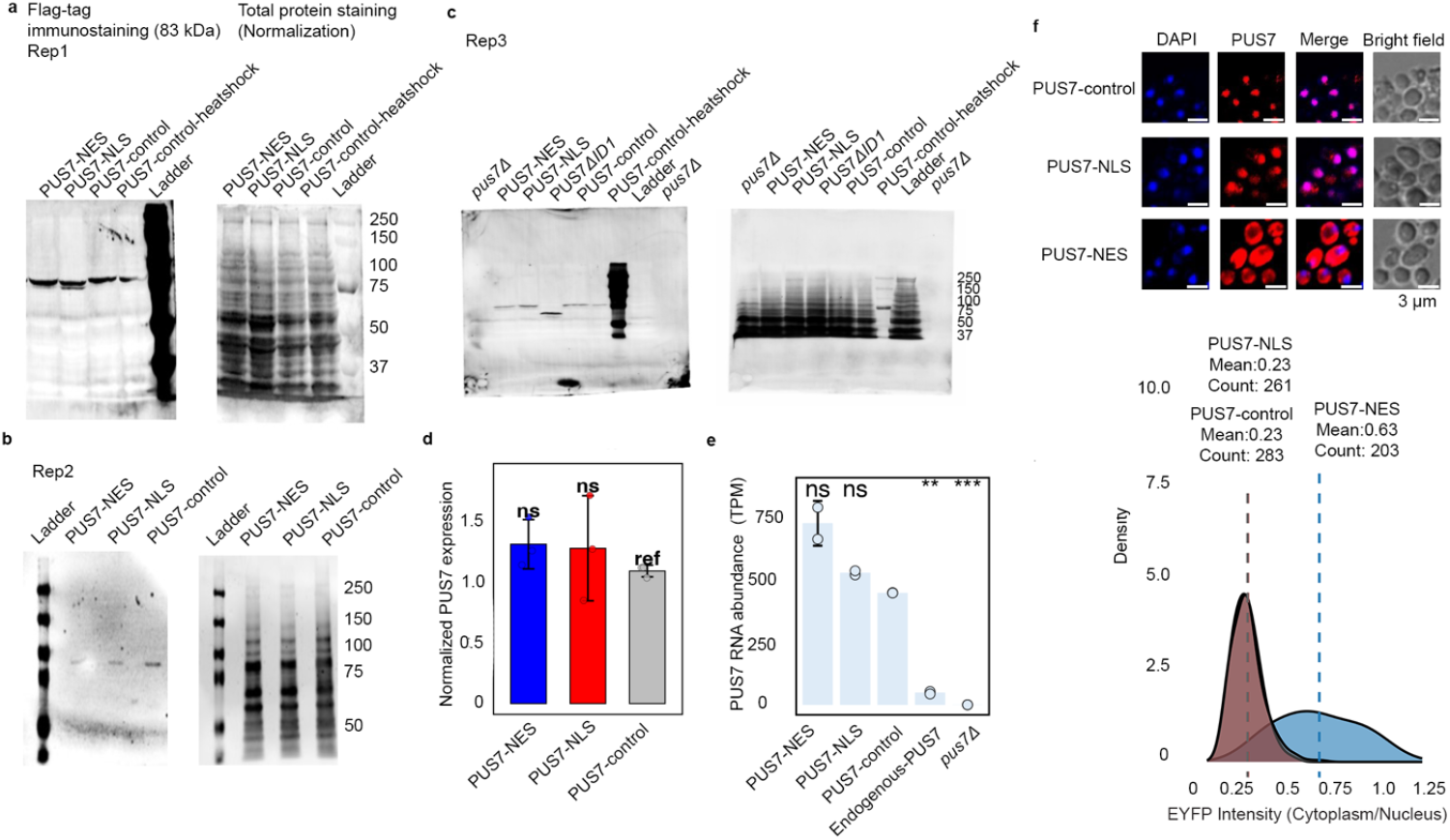
PUS7 localization induced Ψ sites are not PUS7 RNA abundance or protein level dependent. **a-d**, Western blot (triplicates) and quantification of PUS7 (anti-FLAG) in different localized yeast strains. Student’s t-test (unpaired, two-tailed), mean ± SD is depicted. (n = 3). **e**, PUS7 TPMs (Transcripts Per Kilobase Million) of differently localized PUS7, endogenous PUS7, and *pus7*Δ. Welch’s two-sample t-test (unpaired, two-sided), mean ± SD is depicted. *P < 0.05, **P < 0.01, ***P < 0.001 (n = 2). **f**, Immunofluorescence of anti-FLAG showing localization of PUS7-control, -NLS and -NES fused PUS7 and quantification of PUS7 localization with cellpose. Scale bar: 3 µm. (n >100 cells).

**Extended Data Fig. 2.**
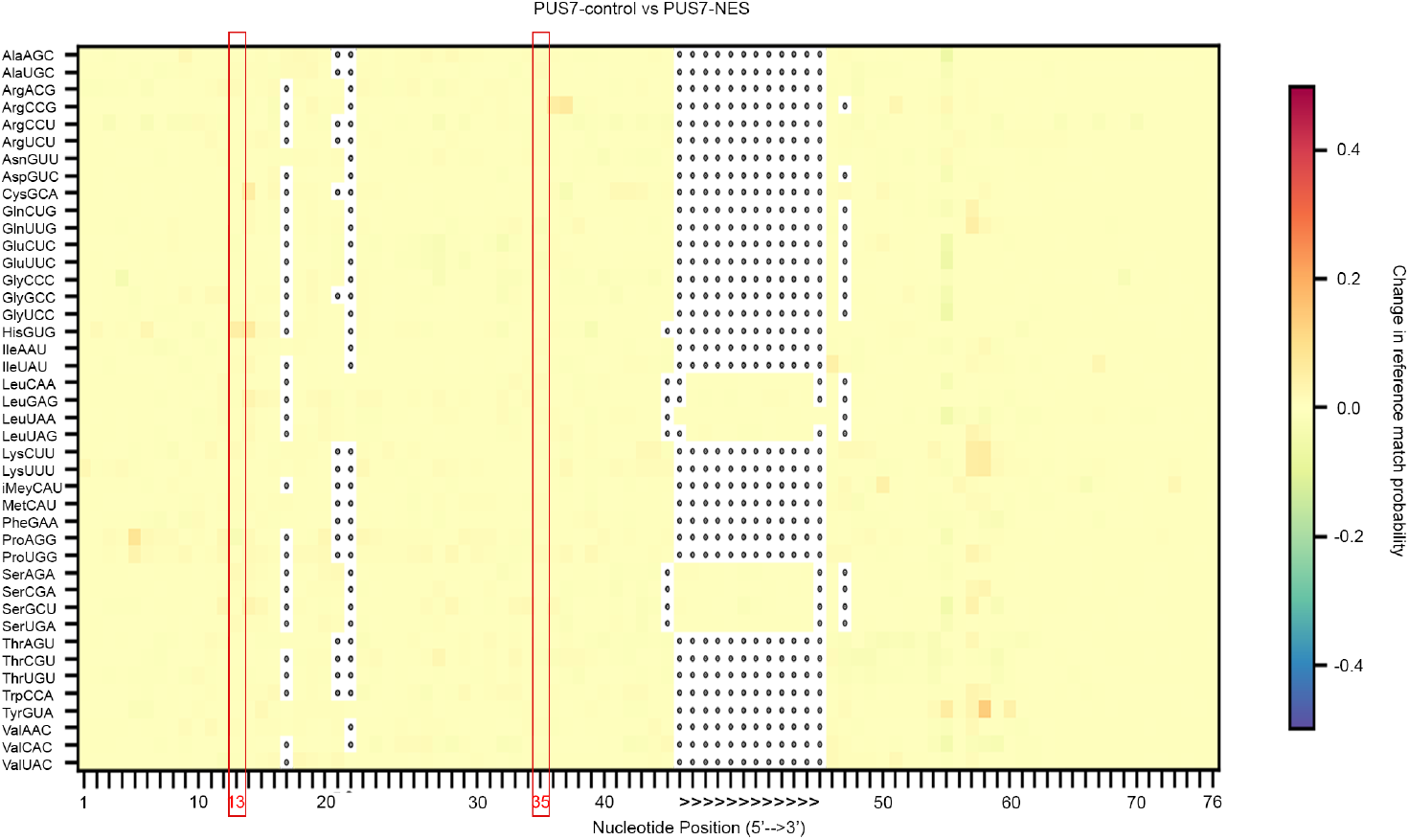
PUS7 localization change between PUS7-control and PUS7-NES does not impact Ψ levels in tRNAs. Ψ13 and Ψ35 modifications appear unaltered by the changes in PUS7 localization, highlighted in red boxes as Ψ13 and Ψ35. (n = 2).

**Extended Data Fig. 3.**
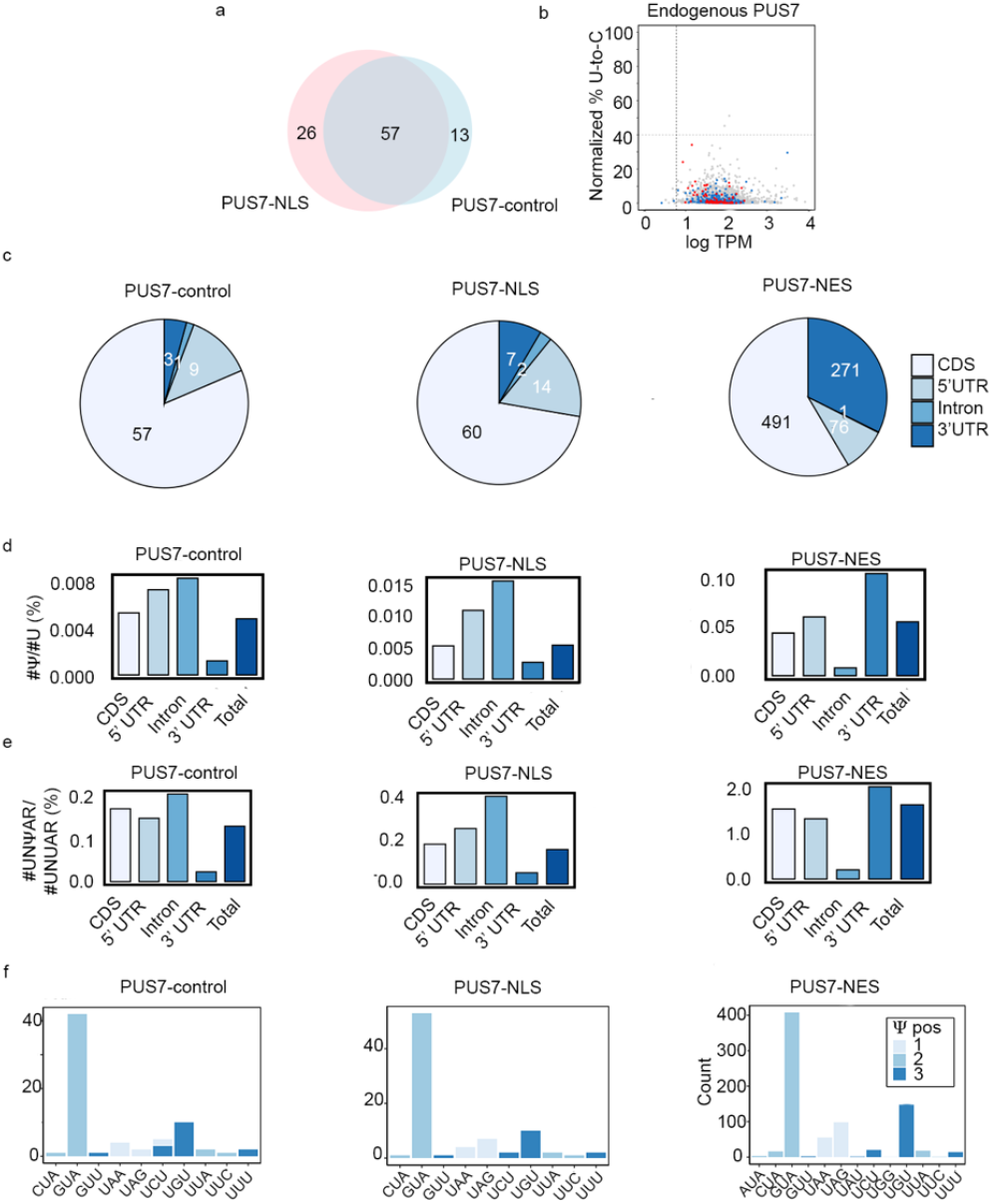
mRNA Ψ sites distribution for different localized PUS7. **a**, Venn diagram showing the overlap of high-confidence Ψ-modified mRNAs between PUS7-control *vs* PUS7-NLS. **b**, Scatter plot showing the number of total direct reads *vs* the relative occupancy of potential Ψ (% U-to-C mismatch) for endogenous PUS7. **c**, The number of high-confidence Ψ sites distribution in CDS, 5’-UTRs, intron, and 3’-UTR for PUS7-control, PUS7-NLS, PUS7-NES, respectively. **d-e**, ratio of #Ψ/#U and #UNΨAR/#UNUAR in different regions for PUS7-control, PUS7-NLS, PUS7-NES, respectively. **f**, Codon distribution of Ψ in PUS7-control, PUS7-NLS, PUS7-NES.

**Extended Data Fig. 4.**
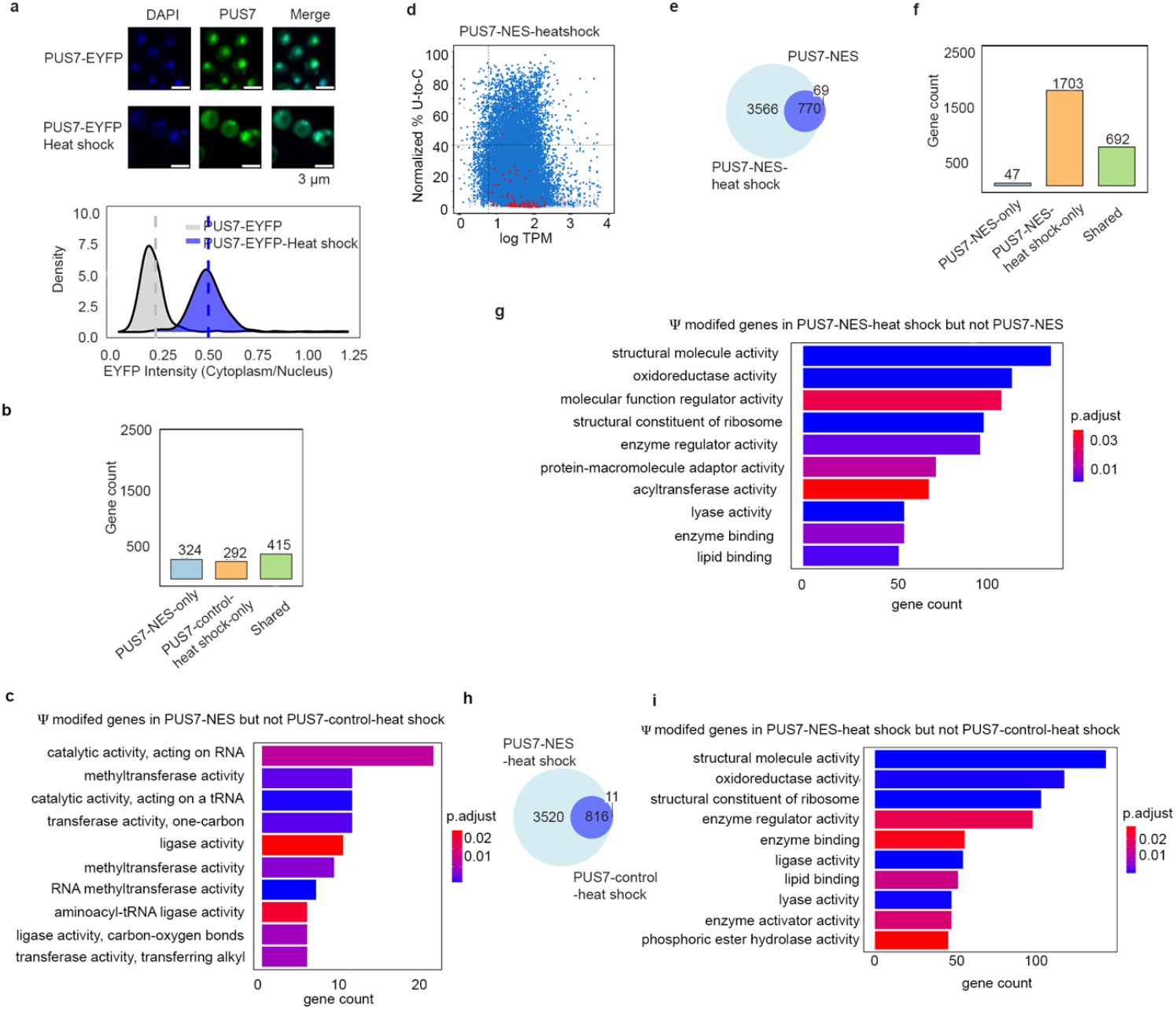
Heat shock-related PUS7 localization shift and Ψ site enrichment. **a**, Imaging and quantification of PUS7 localization of PUS7-EYFP cells under heat shock (45 °C, 1 h). Scale bar: 3 µm. **b**, overlap and unique of mRNAs modified between PUS7-NES *vs* PUS7-control-heat shock (45°C, 15min). **c**, Top10 GO enrichment analysis of Ψ modified mRNAs in PUS7-NES but not PUS7-control-heat shock (45°C, 15min). **d**, Scatter plots showing the number of total direct reads versus the relative occupancy of potential Ψ (% U-to-C mismatch) for PUS7-NES-heat shock. **e**, Venn diagram showing the overlap of Ψ sites between PUS7-NES-heat shock *vs* PUS7-NES. **f**, overlap and unique of mRNAs modified between PUS7-NES-heat shock (45°C, 15min) *vs* PUS7-NES. **g**, Top10 GO enrichment analysis of Ψ modified mRNAs in PUS7-NES-heat shock (45°C, 15min) but not PUS7-NES. **h**, Venn diagram showing the high-confidence Ψ sites modified in PUS7-NES-heat shock *vs* PUS7-control-heat shock (45 ^°^C, 15min). **i**, Top10 GO enrichment analysis of Ψ modified mRNAs in PUS7-NES-heat shock but not PUS7-control-heat shock (45°C, 15min).

**Extended Data Fig. 5.**
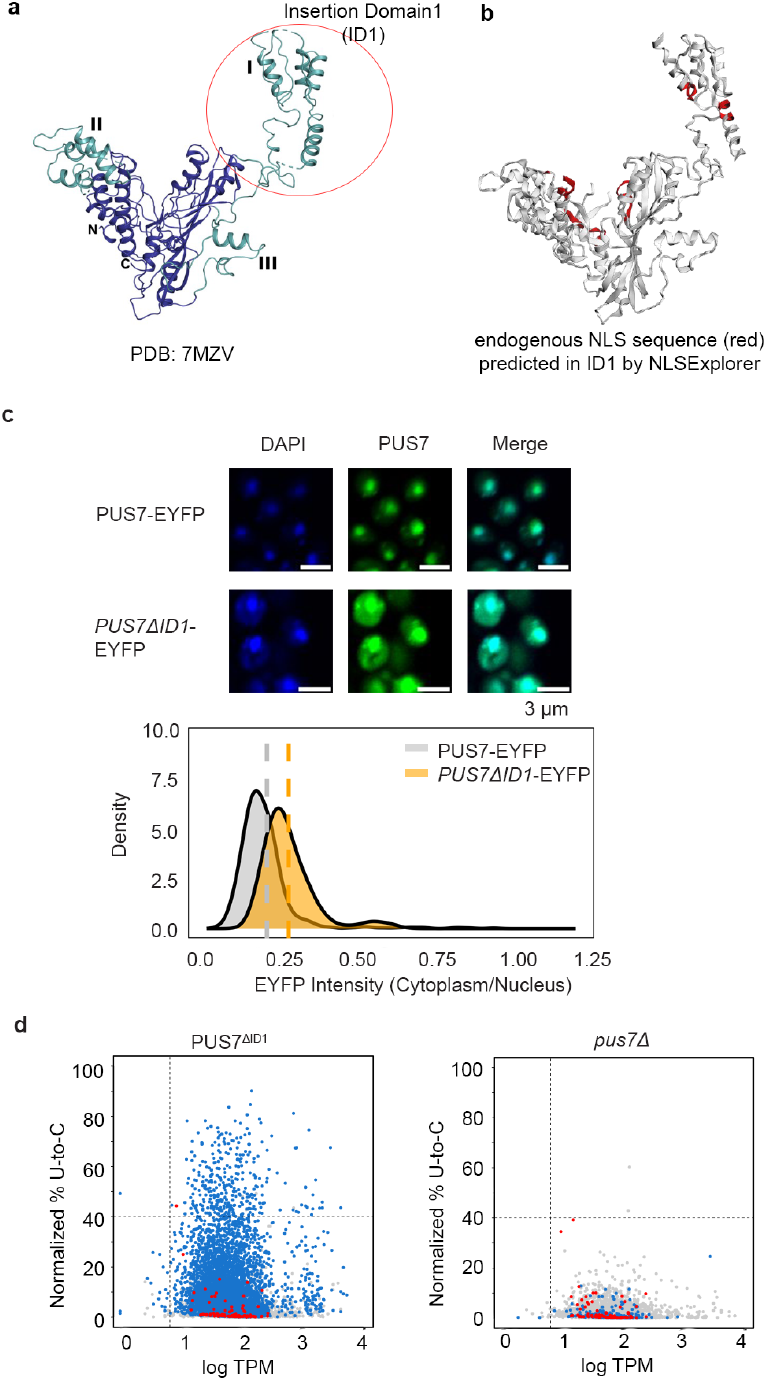
PUS7^ΔID1^-related PUS7 localization shift and Ψ site enrichment. **a**, Yeast PUS7 structure (PDB: 7MZV), with the ID1 region highlighted. **b**, Predicted nuclear transport region shown in red, as predicted by NLSExplorer. **c**, Imaging and quantification of PUS7 localization of PUS7^ΔID1^-EYFP cells, n =107. Scale bar: 3 µm. **d**, Scatter plots showing the number of total direct reads versus the relative occupancy of potential Ψ (% U-to-C mismatch) for PUS7^ΔID1^ and *pus7*Δ. Dot colors indicate 5-mer motifs recognized by PUS7 (blue) and TRUB1 (red).

**Extended Data Fig. 6.**
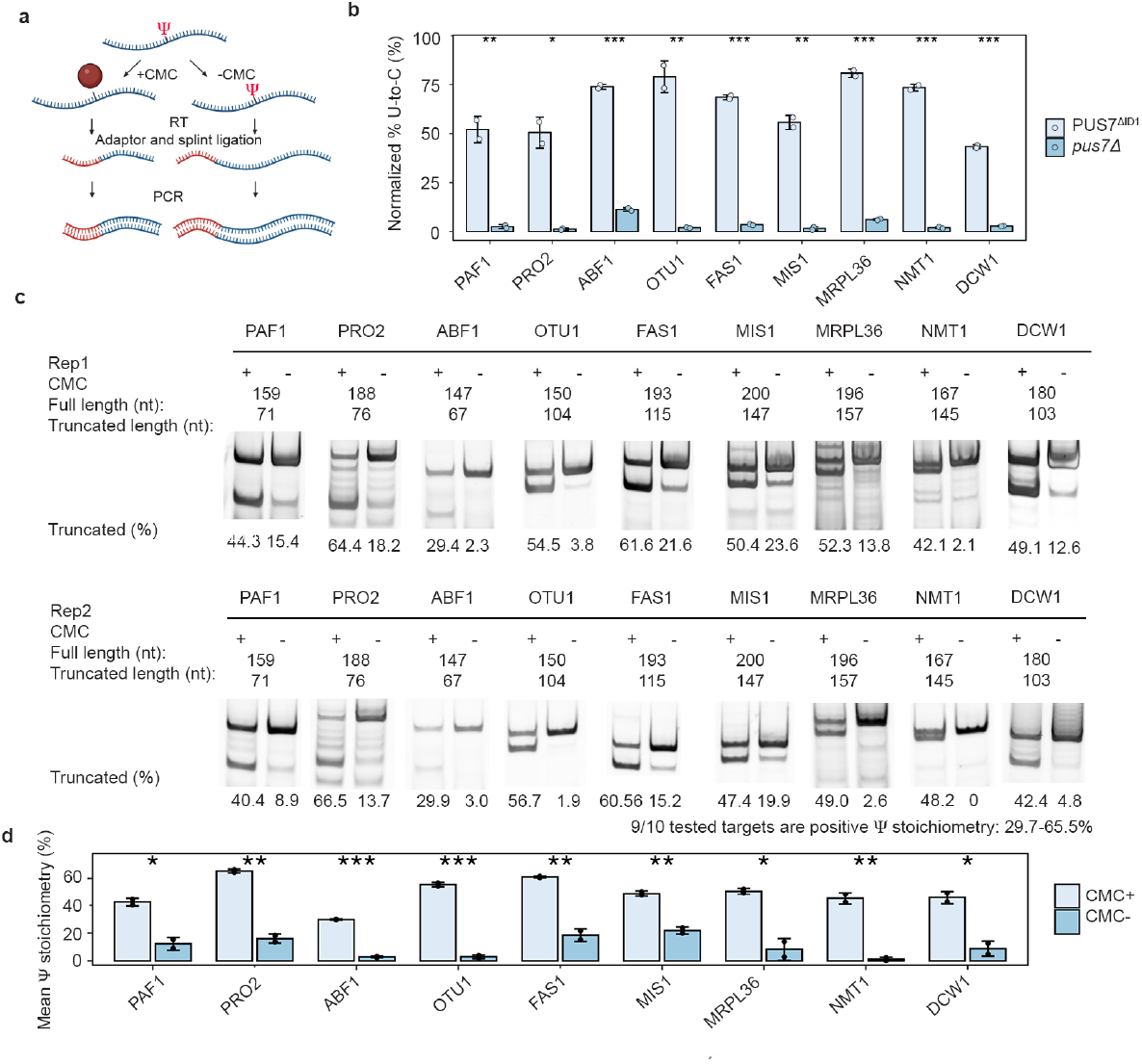
CLAP results and quantification. **a**, The diagram of <u>C</u>MC-RT and ligation <u>a</u>ssisted <u>P</u>CR analysis of Ψ–modification (CLAP) experiment. **b**, Nanopore sequencing detected U to C level (%) for arbitrarily selected targets. Student’s t-test (unpaired, two-tailed), mean ± SD is depicted. (n = 2). **c-d**, Gel result and quantification of Ψ incorporation in mRNAs in CLAP experiment between PUS7^ΔID1^ and *pus7*Δ. For gel source data, see **Supplementary Fig.8**. Student’s t-test (unpaired, two-tailed), mean ± SD is depicted. \**P* < 0.05, \*\**P* < 0.01, \*\*\**P* < 0.001 (*n* = 2).

**Extended Data Fig. 7.**
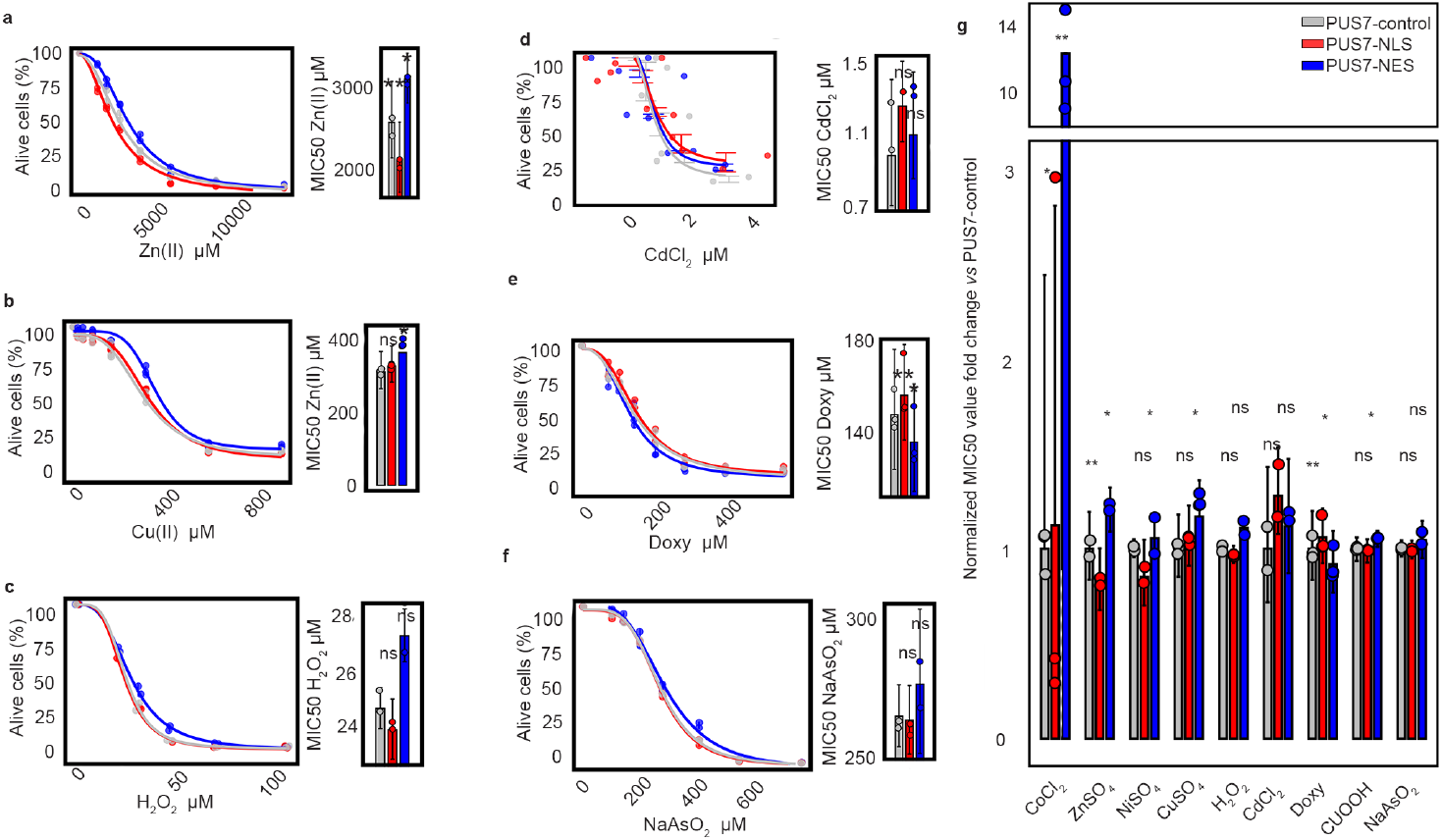
MIC50 result for different localized PUS7 yeast under various stress conditions. **a-f**, MIC50 result for yeast with PUS7-NES compared to yeast with PUS7-NLS and PUS7-control. Paired t test (two tailed), mean ± SD is depicted; p-value 0.0126 for NES-zinc, p-value 0.0053 for NLS-zinc (**a**) (n = 3); p-value 0.0168 for NES-copper, p-value 0.486 for NLS-copper (**b**) (n = 3); p-value 0.1025 for NES-H_2_O_2_, p-value 0.1125 for NLS-H_2_O_2_ (**c**) (n = 2); p-value 0.3452 for NES-CdCl_2_, p-value 0.7156 for NLS-CdCl_2_ (**d**) (n = 2); p-value 0.0242 for NES-Doxycycline, p-value 0.0056 for NLS-Doxycycline (**e**) (n = 3); p-value 0.407 for NES-NaAsO_2_, p-value 0.08 for NLS-NaAsO_2_ (**f**) (n = 2); **g**, Summary plot of the normalized MIC value change between PUS7-NES/NLS and PUS7-control. \**P* < 0.05, \*\**P* < 0.01, \*\*\**P* < 0.001. Paired t-test (two-tailed), mean ± SD is depicted.

**Extended Data Fig. 8.**
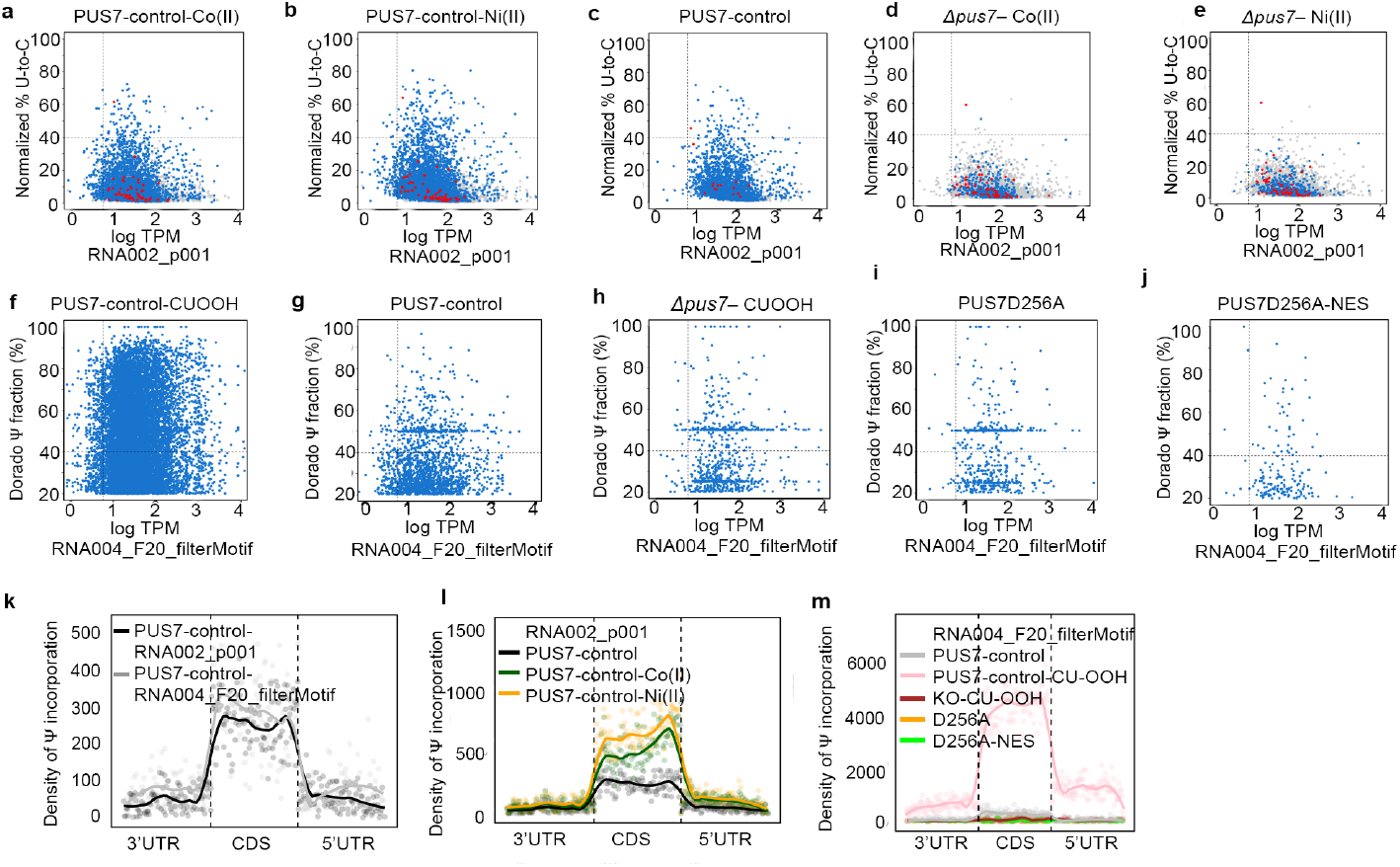
Ψ detection under Co(II), Ni(II), and CU. **a-e**, Scatter plots of total direct reads versus relative occupancy of potential Ψ (Mod-p ID, p < 0.001) for PUS7-control–Co(II) (30 μM, 4 h) (**a**), PUS7-control–Ni(II) (200 μM, 4 h) (**b**), PUS7-control (**c**), *pus7*Δ–Co(II) (30 μM, 4 h) (**d**), *pus7*Δ–Ni(II) (200 μM, 4 h) (**e**) (RNA002). **f-j**, Scatter plots of total direct reads versus relative occupancy of potential Ψ (Dorado Ψ incorporation fraction >20%, UNUAR motif filter) for PUS7-control–CU (2 mM, 1 h) (**f**), PUS7-control (**g**), *pus7*Δ–CU (2 mM, 1 h) (**h**), PUS7D256A (**i**), PUS7D256A–NES (**j**) (RNA004). Dot colors indicate 5-mer motifs recognized by PUS7 (blue) and TRUB1 (red). **k, l, m**, Density maps of Ψ (U to C mismatch and Dorado fraction) distribution across 5′ UTR, CDS, and 3′ UTR: **k**, PUS7-control (RNA002, Mod-p ID p < 0.001) *vs* PUS7-control (RNA004, Dorado Ψ fraction >20%). **l**, PUS7-control, PUS7-control–Co(II) (30 μM, 4 h), and PUS7-control–Ni(II) (200 μM, 4 h) (RNA002, Mod-p ID p < 0.001). **m**, PUS7-control, PUS7-control–CU (2 mM, 1 h), *pus7*Δ–CU (2 mM, 1 h), PUS7D256A, PUS7D256A-NES (RNA004, Dorado Ψ fraction >20%, UNUAR motif).

**Extended Data Fig. 9.**
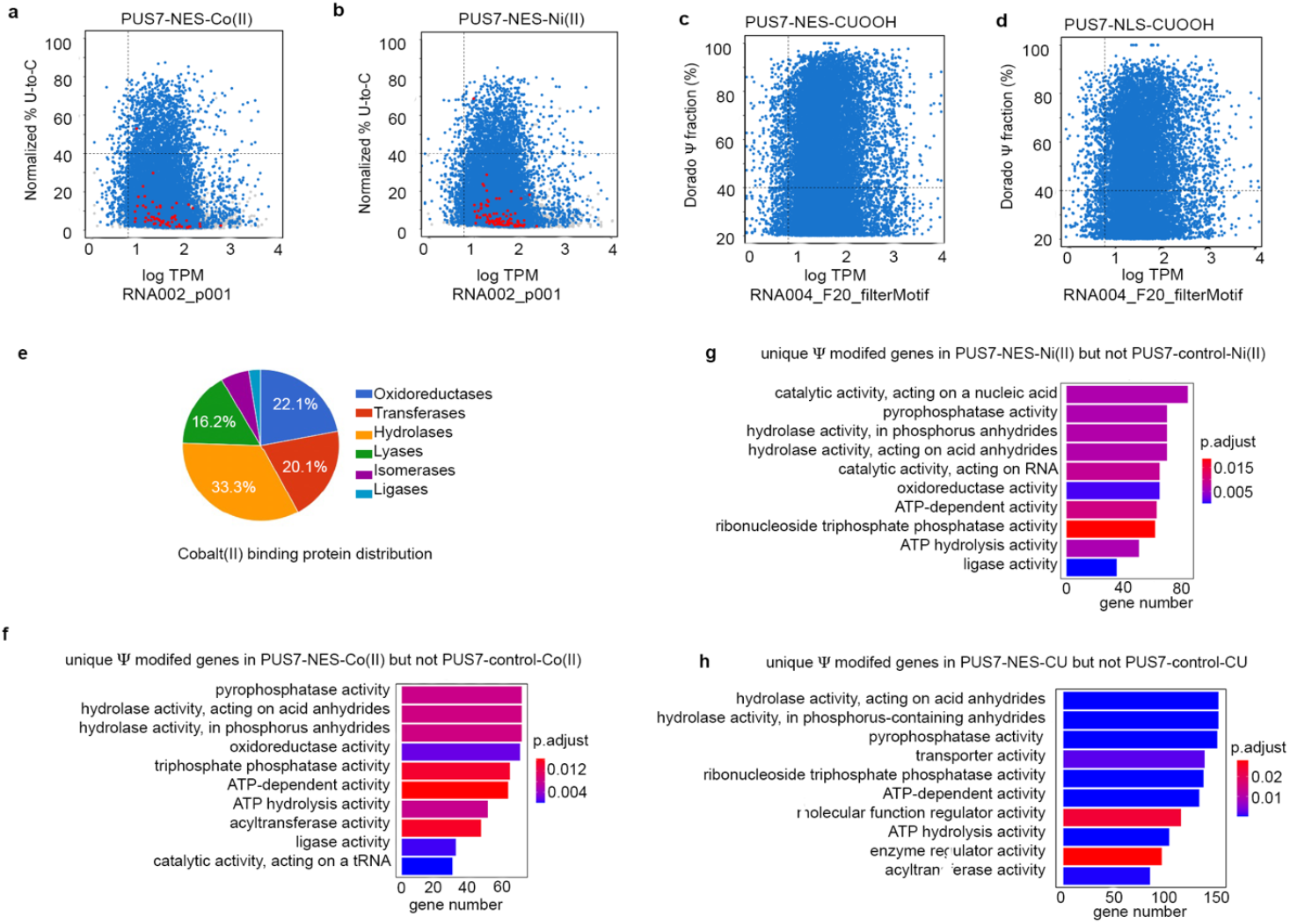
Ψ detection for PUS7-NES/-NLS under Co(II), Ni(II), and CU together with GO enrichment. **a-b**, Scatter plots of total direct reads versus relative occupancy of potential Ψ (Mod-p ID, p < 0.001) for PUS7-NES– Co(II) (30 μM, 4 h) (**a**), PUS7-NES–Ni(II) (200 μM, 4 h) (**b**) (RNA002). **c-d**, Scatter plots of total direct reads versus relative occupancy of potential Ψ (Dorado Ψ incorporation fraction >20%, UNUAR motif) for PUS7-NES–CU (2 mM, 1 h) (**c**) and PUS7-NLS–CU (2 mM, 1 h) (**d**) (RNA004). Dot colors indicate 5-mer motifs recognized by PUS7 (blue) and TRUB1 (red). **e**, Co(II) binding protein distribution in GO groups from MetalPDB, with the top 3 groups being oxidoreductase, transferase activity, and hydrolase activity. **f-h**, Top10 GO terms with significant *p*-values (< 0.05) for genes whose mRNAs are robustly modified by PUS7-NES under Co(II) (30 μM, 4 h) (**f**), Ni(II) (200 μM, 4 h) (**g**) and CU (2 mM, 1 h) (**h**) but not in PUS7-control under the same stresses.

**Extended Data Fig. 10.**
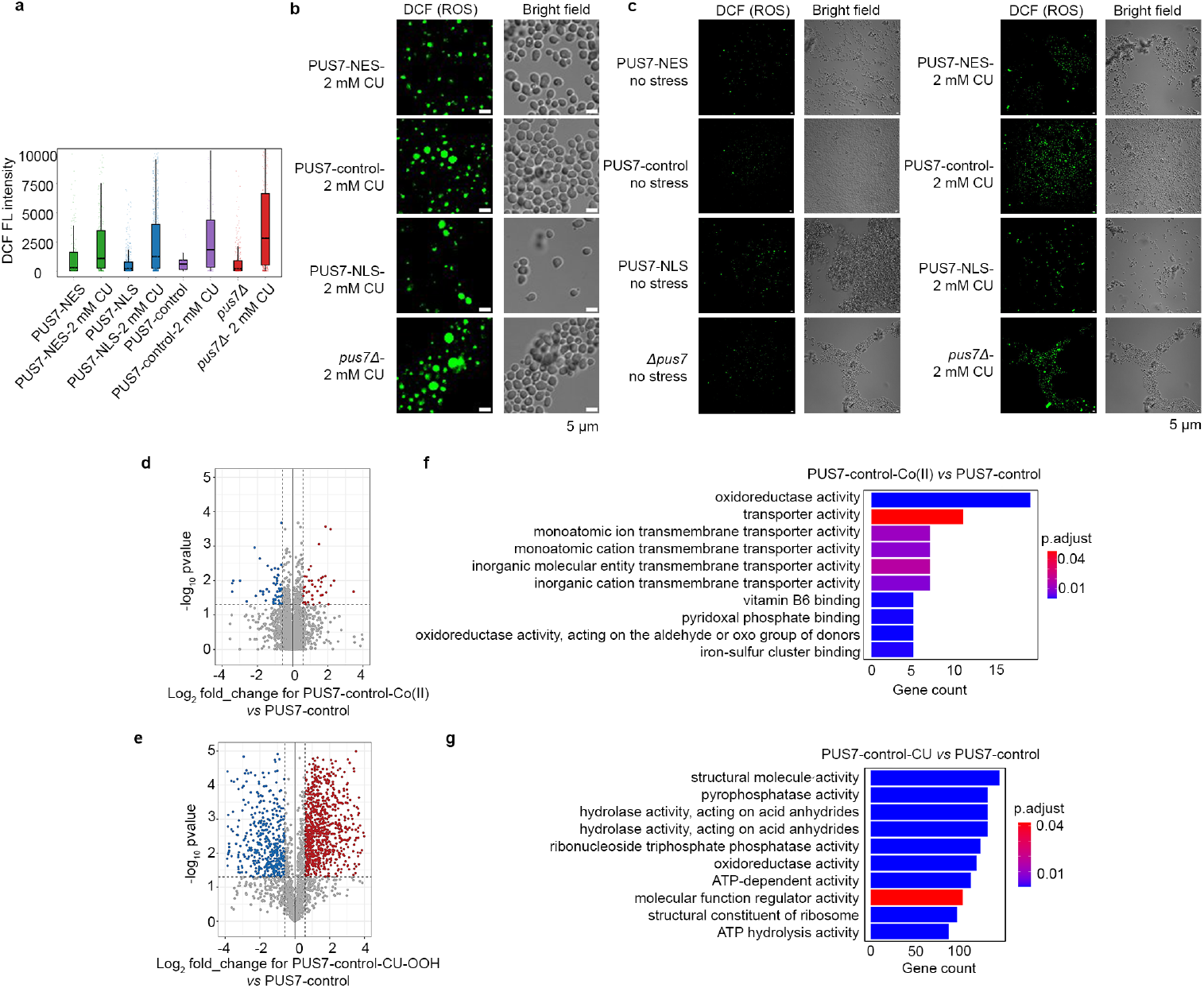
Cellular ROS level for different PUS7 localized cells under no stress condition and 2 mM CU stress for 30 min, and protein expression level change under Co(II) and CU. **a**, Quantification of cellular ROS level when cells are under no stress and 2 mM CU for 30 min, Box center line = median; box limits = 1st (Q1) and 3rd quartiles (Q3). **b-c**, Images for cells under no stress and 2 mM CU for 30 min. Scale bar: 5 µm.. **d-e**, Volcano plot showing yeast protein expression levels between PUS7-control cells under 30 µM cobalt stress for 4 h (**d**) or 2 mM CU for 1 h (**e**) *vs* under no stress conditions. Red dots: upregulation; blue dots: downregulation. Student’s two-sample t-test (unpaired, two-tailed). P<0.05 and |log_2_ fold-change| > log (1.5) as significant changes. (*n* = 3). **f-g**, Top 10 GO terms with significant p-values (< 0.05) for genes whose mRNAs are robustly modified by PUS7-control under Co(II) and CU but not in PUS7-control under no stress conditions.

**Extended Data Fig. 11.**
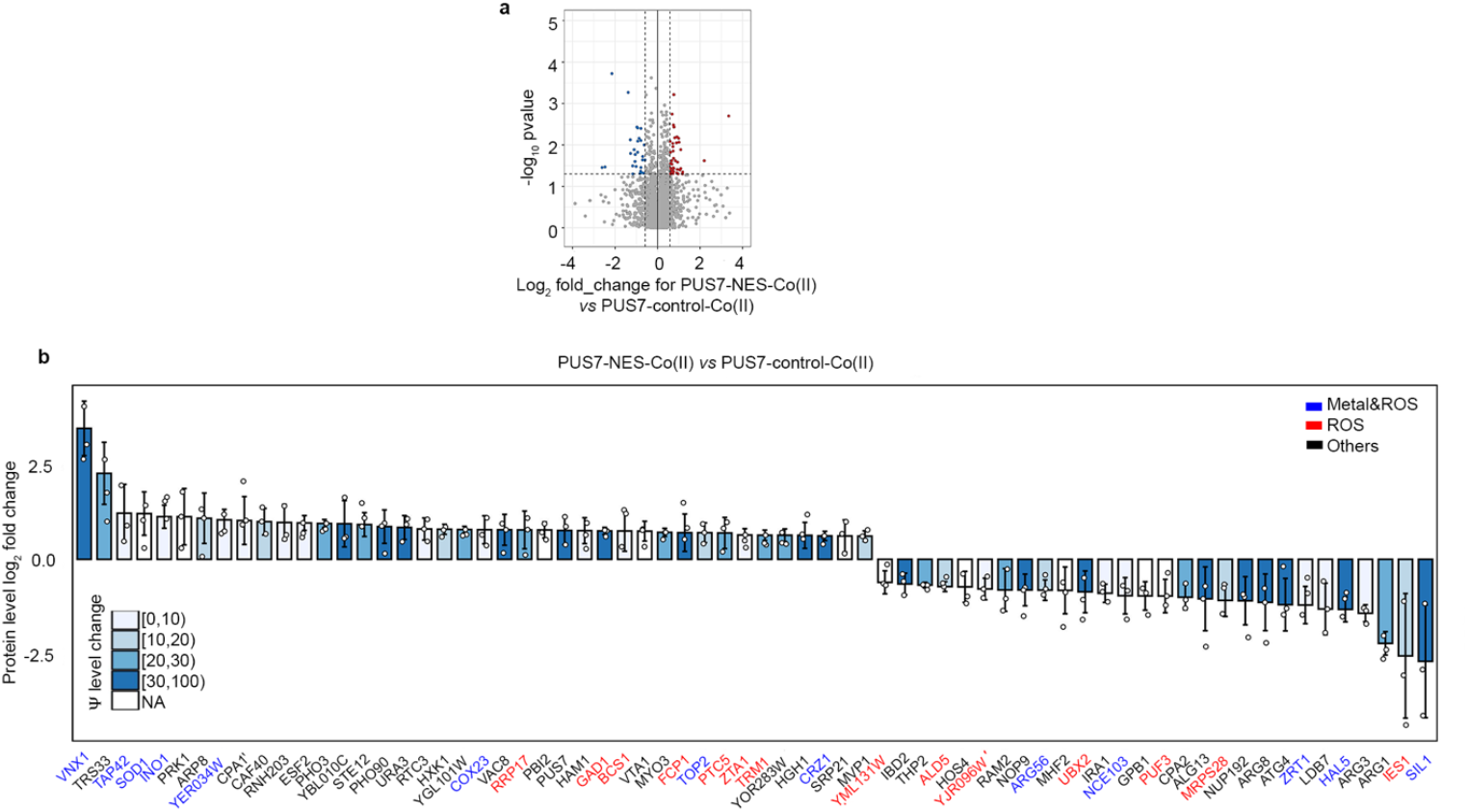
Protein expression level changing between PUS7-NES-Co(II) and PUS7-control-Co(II). **a**, Proteomics data showing yeast protein expression levels between PUS7-NES and PUS7-control cells under 30 µM cobalt stress for 4 h. Red dots: upregulation; blue dots: downregulation. Student’s two-sample t-test (unpaired, two-tailed). P<0.05 and |log_2_ fold-change| > log (1.5) as significant changes. (*n* = 3). **b**, Proteins showing significant expression level changes and their Ψ level changes (shown by bar color) between PUS7-NES and PUS7-control under cobalt 30 µM cobalt stress for 4 h. Metal-binding proteins in blue text label. ROS-related proteins in red text label. Others are shown in a black label. Student’s two-sample t-test (unpaired, two-tailed). P<0.05 and |log_2_ fold-change| > 1.5 as significant changes. Mean ± SD is depicted. (n = 3).

**Extended Data Fig. 12.**
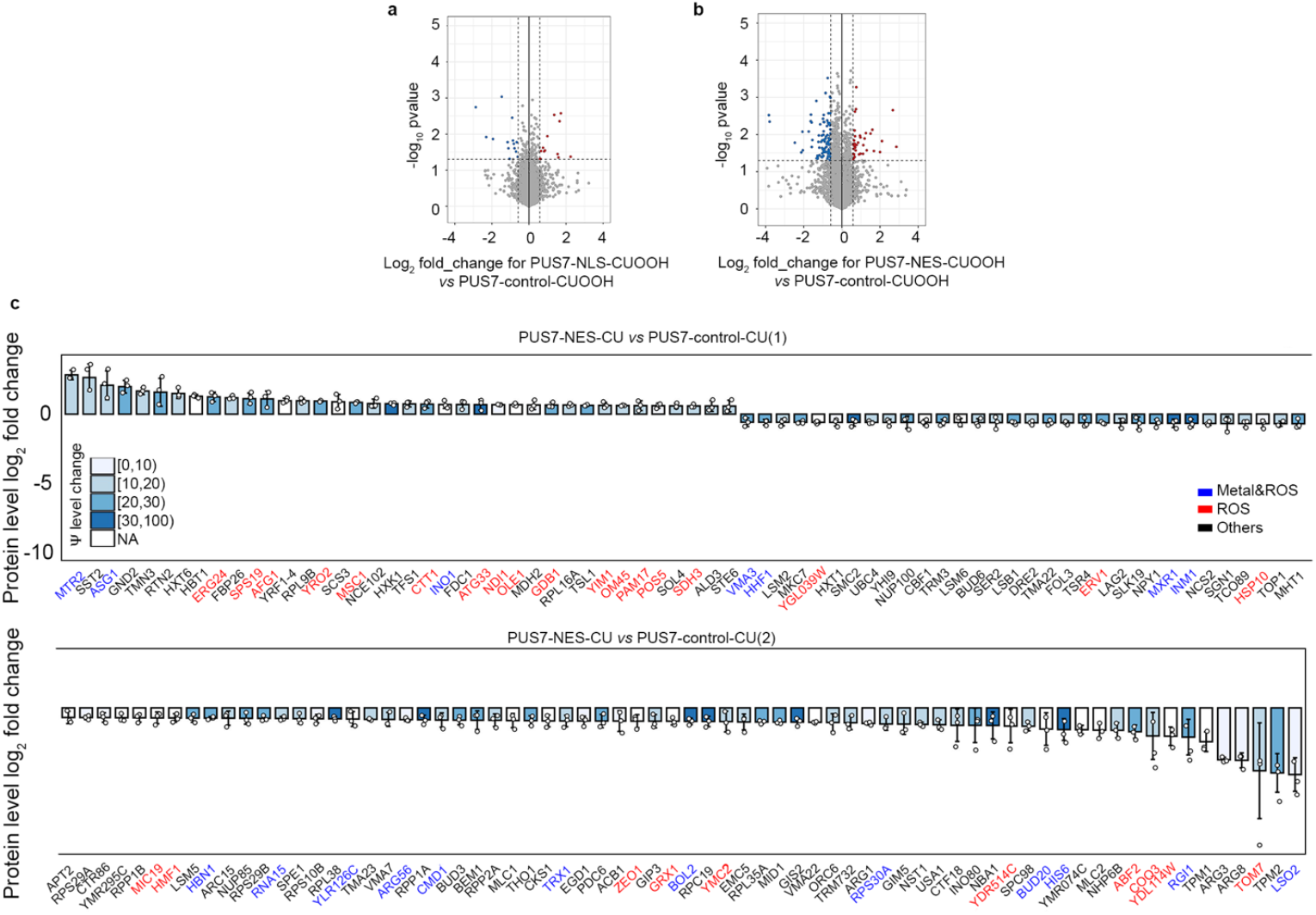
Protein expression level changing between PUS7-NES under CU and PUS7-control under CU. **a-b**, Proteomics data showing yeast protein expression level changes between PUS7-NLS under 2 mM CU for 1 h and PUS7-control cells under the same stress (**a**), PUS7-NES under 2 mM CU for 1 h and PUS7-control cells under the same stress (**b**). Red dots: upregulation; blue dots: downregulation. Student’s two-sample t-test (unpaired, two-tailed). P<0.05 and |log_2_fold-change| > log (1.5) as significant changes. (*n* = 3). **c**, Proteins showing significant expression level changes and their Ψ level changes (shown by bar color) between PUS7-NES under 2 mM CU for 1 h and PUS7-control under the same stress. Metal-binding proteins in blue text label. ROS-related proteins in red text label. Other proteins are in the black text label. Student’s two-sample t-test (unpaired, two-tailed). P<0.05 and |log_2_fold-change| > 1.5 as significant changes. Mean ± SD is depicted. (n = 3).

**Extended Data Fig. 13.**
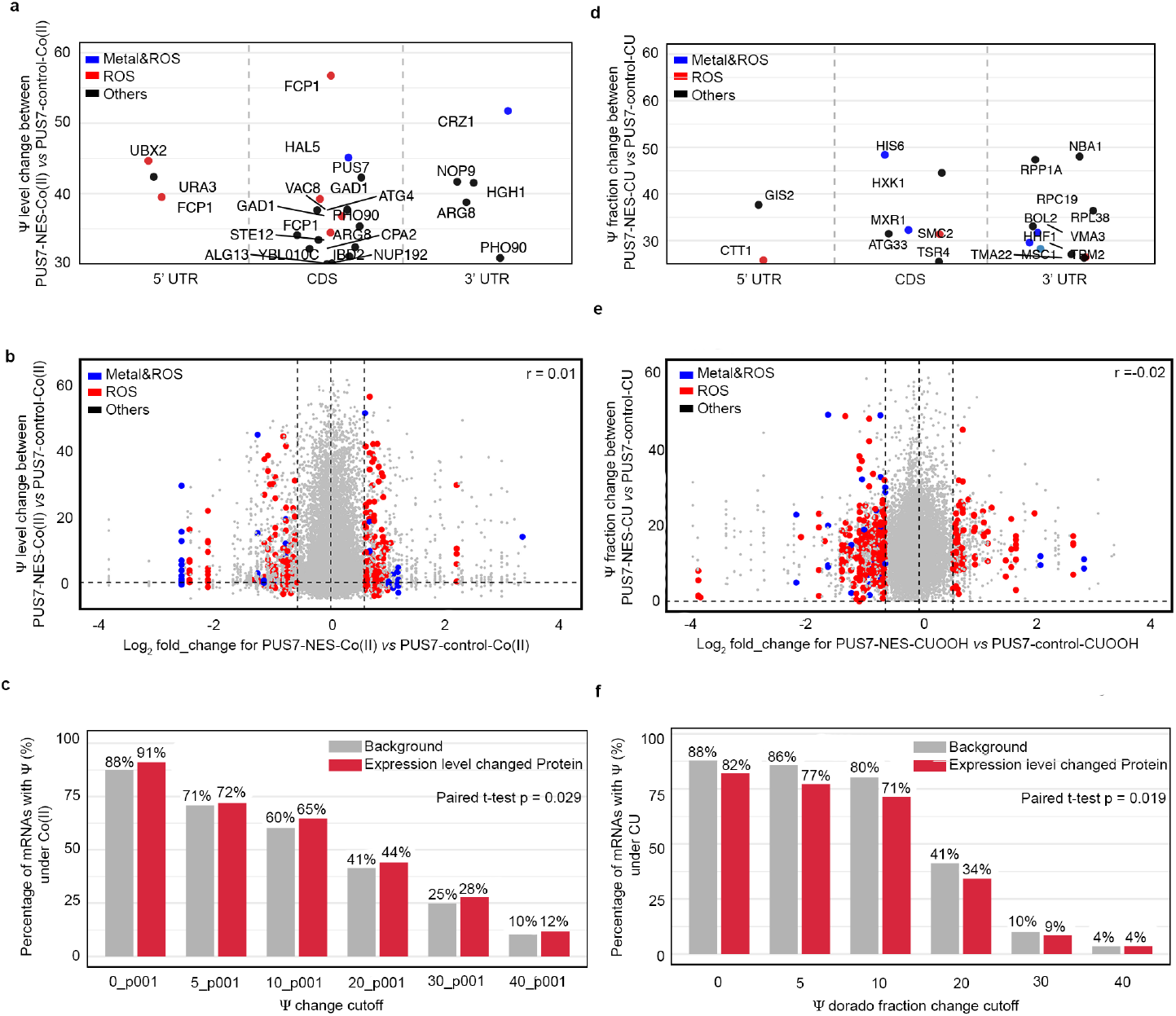
Correlation of proteins showing an expression level change and Ψ level change between PUS7-NES and PUS7-control under stresses. **a**,**d**, Ψ incorporated regions (3’ UTR, CDS, and 5’ UTR) of mRNAs with over 30% Ψ level increasing together with encoded proteins showing a significant expression level change between PUS7-NES under cobalt stress (30 µM cobalt stress for 4 h) (**a**) or CU (2 mM CU for 1 h) (**d**) and PUS7-control under the same stresses. Metal-binding proteins are labeled in blue. ROS-related proteins labeled in red. Others are labeled in black. **b**,**e**, Correlation between pseudouridylation level changes (PUS7-NES-Co(II) vs. PUS7-control-Co(II) (**b**); PUS7-NES-CU vs. PUS7-control-CU (**e**) and protein expression level changes. protein levels showing significant changes are bolded (Student’s two-sample t-test (unpaired, two-tailed), p-value < 0.05, |log_2_fold-change| > 1.5). (n = 3), Metal-binding related proteins with expression level changes are labeled in blue, ROS-related proteins are labeled in red. **c, f**, Percentage of detected proteins (with or without expression changes) that have Ψ-modification in the mRNAs between PUS7-NES and PUS7-control under 30 µM cobalt stress for 4 h (**c**) and 2 mM CU stress for 1 h (**f**), evaluated across different cutoffs.

**Extended Data Fig. 14.**
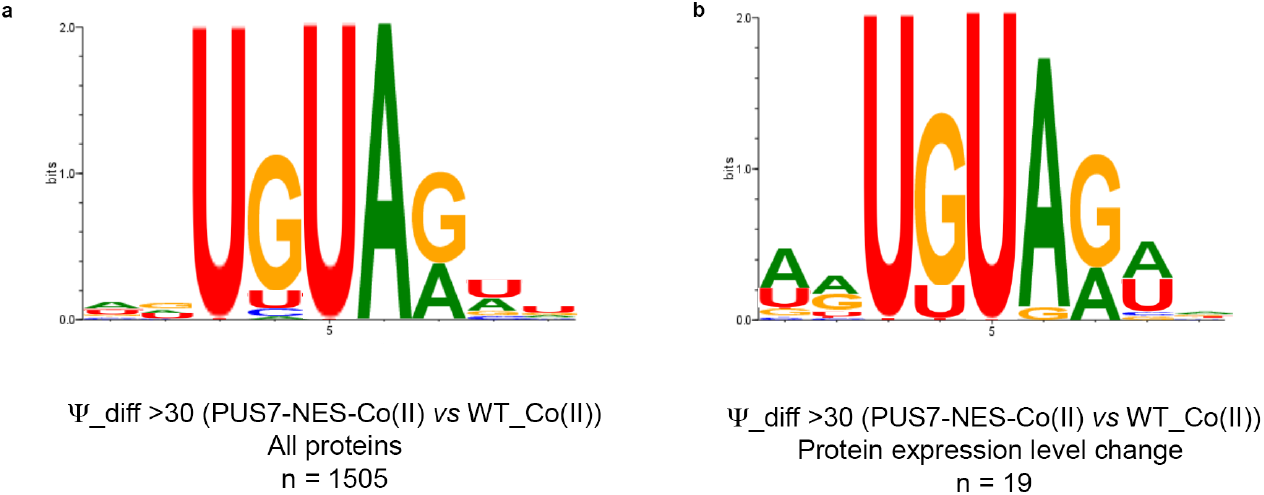
Sequence context of Ψ sites present in PUS7-NES–Co(II) (30 µM cobalt stress for 4 h) but absent in PUS7-control–Co(II) (30 µM cobalt stress for 4 h). **a**, and of detected Ψ sites from genes showing differential protein expression between PUS7-NES–Co(II) (30 µM cobalt stress for 4 h) and **b**, PUS7-control–Co(II) (30 µM cobalt stress for 4 h).

**Extended Data Fig. 15.**
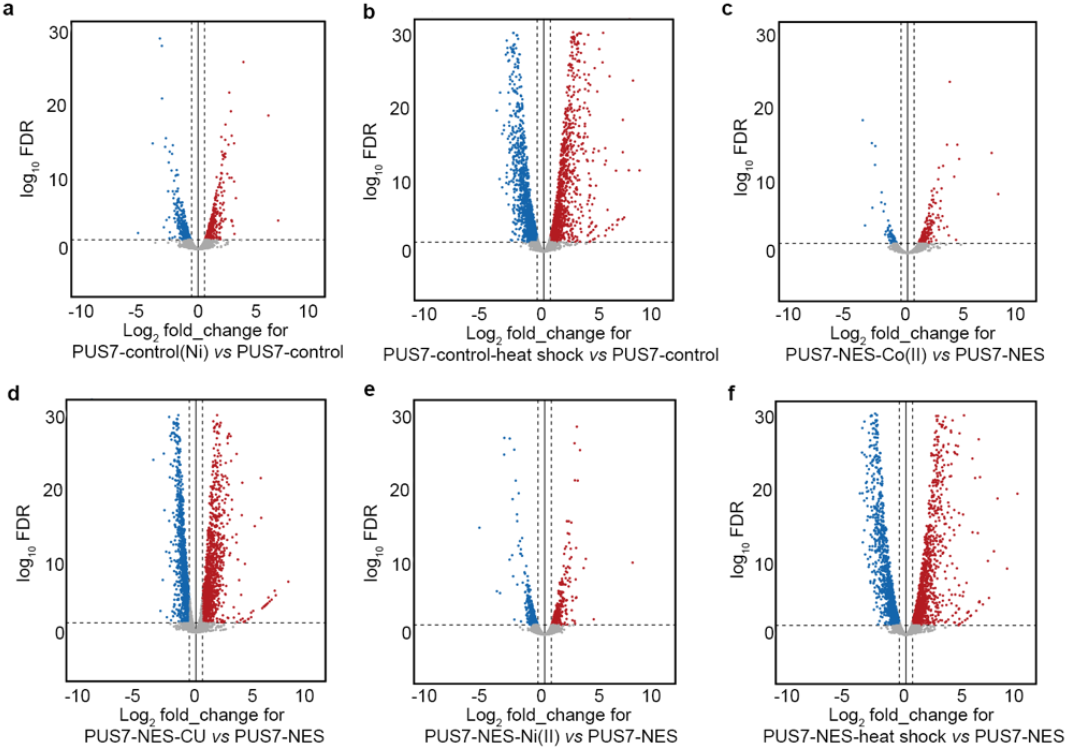
Stresses broadly change mRNA abundance. **a–b**, Volcano plots showing steady-state RNA level changes in PUS7-control cells subjected to Ni(II) (200 μM, 4 h) (**a**) or heat shock (45 °C, 15 min) (**b**), relative to unstressed conditions. **c–f**, Volcano plots showing steady-state RNA level changes in PUS7-NES cells subjected to Co(II) (30 μM, 4 h) (**c**), CU (2 mM, 1 h) (**d**), Ni(II) (200 μM, 4 h) (**e**), or heat shock (45 °C, 15 min) (**f**), relative to unstressed growth conditions (30 °C) Student’s two-sample t-test (unpaired, two-sided). Red dots: upregulation; blue dots: downregulation. (n = 2).

**Extended Data Fig. 16.**
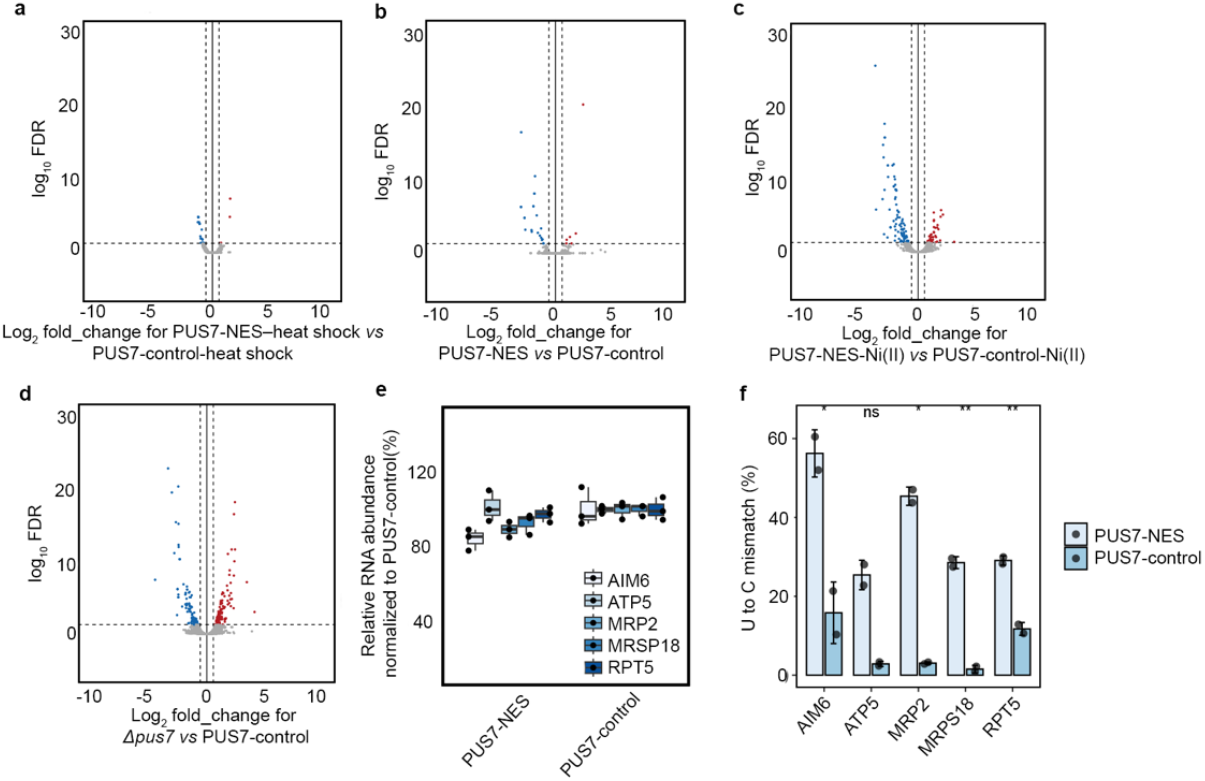
Ψ incorporation in mRNAs does not broadly change steady-state RNA levels. **a–c**, Volcano plots showing RNA abundance log_2_ fold changes between PUS7-NES and PUS7-control cells under heat shock (45 °C, 15 min) (**a**), no-stress conditions (**b**), and Ni(II) (200 μM, 4 h) (**c**). **d**, Volcano plot showing RNA abundance differences between *pus*Δ and PUS7-control yeast. Student’s two-sample t-test (unpaired, two-sided). Red dots: upregulation; blue dots: downregulation. (n = 2). **e**, RT–qPCR analysis of the mRNA abundance of five mRNAs exhibiting high Ψ incorporation in PUS7-NES but not PUS7-control cells under no-stress conditions. **f**, Nanopore sequencing–detected U-to-C levels (%) for selected RT-qPCR targets in PUS7-NES and PUS7-control cells under no-stress conditions. Student’s t-test (unpaired, two-tailed), mean ± SD is depicted. \**P* < 0.05, \*\**P* < 0.01 (*n* = 2).

**Extended Data Fig. 17.**
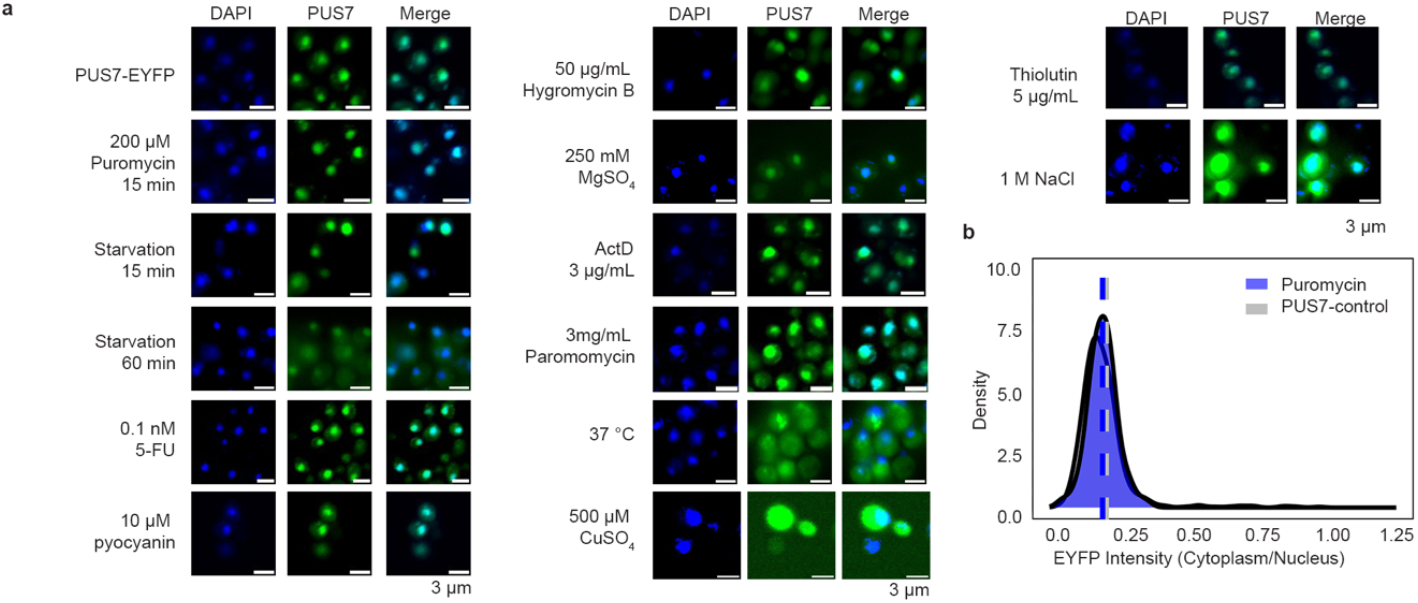
PUS7 localization under different stress conditions (including translation inhibitors and starvation status). **a**, Visualization of PUS7 localization (PUS7–EYFP) under standard growth conditions and various stress conditions. Stress treatments were applied for 60 min before fixation unless otherwise specified. Scale bar: 3 µm. **b**, Quantification of fluorescence intensity (FI) ratio changes in the cytoplasm versus nucleus for PUS7 under normal growth conditions and puromycin treatment, represented by intensity mapping. Statistical significance was assessed using Student’s two-sample t-test (unpaired, one-sided), with Bonferroni correction applied; not significant (n=133).

**Extended Data Fig. 18.**
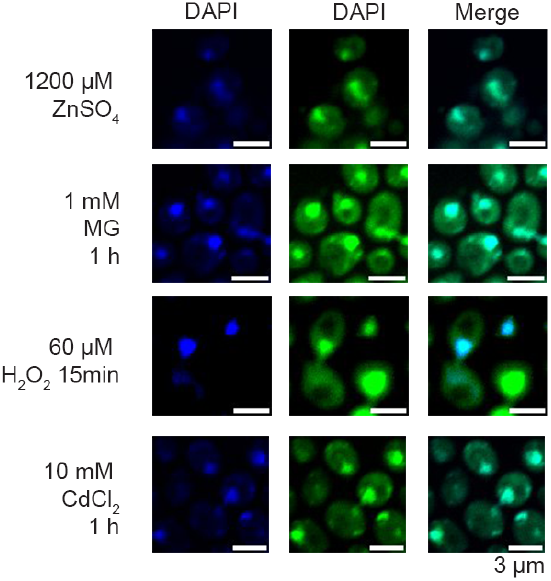
PUS7 relocalization under divalent metal and ROS-related stress conditions. Visualization of PUS7 localization (PUS7–EYFP) under various stress treatments, including ZnSO_4_ (1200 µM, 1 h), MG (1 mM, 1 h), H_2_O_2_ (60 µM, 15 min), and CdCl_2_ (10 mM, 1 h). Scale bar: 3 µm.

**Extended Data Fig. 19.**
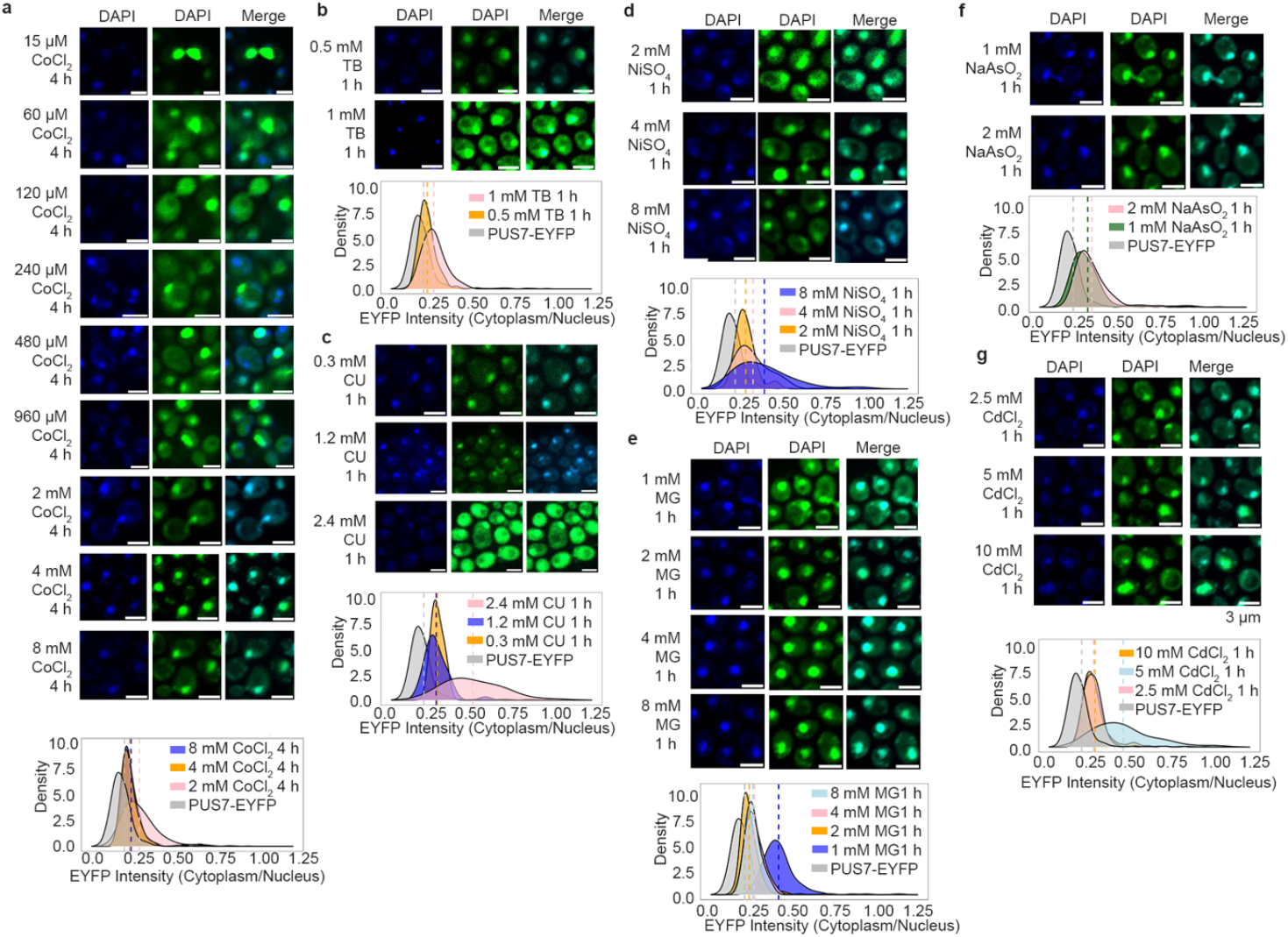
Dose-dependent changes in stress-induced PUS7 relocalization. Visualization and quantification of PUS7 localization (PUS7–EYFP) under varying concentrations of stressors: CoCl_2_ for 4 h (**a**), TB for 1 h (**b**), CU for 1 h (**c**), NiSO_4_ for 1 h (**d**), MG for 1 h (**e**), NaAsO_2_ for 1 h (**f**), and CdCl_2_ for 1 h (**g**). Scale bar: 3 µm.

**Extended Data Fig. 20.**
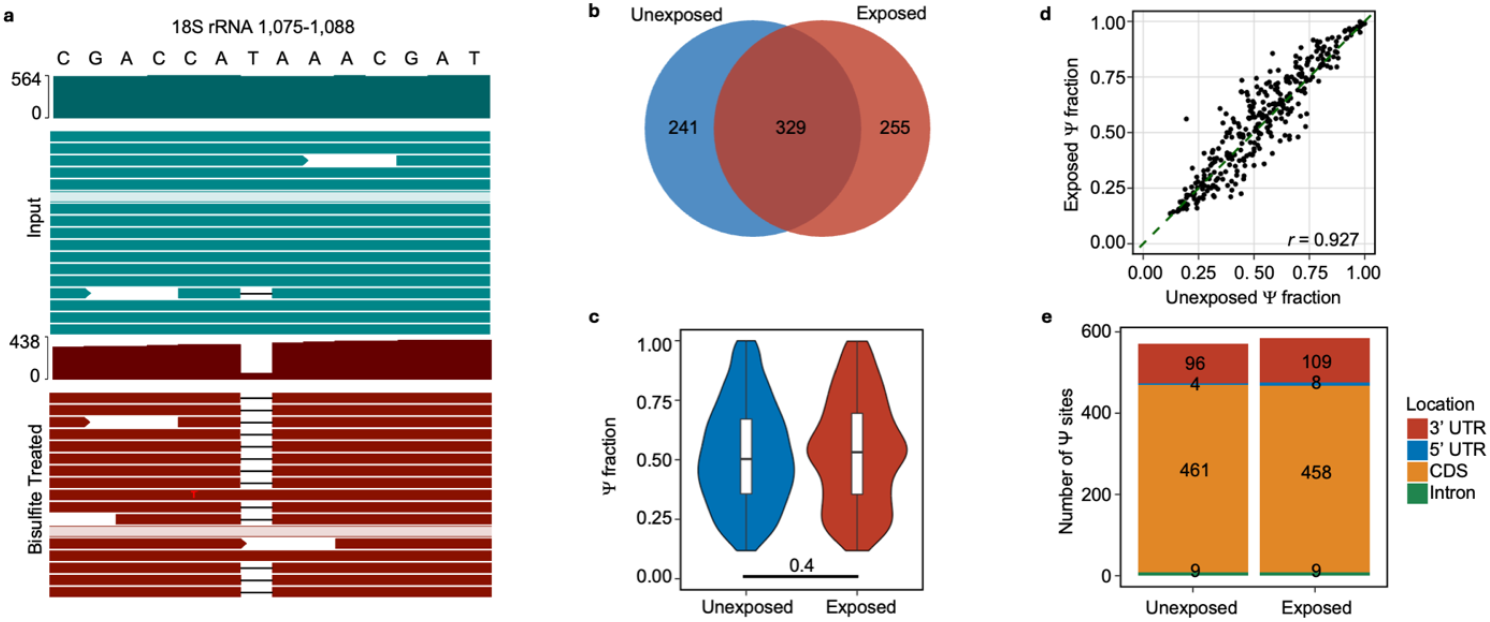
Unique Ψ sites emerge in human lung cells upon exposure to particulate matter, with many sites remaining conserved across exposed and unexposed conditions. **a**, A known Ψ site in rRNA detected in BID-seq. **b**, overlap of Ψ sites between unexposed cells and cells exposed to 125 μg/mL urban PM for 24h. **c**, Violin plots overlaid with box plots showing the distribution of ratios of Ψ incorporation for all detected Ψ sites in both the unexposed and exposed conditions. Welch’s t-test (unpaired, two-tailed). Box center line = median; box limits = 1st (Q1) and 3rd quartiles (Q3). **d**, Comparing the Ψ fraction at each site that is common between the unexposed and exposed conditions shows a high level of correlation (R=0.927) using a Pearson correlation. **e**, Distribution of Ψ sites across transcript regions remains similar, mainly in the coding region (*n=3*).

**Extended Data Fig. 21.**
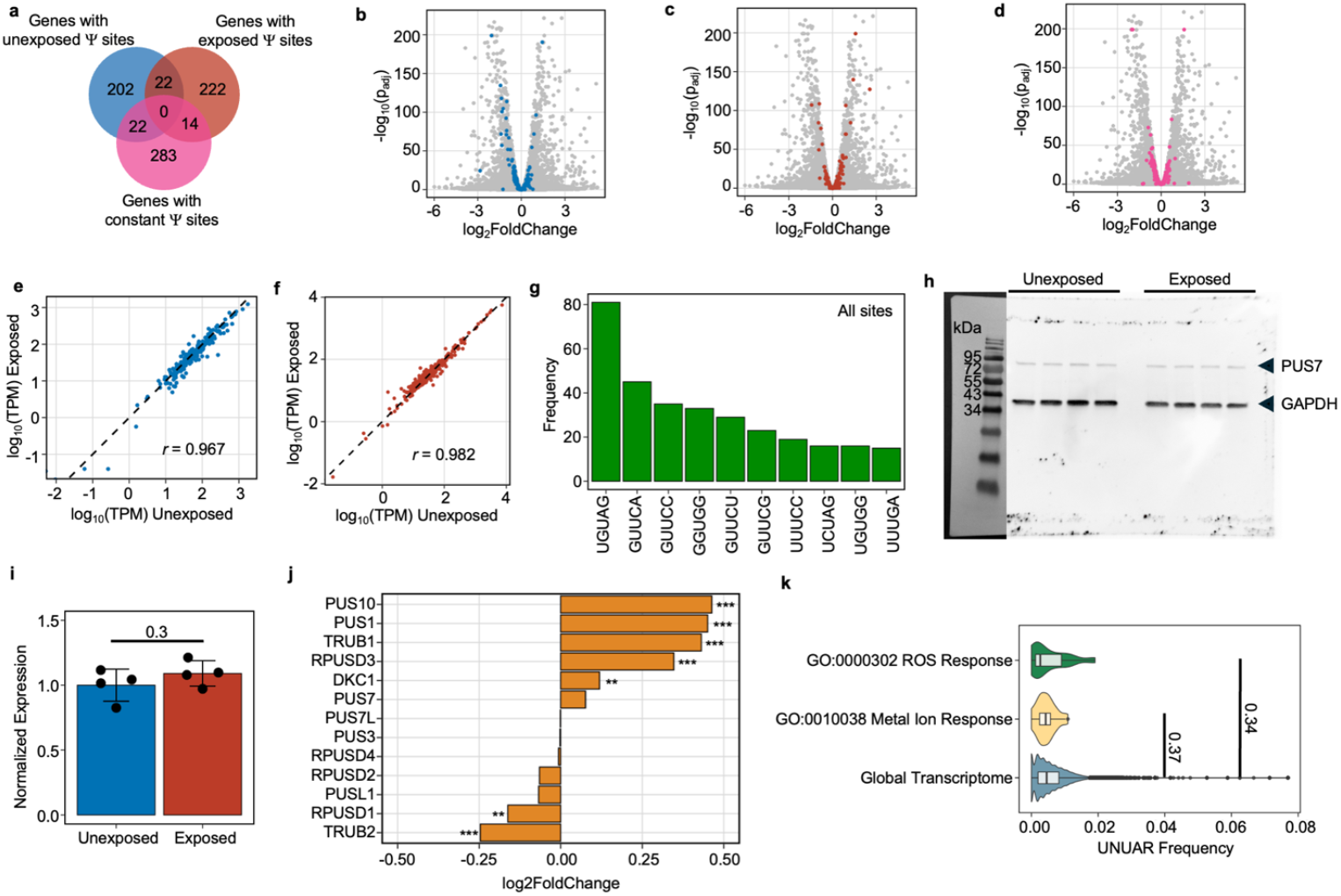
Specific PUS7 motifs get modified more in human lung cells under particulate matter exposure without RNA or protein abundances changing in a specific direction. **a, Ψ** sites are distributed across different transcripts. Some transcripts contain sites unique to either condition, along with sites that remain modified in both conditions. **b-d**, Volcano plots showing differential expression analysis of genes from RNA-seq data (1.6% unique sites in exposure are upregulated, and 0.4% down, |log_2_(FC)|>1). A positive Log_2_FoldChange indicates a gene was more highly expressed in the exposed condition, relative to the unexposed, and vice versa. Colored points represent genes with pseudouridylation sites that were unique to the unexposed (**b**), exposed (**c**) conditions, or common in both (**d**). Padj determined by Wald test with Benjamini Hochberg adjustment (n=3). **e-f**, Correlation plots comparing the relative RNA abundance (log_10_(TPM)) in exposed and unexposed conditions of RNAs that possess a Ψ site unique to the unexposed (**e**) or exposed (**f**) condition. A high degree of correlation was observed for unique unexposed and unique exposed Ψ sites (*r* = 0.967, *r* = 0.982, respectively, Pearson’s correlation coefficient). **g**, Number of detected Ψ sites that fell within the top 10 five-nucleotide motifs for all sites. **h**, Western blot for PUS7 protein levels in unexposed and exposed conditions (125 µg/mL PM, 24 h.). **i**, Densitometry quantification of (**h**). Significance determined using Welch’s t-test (unpaired, two-tailed), mean ± SD is depicted (n=3). **j**, Differential mRNA expression levels of Ψ writer enzymes between unexposed and exposed conditions. Significance determined by Wald test with Benjamini Hochberg adjustment (n = 3, (*** Padj < 1e-07, ** Padj < 0.01). A positive Log_2_FoldChange indicates a gene was more highly expressed in the exposed condition, relative to the unexposed, and vice versa. **k**, Boxplots overlaid onto violin plots depicting enrichment of UNUAR nucleotide motif within ROS response, Metal ion response GO pathways relative to the global transcriptome—significance determined using the Mann-Whitney U test (unpaired, two-tailed).

**Extended Data Fig. 22.**
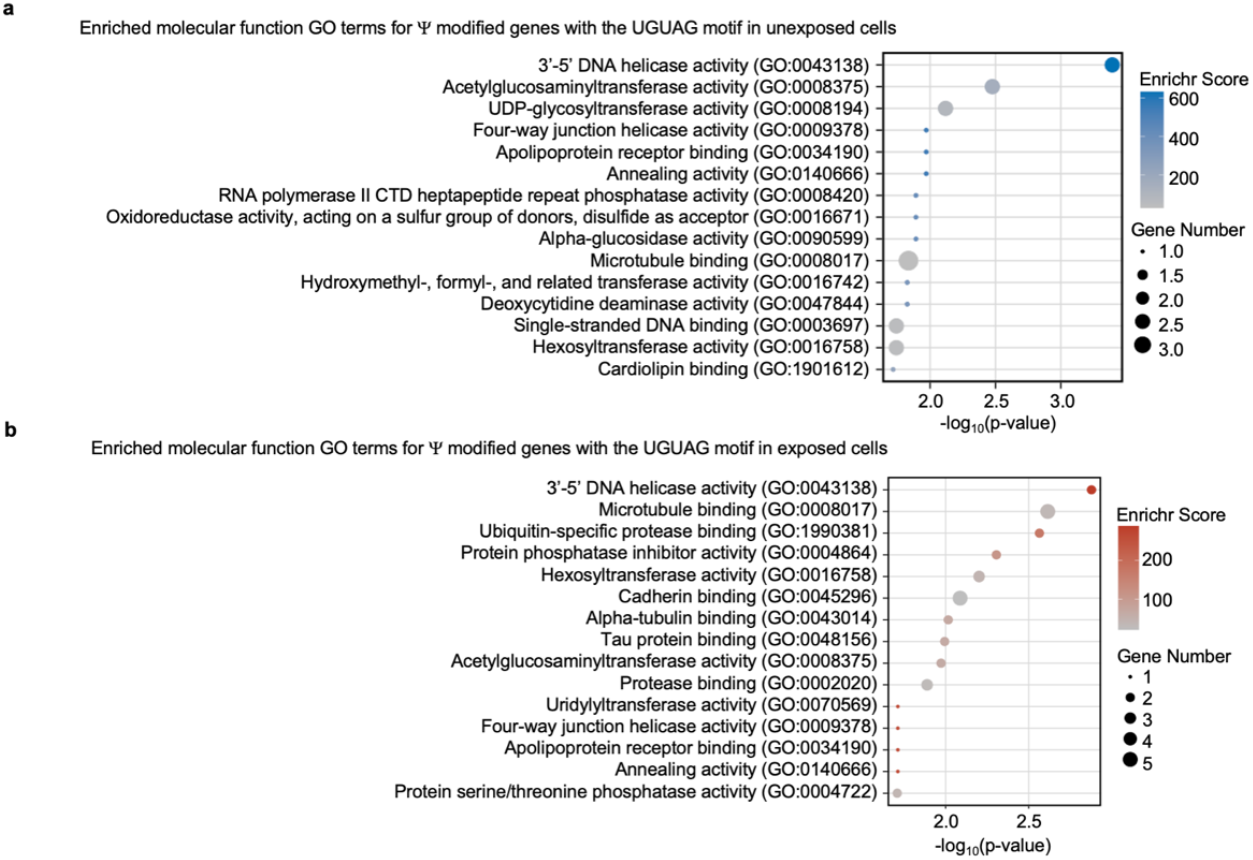
PUS7 differentially modifies mRNA transcripts in human lung cells upon exposure to environmental pollutants. **a-b**, Bubble plots showing the top 15 (P-value ranked) GO molecular function terms enriched in genes containing Ψ modified UGUAG motifs in either the unexposed (**a**) or exposed (**b**) condition.

**Extended Data Fig. 23.**
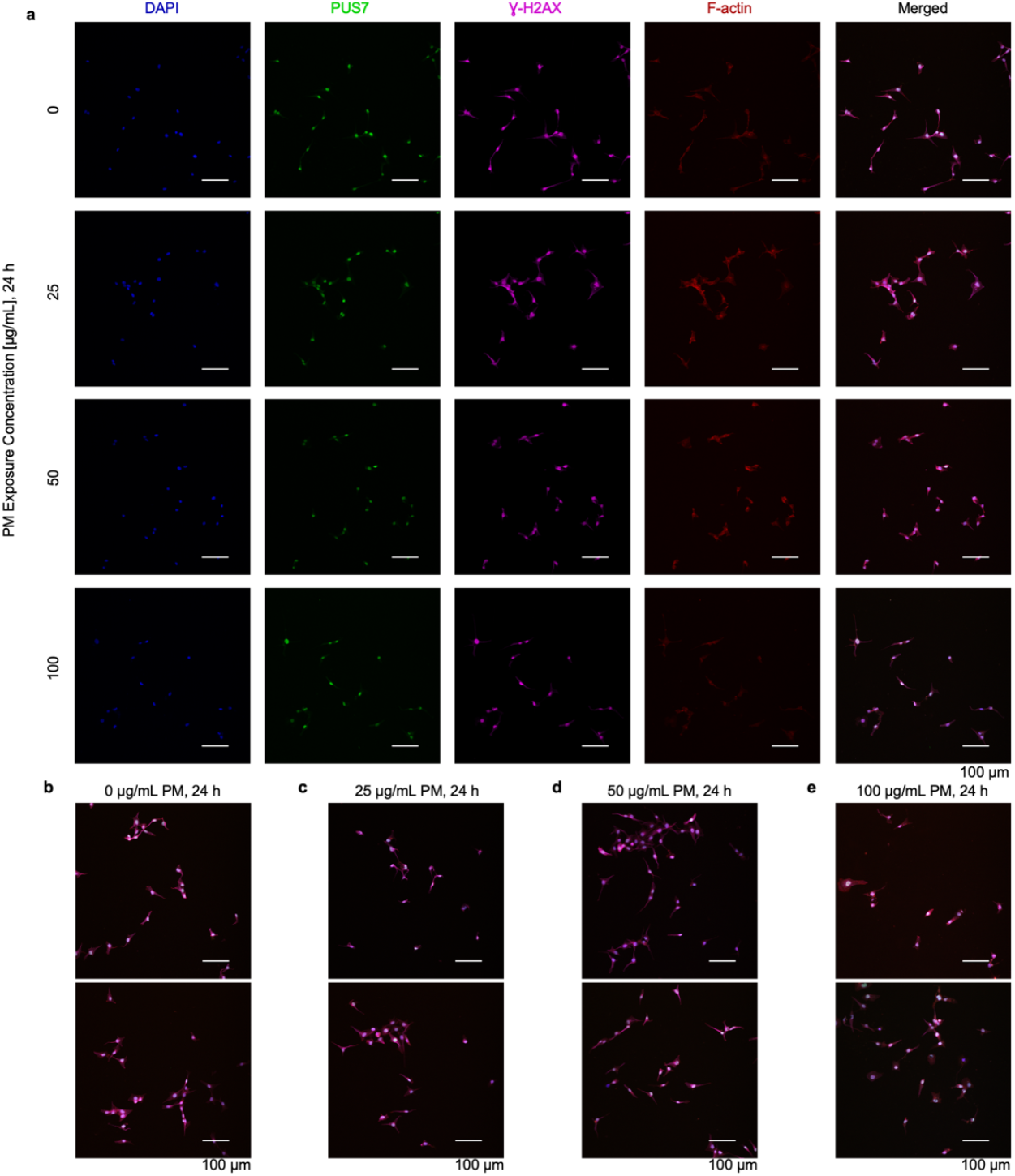
Representative immunofluorescence images of BEAS-2B cells under increasing PM exposure conditions or media-only control. Nuclei (blue), PUS7 (green), Ɣ-H2AX (magenta), F-actin (red). Scale bar: 100 µm. **a**, fluorescence channel separated images. **b-d**, additional merged images of BEAS-2B cells under media only control (**b**), 25 µg/mL PM exposure (**c**), 50 µg/mL PM exposure, and (**d**) 100 µg/mL PM exposure.

**Extended Data Fig. 24.**
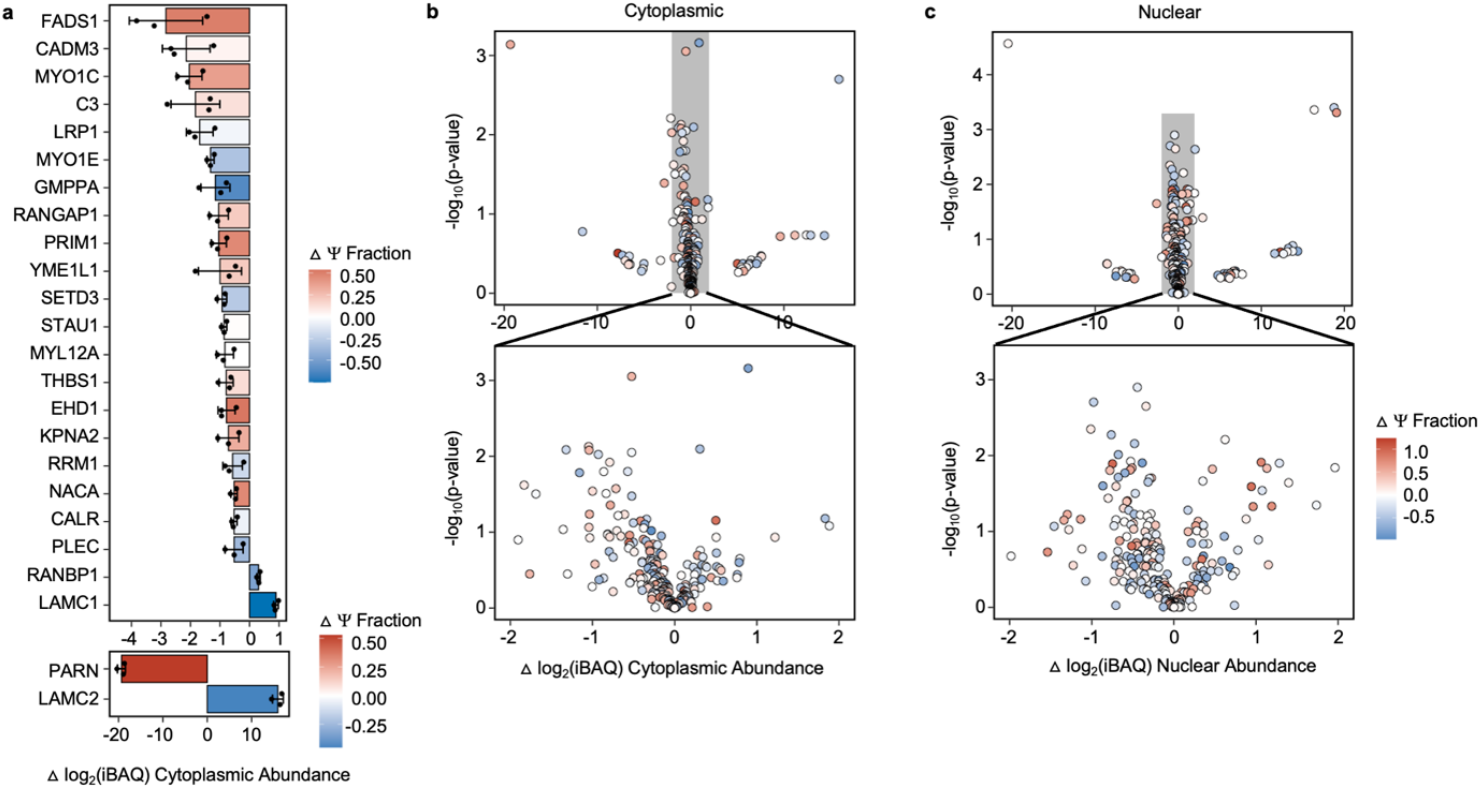
Proteins with significantly altered abundancies following PM exposure. **a**, Bar plots displaying the change in iBAQ score between the exposed and unexposed conditions for proteins in the cytoplasmic fraction as measured by mass spectrometry. A positive value indicates an increase in protein in the exposed condition relative to the unexposed condition, while a negative value indicates the opposite. Bars are colored to indicate the change in Y incorporation between the unexposed and exposed conditions, where a positive value indicates an increase in ψ incorporation in the exposed condition and vice versa (p < 0.05, Welch’s t-test (unpaired, two-tailed)). Mean ± SD is depicted (n=3). **b-c**, All proteins detected through mass spectrometry in the cytoplasmic (**b**) and nuclear (**c**) fractions whose cognate mRNA transcripts contained at least one Y modification in either the exposed or unexposed condition. Points are colored to indicate the change in ψ incorporation between the unexposed and exposed conditions, where a positive value indicates an increase in ψ incorporation in the exposed condition and vice versa. (n = 3). Bottom panels are zoomed-in versions of the top panels.

