## Supplementary Figures for "Cytoplasmic localization of PUS7 facilitates a pseudouridine-dependent enhancement of cellular stress tolerance"

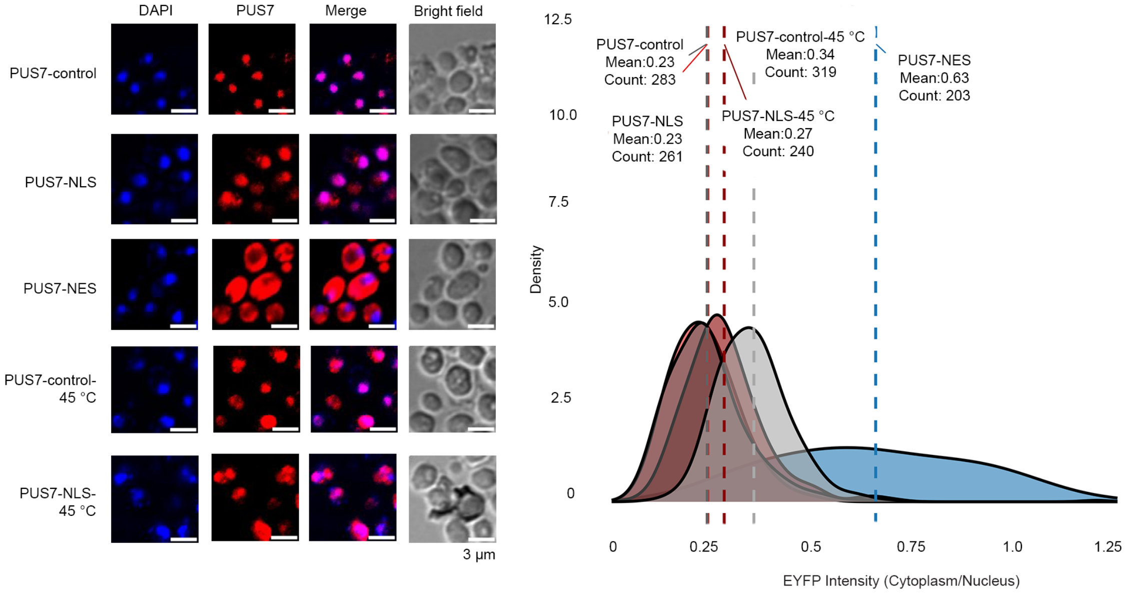


**Supplementary Fig. 1 | Immunofluorescence of anti-FLAG showing localization of PUS7-control, -NLS, and -NES fused PUS7 as well as PUS7-control and -NLS under heat shock (heat shock (45 °C for 15 min)), together with quantification of the Fluorescence Intensity (FI) ratio in the cytoplasm *vs* in the nucleus (n>100 cells).** Scale bar: 3 μm.

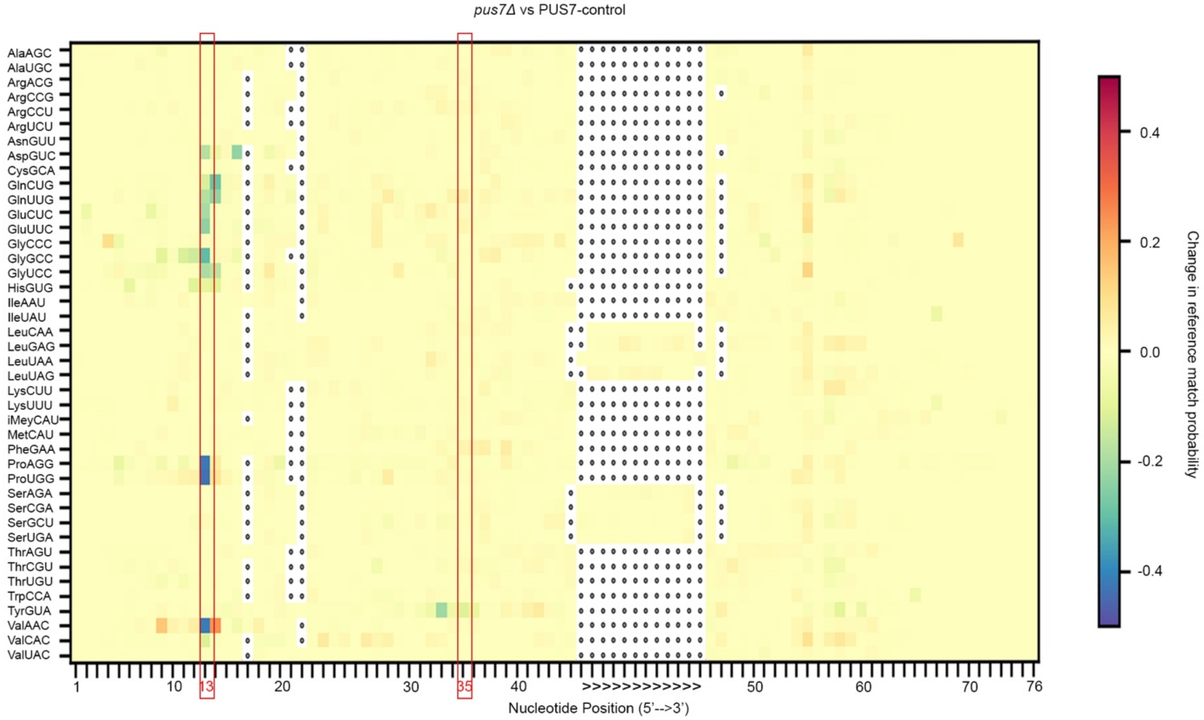


**Supplementary Fig. 2 | Ψ13 and Ψ35 modifications appear upregulated in PUS7-control compared to *pus7*Δ cells, highlighted in red boxes.**

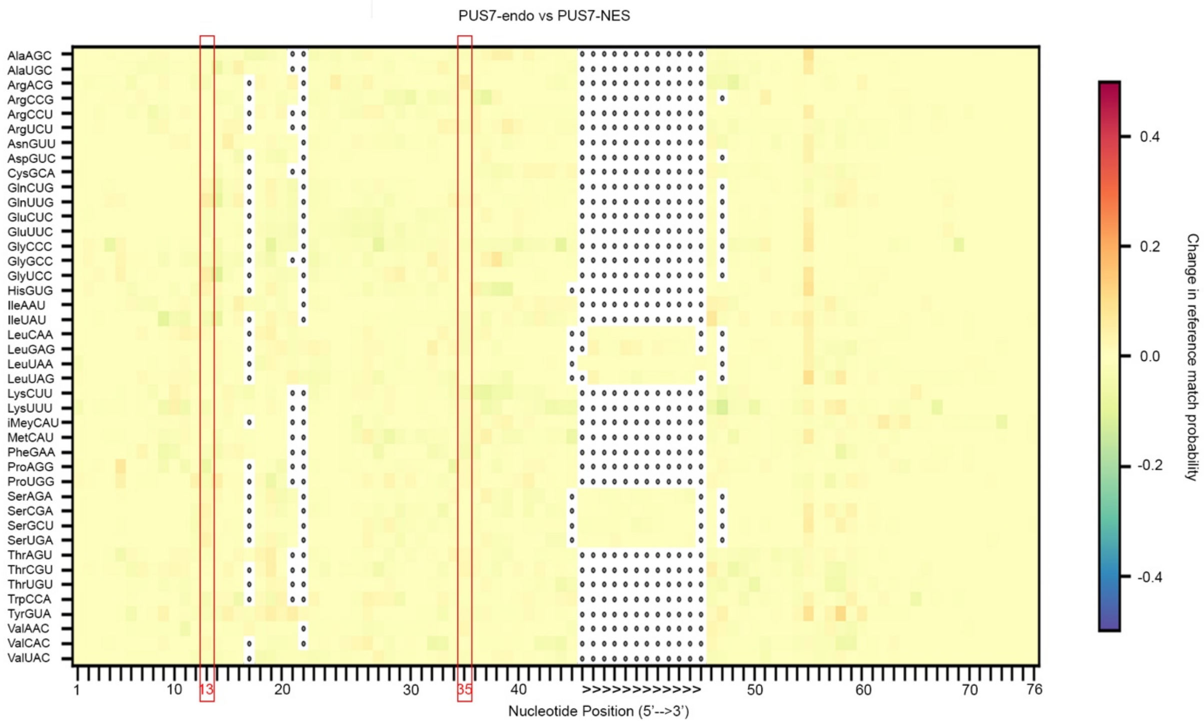


**Supplementary Fig. 3 | Ψ incorporation on tRNA positions targeted by PUS7 in eukaryotes (Ψ13 and Ψ35) is unaltered between cells expressing only endogenous PUS7 and PUS7-NES, highlighted in red boxes.**

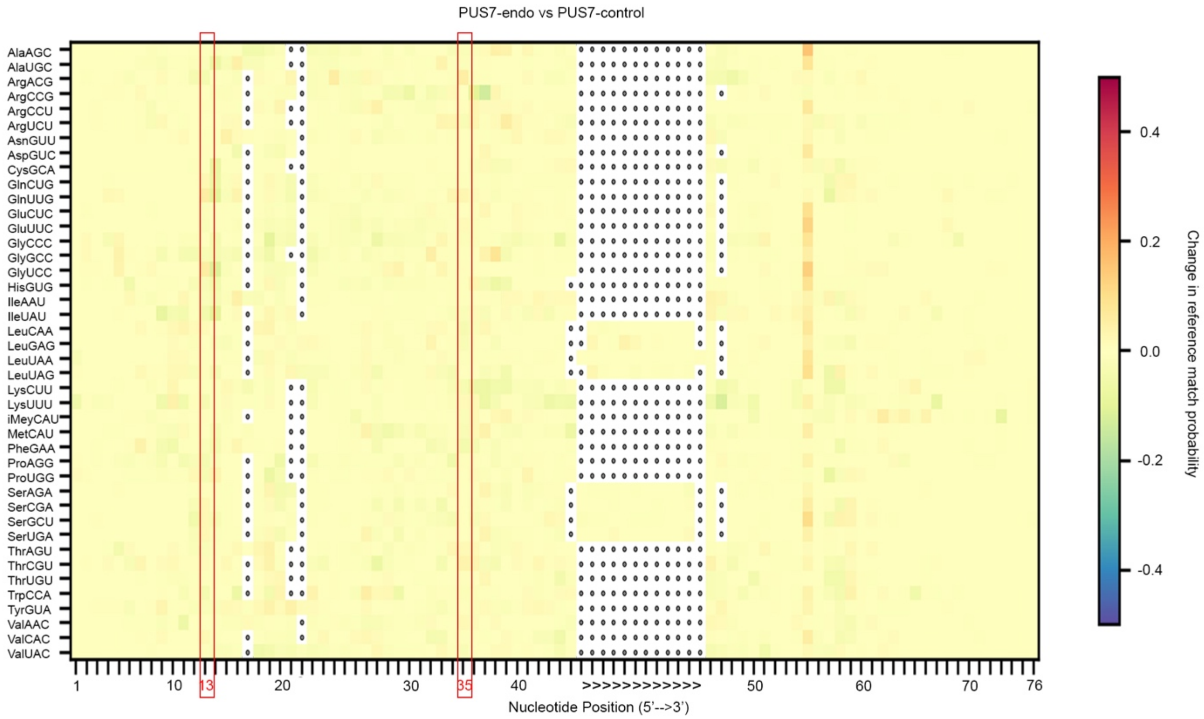


**Supplementary Fig. 4 | Ψ incorporation on tRNA positions targeted by PUS7 in eukaryotes (Ψ13 and Ψ35) is unaltered between cells expressing only endogenous PUS7 and PUS7-control, highlighted in red.**

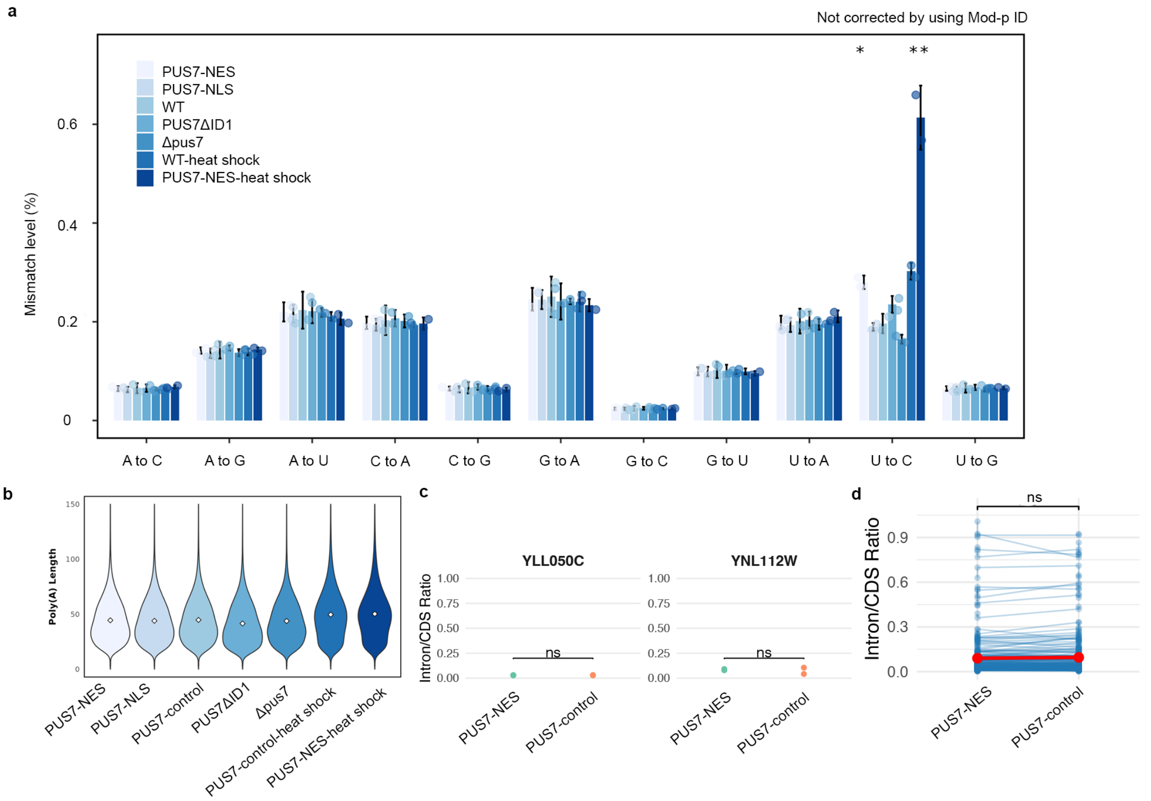


**Supplementary Fig. 5 | U-to-C mismatch level changes observed in different cell groups without affecting poly(A) length or pre-mRNA splicing. a**, Nucleotide mismatch levels for all four nucleotides across different cell groups from RNA002. Student's two-sample t-test (unpaired, two-tailed) with Bonferroni correction was used to calculate adjusted p-values. Significant U-to-C mismatch increases were observed in PUS7-NES ($p = 0.038$), PUS7-FLAG-heat shock ($p = 0.029$), and PUS7-NES-heat shock ($p = 0.013$). Data are shown as mean ± SD. (*$P < 0.05$; $n = 2$). **b**, polyA length is not getting affected among different groups. **c**, PUS7 relocalization does not broadly affect RNA splicing. Intron/CDS read ratios are shown for COF1/YLL050C (Ψ site detected in PUS7-control) and DBP2/YNL112W (Ψ site detected in PUS7-NES); paired, two-tailed t-test, $n = 2$. **d**, Overall splicing levels of 172 genes containing introns are similar between PUS7-NES and PUS7-control ($n = 172$).

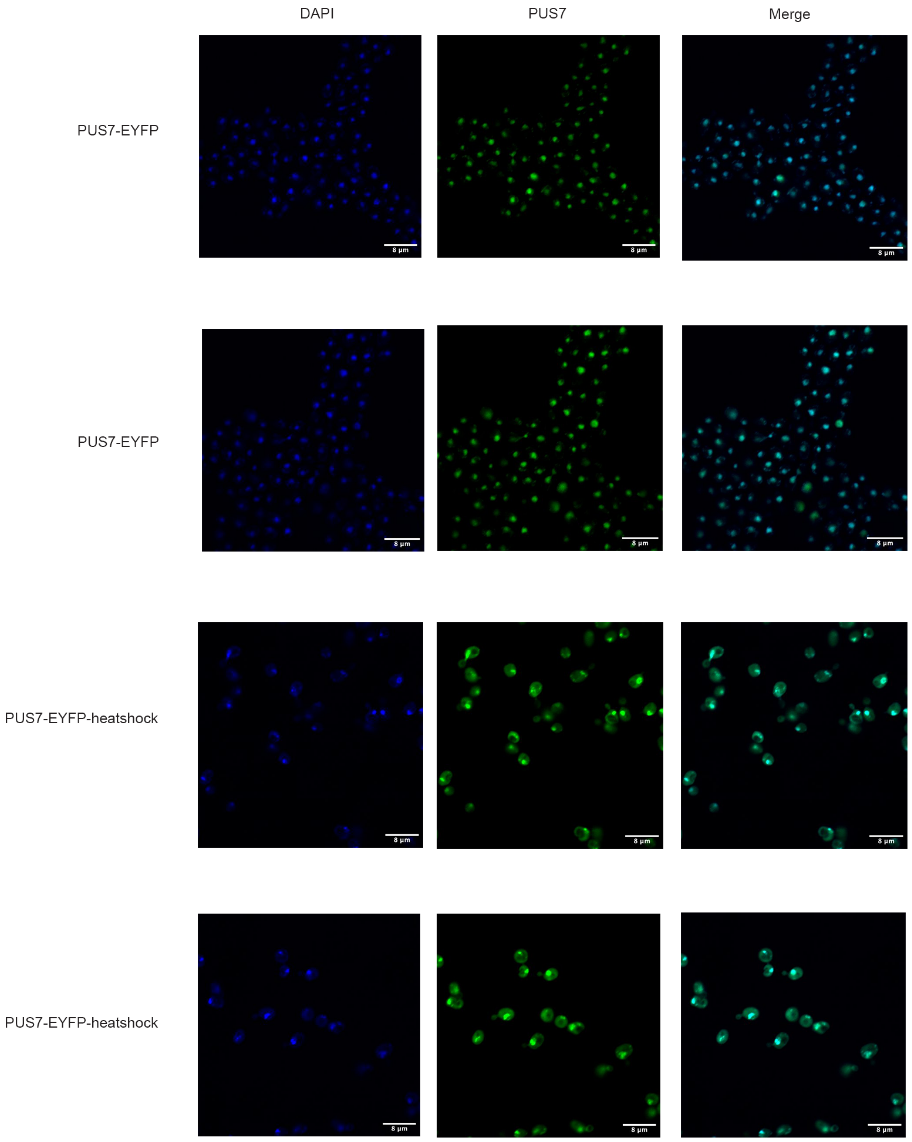


**Supplementary Fig. 6 | More images showing PUS7-EYFP relocalization under no stress conditions and heat shock (45 °C, 1h).** Scale bar: 8 μm.

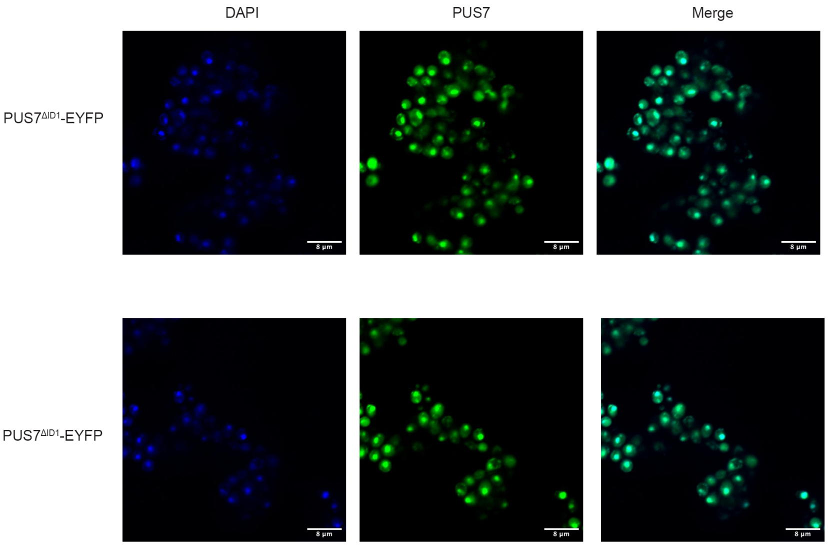


**Supplementary Fig. 7 | More images showing PUS7 relocalization for cell expression PUS7$^{\Delta ID1}$-EYFP.** Scale bar: 8 µm.

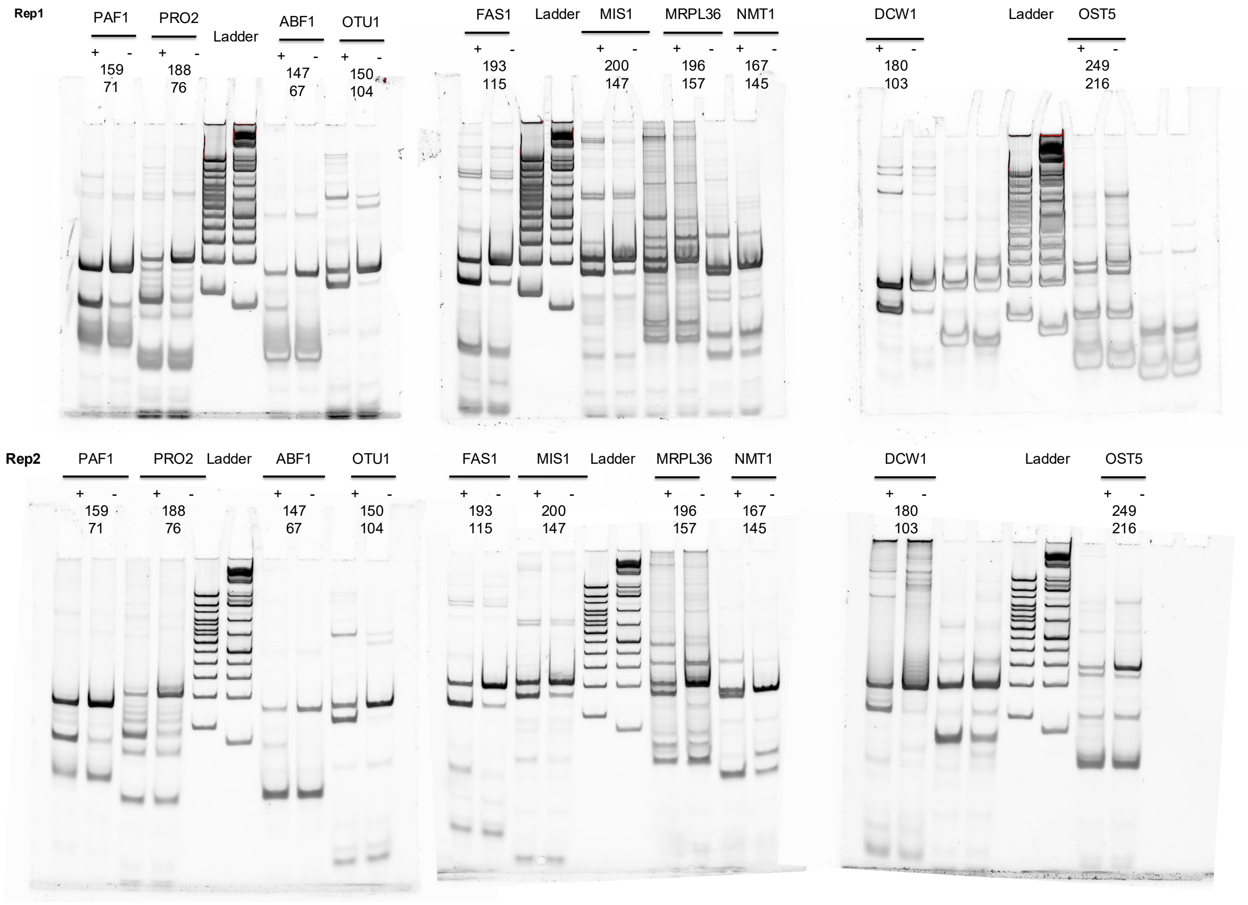


**Supplementary Fig.8 | CLAP gels.**

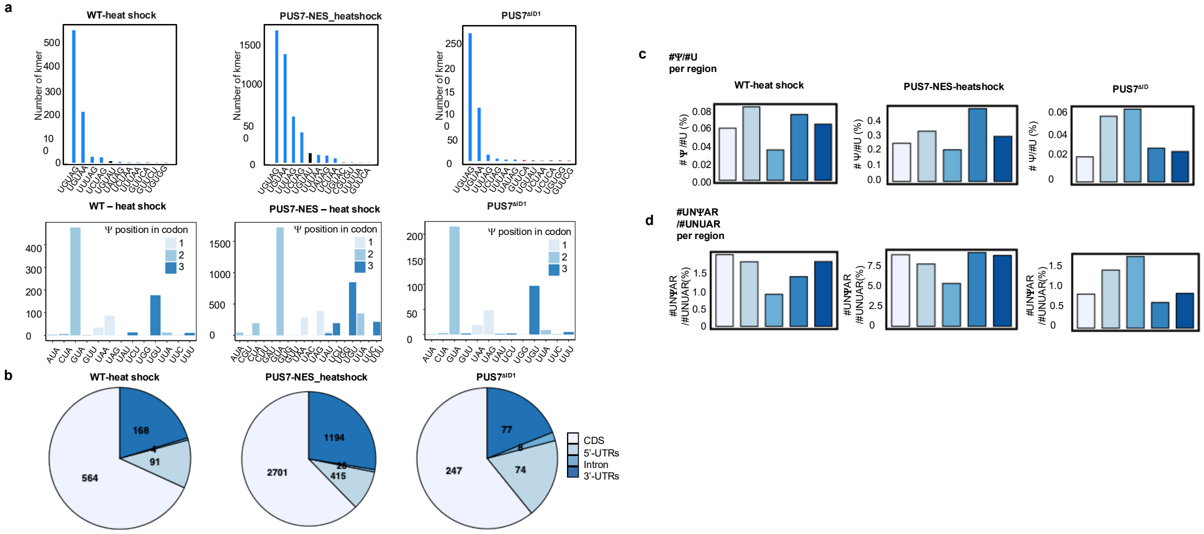


**Supplementary Fig.9 | High-confidence Ψ sites for different cells under various stress conditions. a,** 5-mer accumulation and codon enrichment for mRNAs isolated from different localized PUS7 cells with U to C mismatch over 40%. **b,** The number of Ψ sites distribution in coding-region, 5'-UTRs, intron, and 3'-UTR for PUS7-control-heat shock, PUS7-NES-heat shock, PUS7$^{\Delta ID}$, respectively. **c,d,** ratio of #Ψ/#U and #UNΨAR/#UNUAR in different regions for PUS7-control-heat shock, PUS7-NES-heat shock, PUS7$^{\Delta ID}$, respectively. ($n = 2$).

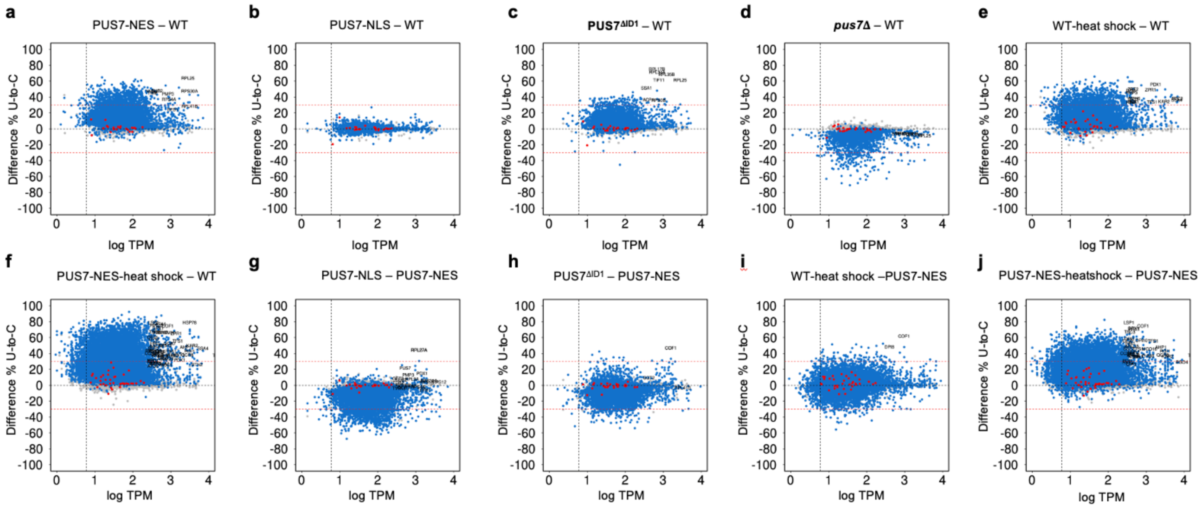


**Supplementary Fig. 10 | Potential Ψ-positions determined are plotted according to the difference of U-to-C base-calling error (%) between different PUS7 localized samples and PUS7-control against the number of total direct reads (log$_{10}$TPM).** (*n=2*). Sites are compared when over 10 reads coverage is achieved on both groups, and a p-value <0.001 is achieved for at least one group (IVT reads>7 and mm_IVT% < 10%).

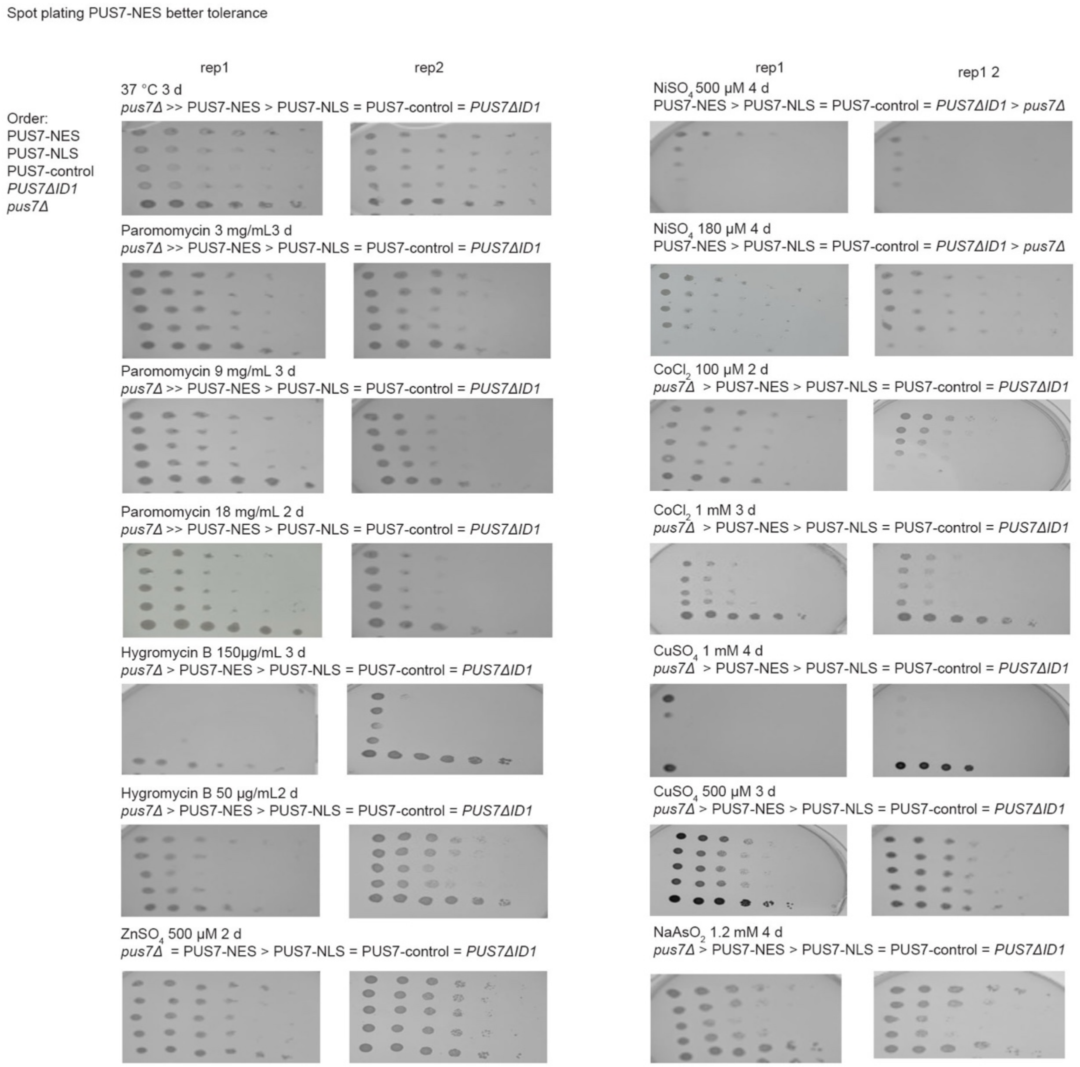


**Supplementary Fig.11 | Spot plating assay for PUS7-NES, PUS7-NLS, PUS7-control, PUS7$^{\Delta ID1}$, and *Pus7*Δ yeast with PUS7-NES showing better tolerance**. The cells are placed in the plate in the order of PUS7-NES, PUS7-NLS, PUS7-control, PUS7$^{\Delta ID1}$, *Pus7*Δ cells (*n=2*).

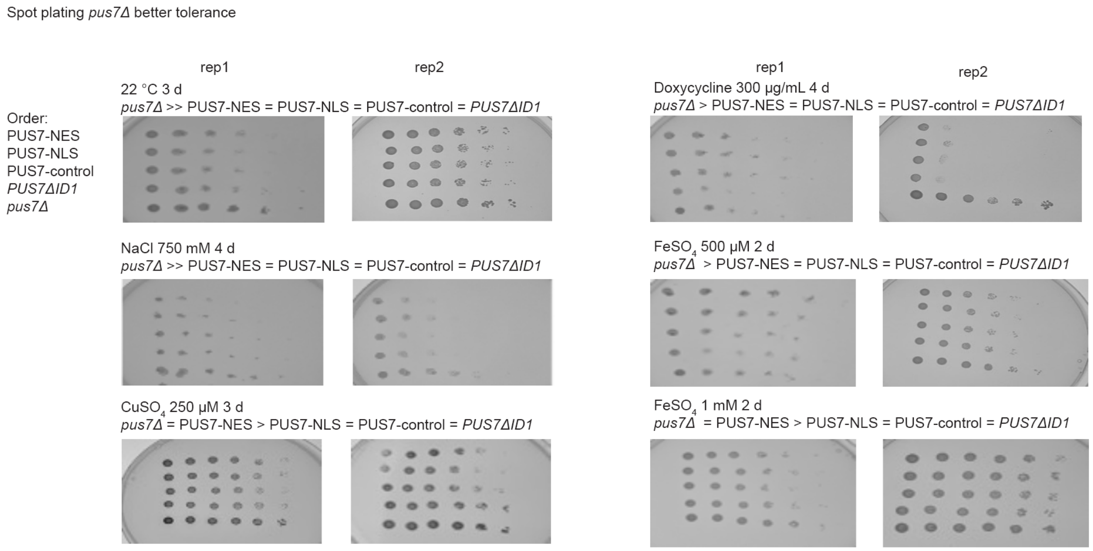


**Supplementary Fig.12 | Spot plating assay for PUS7-NES, PUS7-NLS, PUS7-control, PUS7$^{\Delta ID1,}$ and *Pus7*Δyeast with *Pus7*Δand PUS7-NES or only *Pus7*Δ showing better tolerance.** The cells are placed in the plate in the order of PUS7-NES, PUS7-NLS, PUS7-control, PUS7$^{\Delta ID1}$*, Pus7*Δ cells (*n=2*).

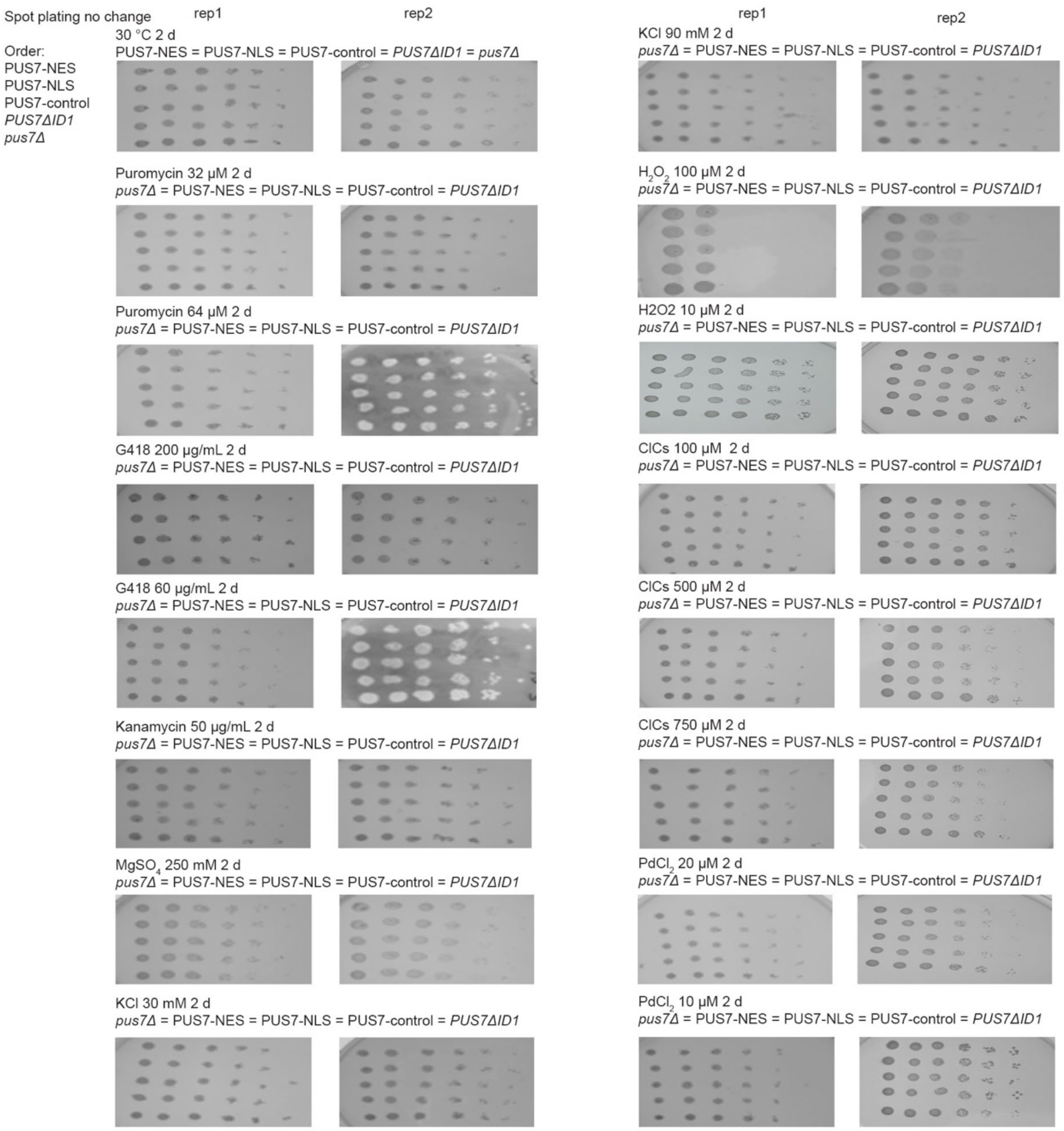


**Supplementary Fig.13 | Spot plating assay for PUS7-NES, PUS7-NLS, PUS7-control, PUS7$^{\Delta ID1}$, and *Pus7*Δ yeast with no difference.** The cells are placed in the plate in the order of PUS7-NES, PUS7-NLS, PUS7-control, PUS7$^{\Delta ID1}$*, Pus7*Δ cells (*n=2*).

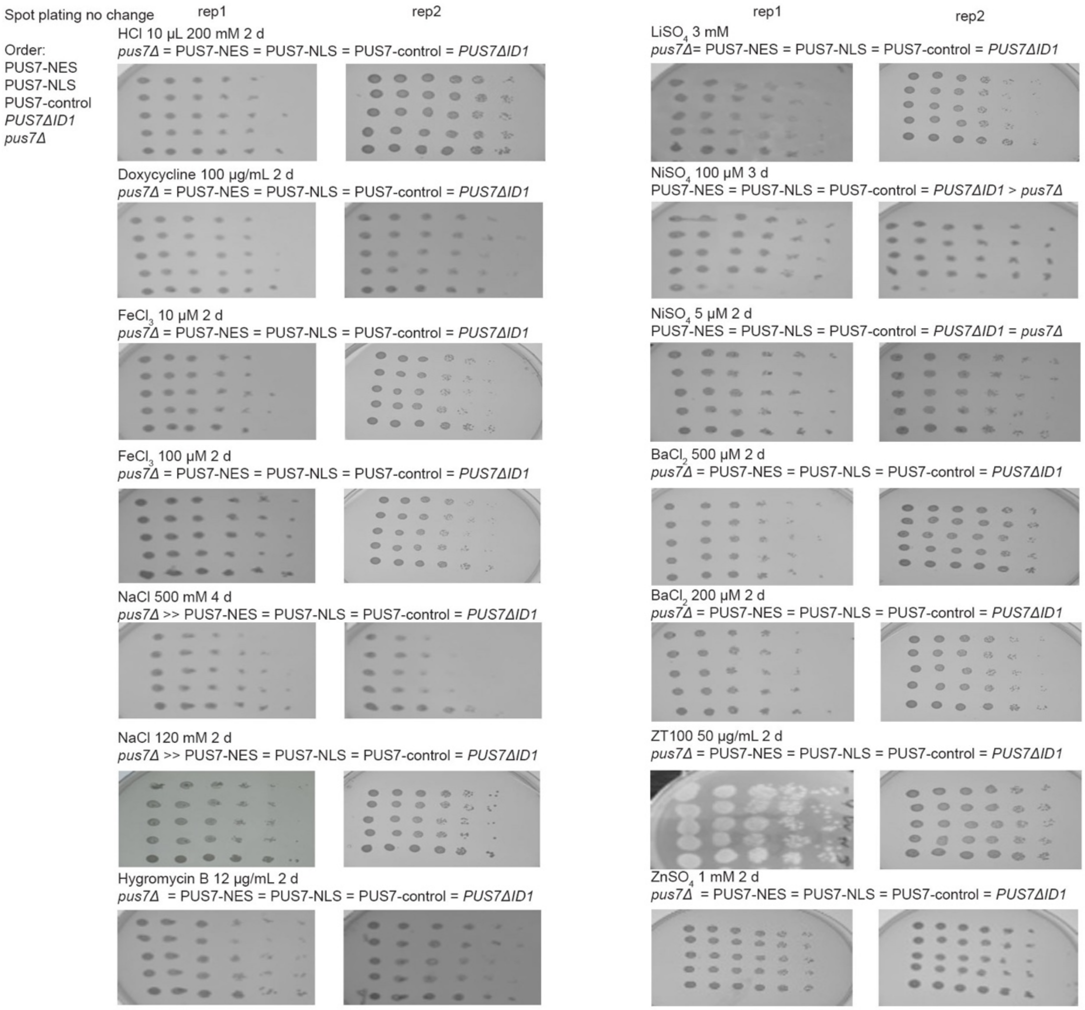


**Supplementary Fig.14 | Spot plating assay for PUS7-NES, PUS7-NLS, PUS7-control, PUS7$^{\Delta ID1}$ and *Pus7*Δ yeast with no difference (part2).** The cells are placed in the plate in the order of PUS7-NES, PUS7-NLS, PUS7-control, PUS7$^{\Delta ID1}$, *Pus7*Δ cells (*n=2*).

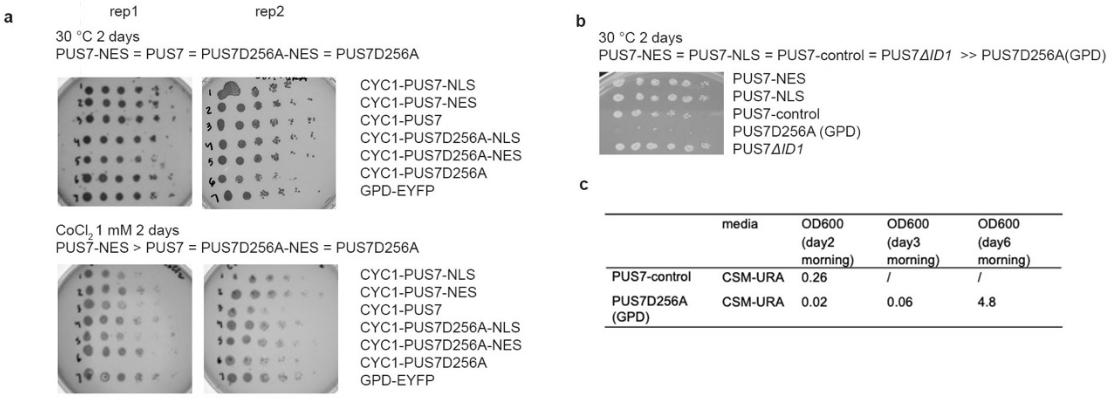


| | media | OD600 (day2 morning) | OD600 (day3 morning) | OD600 (day6 morning) |
|---|---|---|---|---|
| PUS7-control | CSM-URA | 0.26 | / | / |
| PUS7D256A (GPD) | CSM-URA | 0.02 | 0.06 | 4.8 |

**Supplementary Data Fig. 15 | Expression of catalytically inactive PUS7 (D256A) under a low-expression promoter CYC1 in *pus7*Δ BY4741 cells demonstrates that the stress tolerance conferred by PUS7-NES is not due to a scaffold effect. a,** Strains expressing PUS7 or catalytically inactive PUS7 under the lower-expression CYC1 promoter show that the stress tolerance conferred by PUS7-NES is not due to its cytoplasmic localization, as PUS7^D256A-NES does not provide a growth advantage. **b-c,** *pus7*Δ BY4741 cells expressing a catalytically inactive version of PUS7 (D256A) under the GPD promoter display general toxicity, as shown by spot plating (**b**) and liquid media growth (**c**), which leads us to switch to the CYC1 promoter to exclude the scaffold effect.

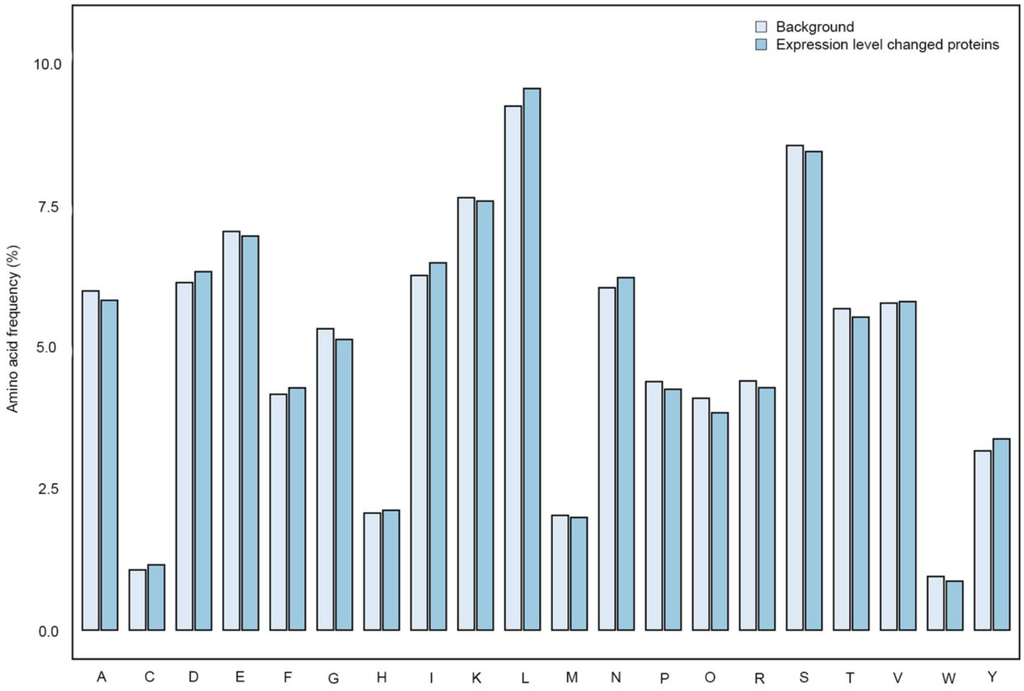


**Supplementary Fig. 16 | Distribution of codons within proteins exhibiting altered expression levels between PUS7-NES-Co(II) *vs* PUS7-control-Co(II) *vs* all detected background proteins.**

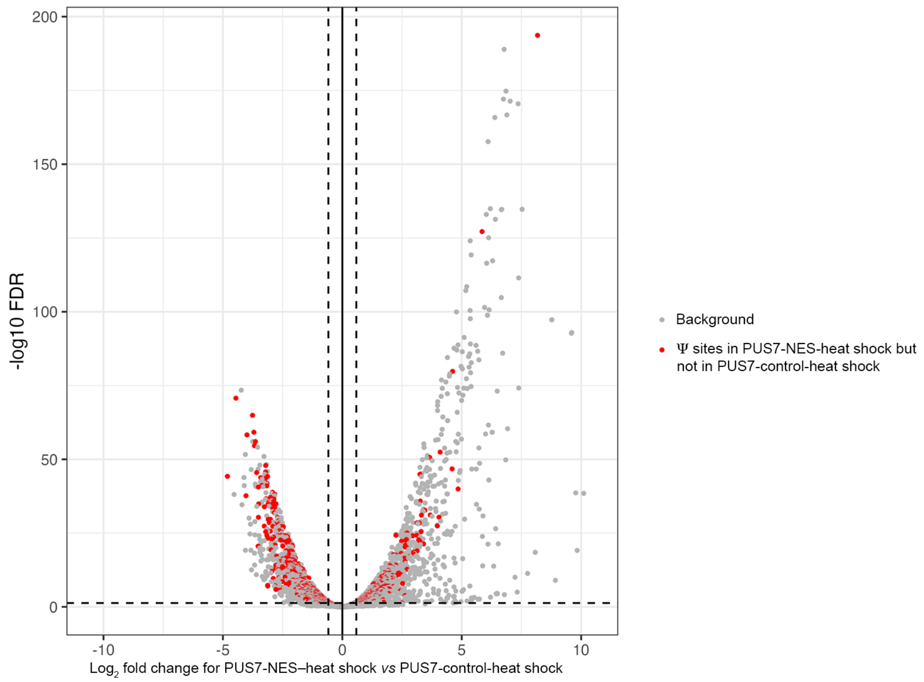


**Supplementary Fig. 17 | Volcano plot for PUS7-NES under heat shock *vs* PUS7-NES under normal conditions, with transcripts containing uniquely incorporated Ψ sites in PUS7-NES-heat shock highlighted in red.**

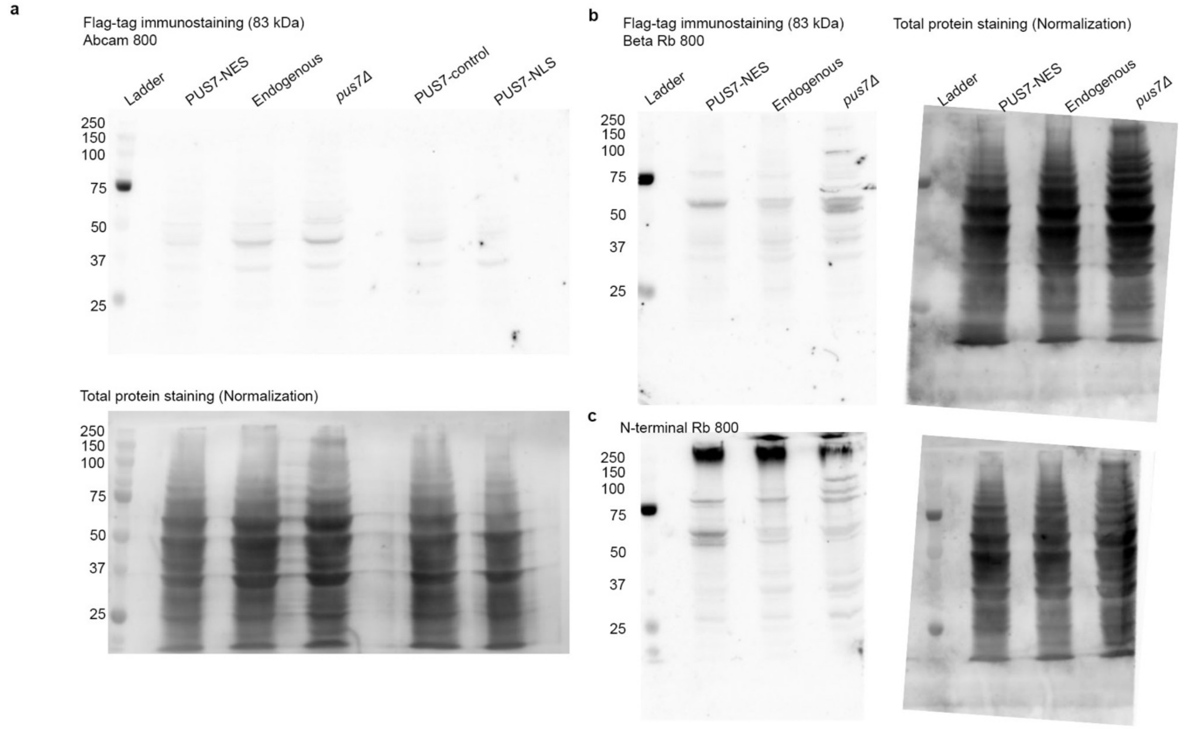


**Supplementary Fig. 18 | Negative result of western blot of PUS7 antibodies in yeast. a,**

ab224119, Abcam rabbit; **b**, #A305-147A, Bethyl Laboratories; **c**, ABIN310801, antibodies online.

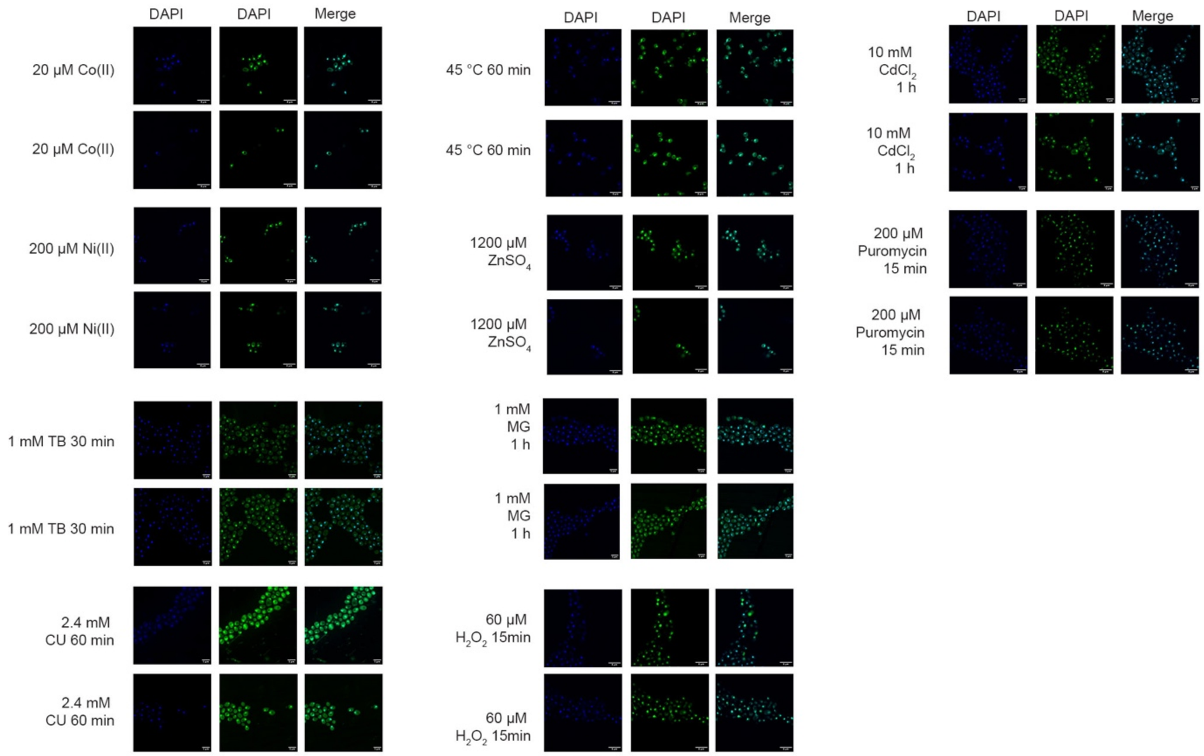


**Supplementary Fig. 19 | Additional images of PUS7 localization (PUS7–EYFP) under various stress conditions.** Stress conditions include: cobalt (30 µM, 4 h), nickel (200 µM, 4 h), TB (1 mM, 30 min), CU (2.4 mM, 1 h), heat shock (1 h), $ZnSO_4$ (1200 µM, 1 h), MG (1 mM, 1 h), $H_2O_2$ (60 µM, 15 min), $CdCl_2$ (10 mM, 1 h) and puromycin (200 µM, 15 min). Scale bar: 3 µm.

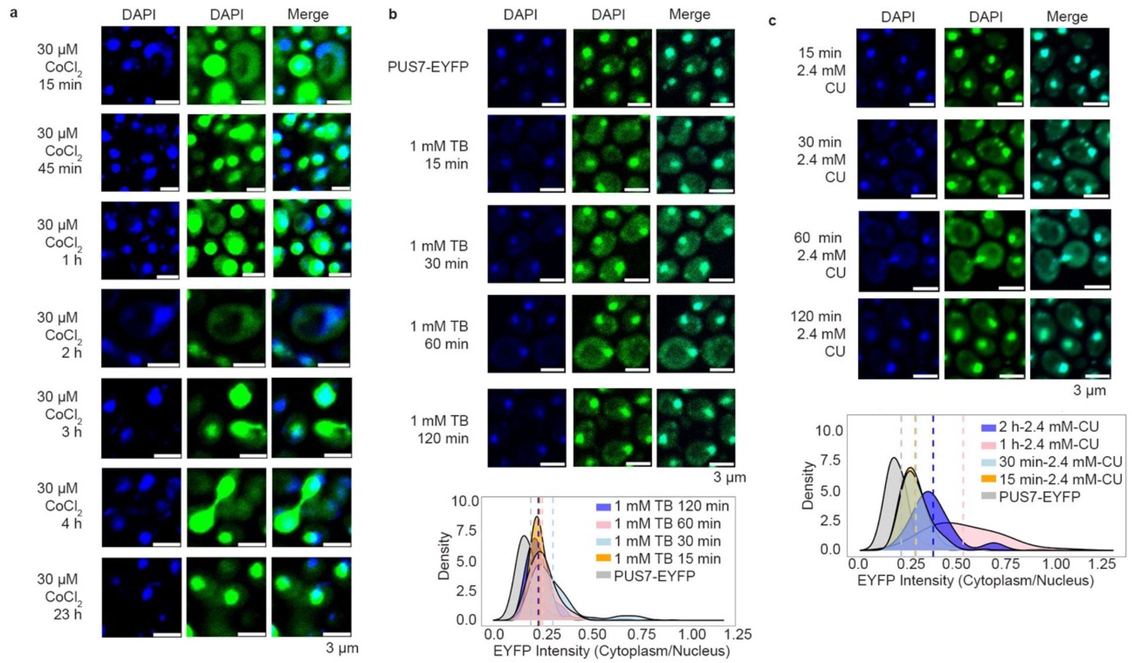


**Supplementary Fig. 20 | PUS7 localization changes under stress are time dependent.** Visualization and quantification of PUS7 localization (PUS7–EYFP) under 30 μM $CoCl_2$ (**a**), 1 mM TB (**b**), and 2.4 mM CU (**c**) across different time points. Scale bar: 3 μm.

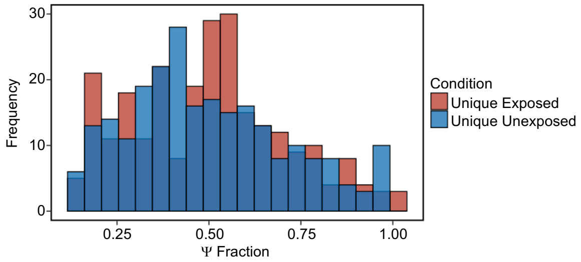


**Supplementary Fig. 21 | Distributions of Ψ site fractions that are unique to PM-exposed and unexposed conditions. Histogram showing the frequency of unique Ψ sites for each condition across increasing modification fraction level.**

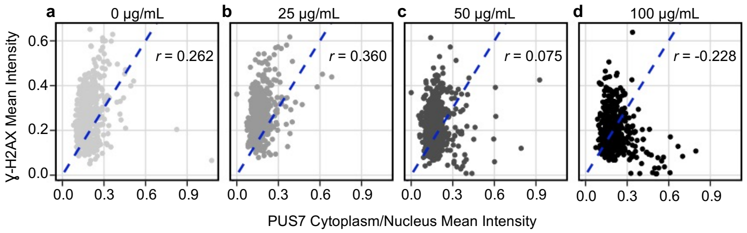


**Supplementary Fig. 22 | PUS7 relocalization in BEAS-2B cells exposed to particulate matter is not correlated with DNA damage.** Quantification of ɣ-H2AX mean intensity for individual cells exposed to increasing levels of particulate matter for 24 h, measured through immunofluorescence staining. (*n*=400+ *cells per condition*). Correlation plots showing the relationship between ɣ-H2AX mean intensity and the cytoplasmic-to-nuclear ratio of PUS7 mean intensity for these individual cells. Dashed blue lines represent y = x. r = Pearson's correlation coefficient.

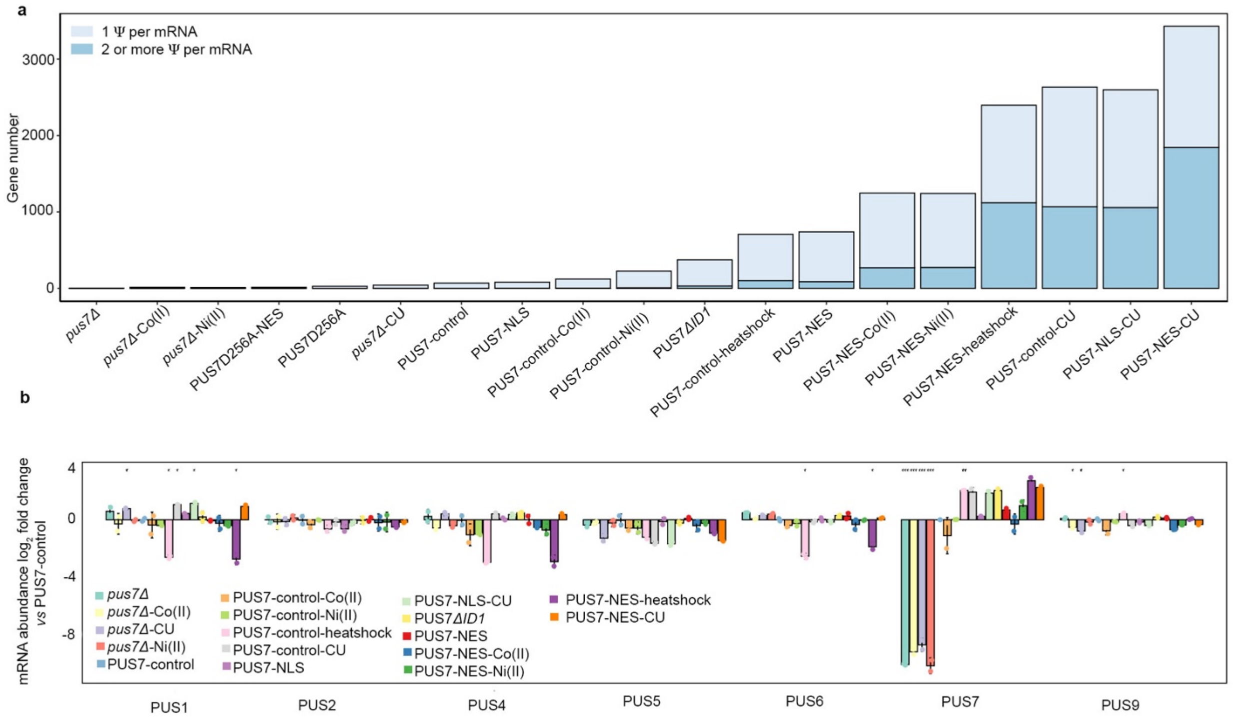


**Supplementary Fig. 23 | Ψ distribution and PUS enzyme abundance (TPM) changes under different PUS7 localization and stress conditions. a**, Number of genes with one Ψ site versus multiple Ψ sites in different cell groups. **b**, Changes in mRNA abundance for various PUS enzymes when PUS7 localization differs or under heat shock. Data are shown as mean ± SD. Welch's two-sample t-test (unpaired, two-sided) was used. *P < 0.05, **P < 0.01, ***P < 0.001 (n = 2).

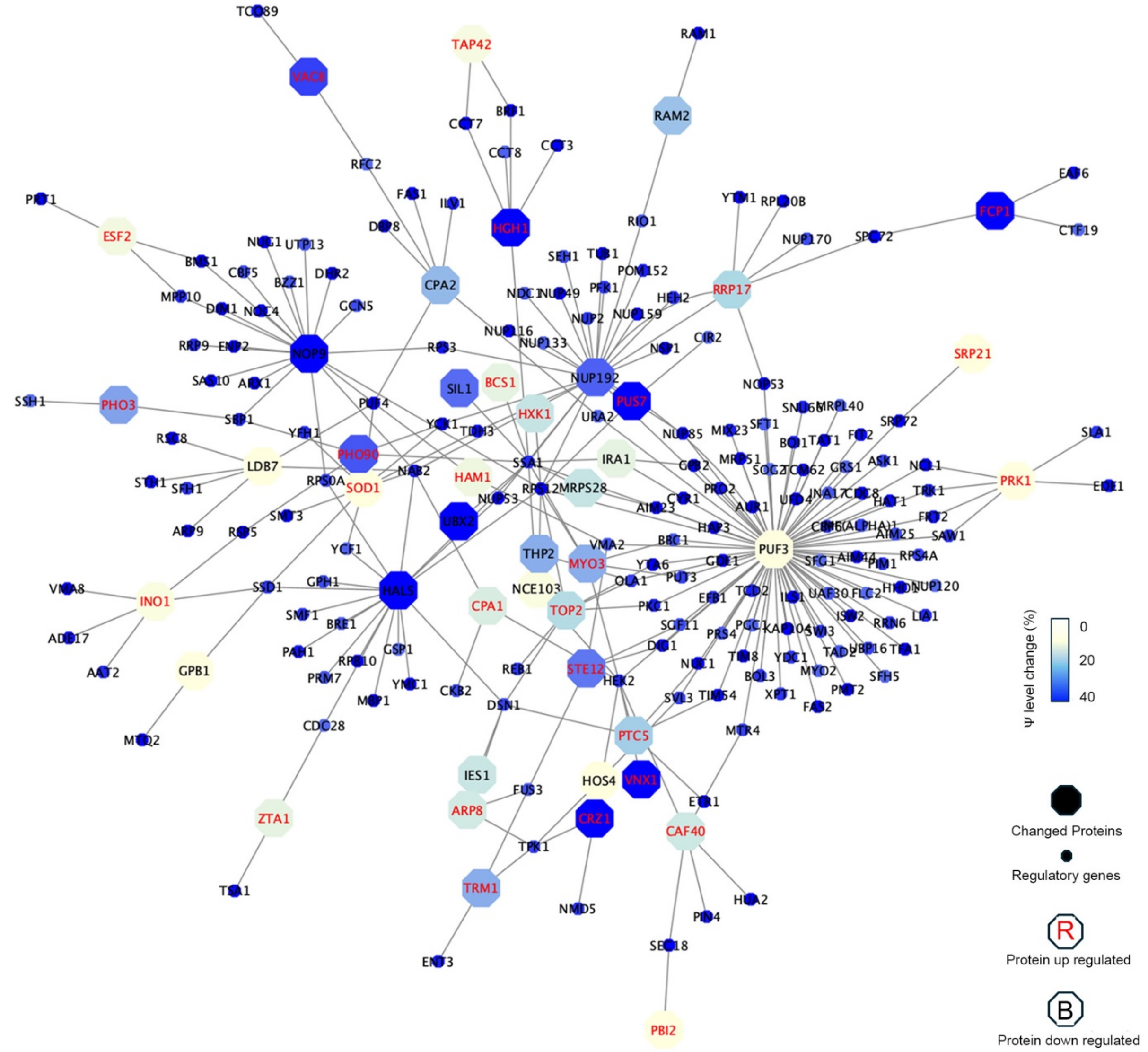


**Supplementary Fig. 24 | Physical interactions between expression level changed proteins and proteins whose encoding RNAs are robustly being incorporated by Ψ in PUS7-NES-Co(II) but not PUS7-control-Co(II) (30 µM cobalt stress for 4 h).** 48 out of 68 proteins with expression level changes between PUS7-NES-Co(II) and PUS7-control-Co(II) have physical interaction with genes whose encoding RNAs are robustly incorporated by Ψ in PUS7-NES-Co(II) but not PUS7-control-Co(II). Circle color indicates Ψ increasing level (big circle: expression level changed proteins; small circle: physically interacting genes). Upregulated proteins are labeled in red, downregulated proteins are labeled in black.
